# Pooled analysis of radiation hybrids identifies loci for growth and drug action in mammalian cells

**DOI:** 10.1101/2019.12.16.877365

**Authors:** Arshad H. Khan, Andy Lin, Richard T. Wang, Joshua S. Bloom, Kenneth Lange, Desmond J. Smith

## Abstract

Genetic screens in mammalian cells commonly focus on loss-of-function approaches. To evaluate the phenotypic consequences of extra gene copies, we used bulk segregant analysis (BSA) of radiation hybrid (RH) cells. We constructed six pools of RH cells, each consisting of ~2500 independent clones, and placed the pools under selection in media with or without paclitaxel. Low pass sequencing identified 859 growth loci, 38 paclitaxel loci, 62 interaction loci and 3 loci for mitochondrial abundance at genome-wide significance. Resolution was measured as ~30 kb, close to single-gene. Divergent properties were displayed by the RH-BSA growth genes compared to those from loss-of-function screens, refuting the balance hypothesis. In addition, enhanced retention of human centromeres in the RH pools suggests a new approach to functional dissection of these chromosomal elements. Pooled analysis of RH cells showed high power and resolution and should be a useful addition to the mammalian genetic toolkit.

## Introduction

A variety of experimental strategies have been developed over the last two decades to explore genotype-phenotype correlations in mammalian cells. Gene “knockdowns” use either interfering RNA (RNAi) or small interfering RNAs (siRNAs) to specifically degrade transcripts and have given insights into multiple phenotypes (Blakely et al. 2011; Boettcher and McManus 2015; Mohr, Smith, et al. 2014; Root et al. 2006). More recently, CRISPR-Cas9 gene editing approaches that introduce targeted mutations into DNA have emerged as a powerful strategy to enable genome-wide screens for growth and pharmaceutical mechanisms (Hart et al. 2015; Koike-Yusa et al. 2014; Liu, Horlbeck, et al. 2017; Pawluk 2018; Shalem et al. 2014; Wang, Birsoy, et al. 2015).

Although these technologies have been undeniably fruitful, there are some drawbacks. Potential off-target effects exist for both RNAi (Alagia and Eritja 2016; Bofill-De Ros and Gu 2016) and CRISPR-Cas9 (Mohr, Hu, et al. 2016; Tycko et al. 2016). Furthermore, cell delivery requires the use of lentiviral libraries of unknown stability and high complexity. Assuming 20 000 protein coding genes, each knockout is represented by ~500 cells in a 75 cm^2^ flask. Decreased power, or even missing data, are therefore risks. The addition of non-coding transcripts trebles the number of genes (Derrien et al. 2012; Frankish et al. 2015; Kozomara and Griffiths-Jones 2014; Liu, Horlbeck, et al. 2017), augmenting the problem.

Overexpression can provide complementary information to loss-of-function screens (Prelich 2012). For example, screens for viability in *Saccharomyces cerevisiae* revealed no overlap of overexpressed and knockout genes, suggesting that overexpression alters cell physiology in a distinct fashion to loss-of-function (Sopko et al. 2006).

Gain-of-function screens can be particularly revealing in cancer, where overexpression is a key disease driver (Vogelstein et al. 2013). Overexpression is also valuable when genes are haploinsufficient, ruling out the use of knockouts. While cDNA overexpression can be achieved with lentiviral libraries, genome-wide screens have yet to be be performed in mammalian cells (Arnoldo et al. 2014; Yang, Boehm, et al. 2011). In addition, cDNA screens typically omit novel isoforms, whose number exceeds that of known transcripts (Eksi et al. 2013; Li, Menon, et al. 2014; Škalamera et al. 2011), as well as non-coding genes (Derrien et al. 2012; Frankish et al. 2015; Kozomara and Griffiths-Jones 2014; Liu, Horlbeck, et al. 2017).

Gene expression can be manipulated in a targeted fashion using CRISPR interference (CRISPRi) or CRISPR activation (CRISPRa), in which a catalytically dead Cas9 (dCas9) protein is fused with a transcriptional repressor or activator domain, respectively (Gilbert et al. 2014; Kampmann 2018). The approach has been used in a number of cellular screens, but the overexpression is strongly constitutive with no tissue specific regulation.

As a new strategy to dissecting genotype/phenotype relationships in mammalian cells, we have been retooling radiation hybrid (RH) technology (Ahn et al. 2009; Khan et al. 2016; Lin et al. 2010; Park et al. 2008; Ranola et al. 2010; Wang, Ahn, et al. 2011). This venerable approach to genetic mapping employs lethal doses of radiation to break the genome of a human cell line into small fragments, which are then transferred to living hamster cells by cell fusion (Cox et al. 1990; Goss and Harris 1975; Walter et al. 1994). A typical RH panel consists of ~100 clones, each containing ~4–10% of the human genome as extra copies.

Human markers that are close together tend to be found in the same set of RH clones, while markers far apart tend to be inherited independently. The number of breakpoints in RH mapping is far larger than in meiosis, giving the approach close to single gene resolution and facilitating high resolution maps of mammalian genomes (Avner et al. 2001; Hudson et al. 2001; Kwitek et al. 2004; McCarthy et al. 2000; Olivier et al. 2001).

Although RH panels were widely used in the genome projects, their potential for dissecting biological traits was not fully realized. Recently, we measured the growth of clones from a human RH panel in the presence of the antibulin drugs paclitaxel (Taxol®) and colchicine and identfied a single gene, ZNRF3, as important to the action of both agents (Khan et al. 2016). However, statistical power was limited by the modest size of the RH panel (79 clones).

Bulk segregant analysis (BSA) was developed as an efficient approach to enhancing statistical power in genetic mapping (Michelmore et al. 1991). A pool of diverse cells is placed under selection and loci that contribute to survival are revealed by enrichment of nearby genetic variants (Ehrenreich et al. 2010). The approach surveys a large population of cells using a small number of assays.

Here we describe experiments that combine RH mapping with BSA. We created six independent pools of RH cells, each with ~2500 clones. The mean length of the human DNA fragments was ~7 Mb and the number of breakpoints in each pool was ~10^5^, giving an empirical mapping resolution of ~30 kb, close to single gene. Each cell contained an average of ~4% of the human genome as extra copies, with each pool carrying the human genome with ~100× redundancy.

Unlike current screening technologies, where each cell harbors one altered gene, in RH-BSA each cell harbors multiple genes. This extra layer of multiplexing helps assure equal and robust representation of the human genome in the pooled analysis.

We placed the RH pools under growth selection in either normal medium or medium supplemented with paclitaxel. The pools were then genotyped by low pass sequencing. We correlated human copy number changes in the RH pools with cell survival, identifying over 800 growth loci, nearly 40 paclitaxel loci, over 60 interaction loci and 3 loci that controlled mitochondrial copy number. Both coding and non-coding genes were detected and the growth genes possessed complementary properties to those identified in loss-of-function screens. The RH-BSA strategy appears to be a useful complement to existing genetic strategies in mammalian cells, revealing new dimensions in the relationship between genotype and phenotype.

## Results

### Creating six RH pools

The strategy for RH-BSA is shown in **Figure 1**. Lethally irradiated human HEK293 cells were fused to living hamster A23 cells and plated in selective medium. The surviving RH cells receive the selectable marker, thymidine kinase (TK1), from the donor cells in addition to a random assortment of human DNA fragments. We maintained the RH cells as heterogeneous pools, rather than picking individual clones as is usually done for genetic mapping. The pools were then genotyped by low pass sequencing before and after competition due to growth or drug exposure. Human genes that confer a proliferative advantage become more abundant, while genes that confer a disadvantage become less abundant.

**Figure 1.**
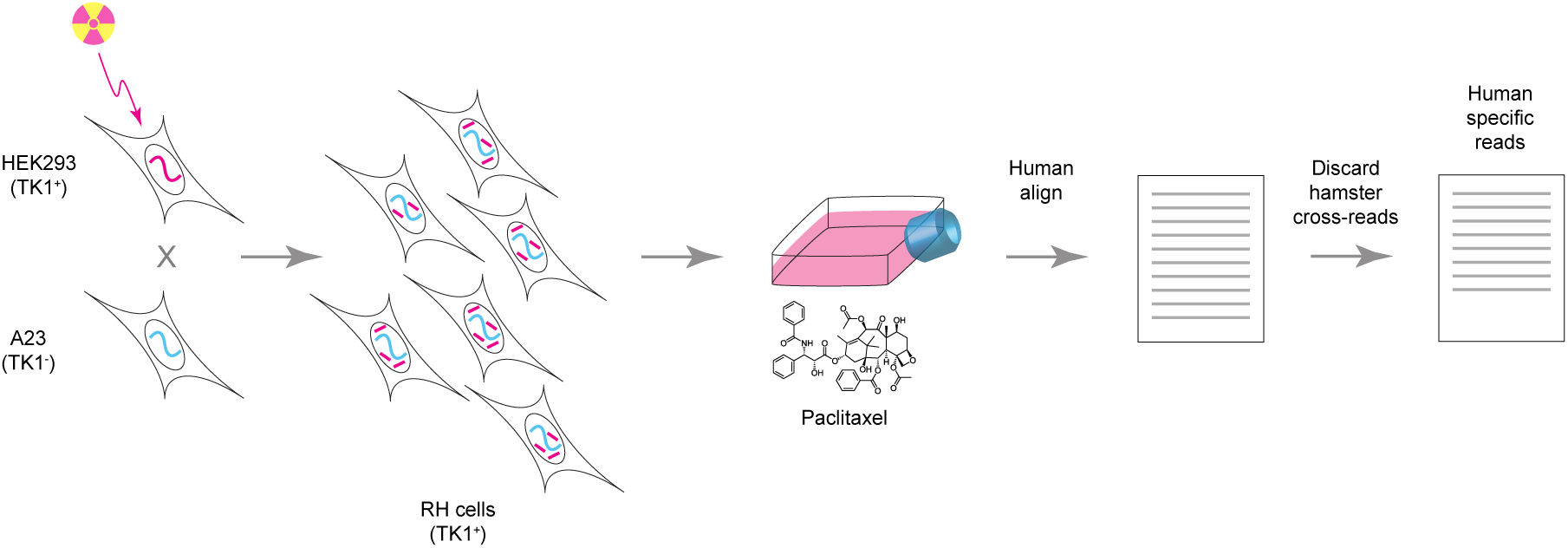
Creating the RH pools. Human HEK293 cells (TK1^+^) were lethally irradiated, fragmenting their DNA, and fused with living hamster A23 cells (TK1^-^). RH cells, containing TK1 and a random sample of DNA fragments from the human donor cells, were selected in HAT medium. Six independent pools were grown for various times in the HAT medium supplemented with differing concentrations of paclitaxel. (One pool is shown here.) The pools were genotyped by low pass sequencing. Reads aligning to the human genome were purged of cross-species reads from hamster before further analysis and vice versa.

We created six independent RH pools, each with a mean fragment size of 7.4±0.1 Mb (**Figure S1, Supporting Information**). Each pool had 2621±441 clones, after taking into account a spontaneous revertant rate of 26%±2% (**Table S1**).

When the cells from the fusion reactions reached confluence, an aliquot from each of the pools was reserved for genotyping. These samples represent week 0 and 0 nM paclitaxel (**Table S2**) and are referred to as the “RH pools”. The cells were further propagated as separate pools and genotyped after 1, 2, 3, 4 and 6 weeks in HAT medium supplemented with 0, 8, 25 or 75 nM paclitaxel. There were 115 RH samples in total.

### Human DNA in the RH samples

We used low pass sequencing to analyze the RH samples under the various conditions of growth and drug (**Figure 1, Tables S3–S6, Supporting Information**). A mean of 40.5±0.7 million (M) reads were obtained for each sample, corresponding to a sequencing depth of 0.84±0.01 times the human genome. Reads were aligned to the human and hamster genomes and cross-species reads discarded, leaving only species-specific alignments for subsequent analyses.

The hamster genome in the six RH pools was not appreciably changed by the cell fusions (**Figures 2A, 2B, S2 and S3A, Supporting Information**). In contrast, a selection of human DNA fragments from the HEK293 cells were retained by each RH pool (**Figures 2C–2F, S3B, S4–S6**).

**Figure 2.**
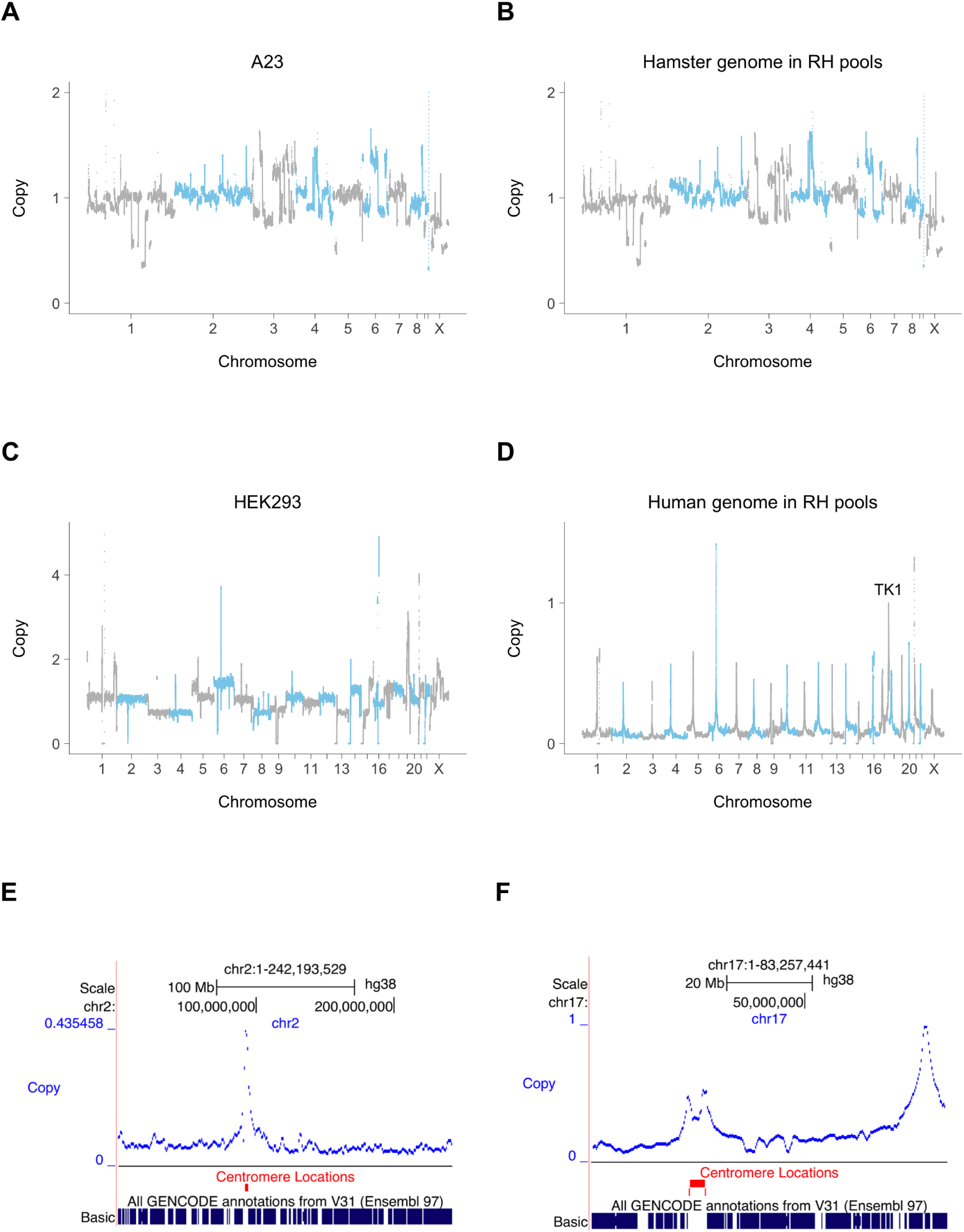
Genome copy number in the RH pools. (**A**) Hamster copy number in the A23 cells, normalized to a mean value of 1. (**B**) Hamster copy number averaged across the six RH pools. (**C**) Human copy number in the HEK293 cells. (**D**) Human DNA retention averaged across the six RH pools, TK1 assigned retention of 1. (**E**) Increased retention of human centromere on chromosome 2. (**F**) Increased retention of human centromere on chromosome 17 and TK1 near 17q.

Since human TK1 was used as the selectable marker, this gene has an expected retention frequency of 1 in the hybrid cells. Donor centromeres also showed increased retention, as these chromosomal regions confer increased stability on the DNA fragments (**Figures 2D–2F, S5 and S6, Supporting Information**) (Wang, Ahn, et al. 2011). In contrast, non-centromeric regions showed relatively constant retention across the genome, as previously documented in RH panels (Avner et al. 2001; Hudson et al. 2001; Kwitek et al. 2004; McCarthy et al. 2000; Olivier et al. 2001) (**Figures 2D–2F, S5 and S6**).

The mean retention frequency in the RH pools was 3.9%±0.3% (**Table S7, Supporting Information**), with each gene harbored by 3.9 × 10^5^ (±3.4 × 10^4^) cells in a 75 cm^2^ flask of 10^7^ RH cells. The human genome is represented with 103±20-fold redundancy in each of the six individual pools.

The mapping resolution of RH-BSA depends on the length of the DNA fragments, their average retention frequency and the number of clones. At the mean fragment size of 7.4 ± 0.1 Mb, the number of breakpoints in the individual RH pools is 8.7 × 10^4^ (± 1.7 × 10^4^) and is expected to yield a mapping resolution of between 71 *±* 14 kb for each pool and 12 ± 3 kb for all six pools together.

### Copy number changes in the RH samples

Human DNA is commonly lost during growth of hybrid cells, and this phenomenon has been exploited as a tool for gene mapping (Harris 1993; Kucherlapati and Ruddle 1975; Wasmuth 2001). In a pooled setting, alterations in human DNA levels can also occur as a result of changes in relative clone abundance.

The overall mean retention of human DNA in the RH samples showed a significant decrease as a result of growth (3.9% ± 0.3%, week 0; 1.1% ± 0.2%, week 6), but not paclitaxel (**Figures S7–S9, Supporting Information**). When profiled across the genome, the hamster genome was stable, while human DNA levels showed significant decreases together with increased variance (**Figures 3 and S10, Tables S8 and S9, Supporting Information**).

**Figure 3.**
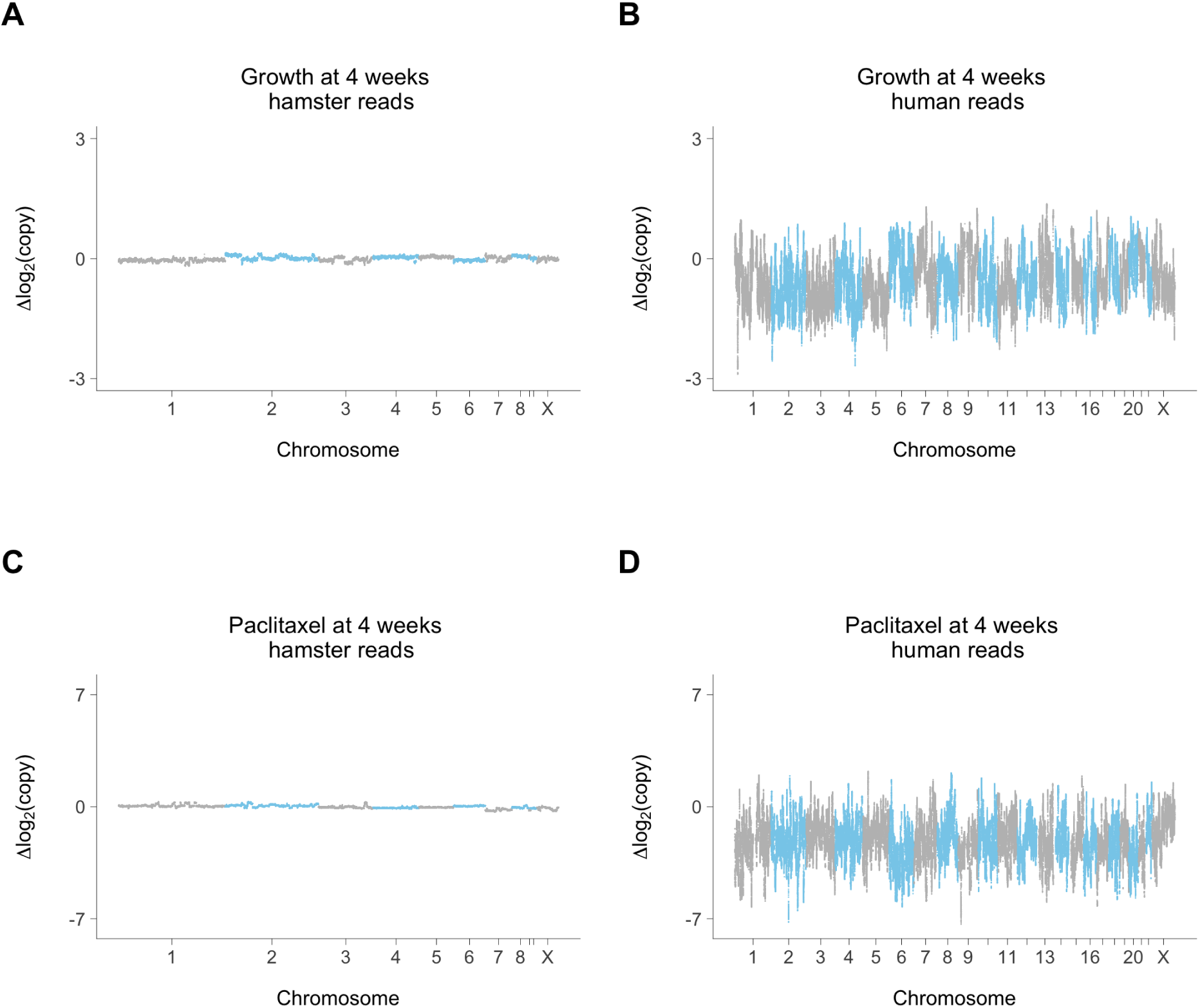
Copy number changes at week 4. (**A**) Hamster genome, growth at week 4 compared to week 0; 0 nM paclitaxel. (**B**) Human genome, growth at week 4 compared to week 0; 0 nM paclitaxel. (**C**) Hamster genome, 75 nM paclitaxel compared to 0 nM paclitaxel; week 4. (**D**) Human genome, 75 nM paclitaxel compared to 0 nM paclitaxel; week 4. Hamster and human copy number changes on same log_2_ scale, representing change in relative copy number normalized to hamster genome averaged across the six RH pools.

### Growth loci

#### Growth loci tally

Human growth loci were identified by detecting significant changes in copy number across the genome as a result of growth time at the various paclitaxel concentrations (**Figures S11–S17, Materials and Methods, Supporting Information**). Cross-validation procedures confirmed high levels of reproducibility (**Figure S18, Supporting Information**). In total, there were 859 unique growth loci, ∼1.4% of all known genes (**Figures 4 and S19–S21, Tables S10–S16, Supporting Information**).

**Figure 4.**
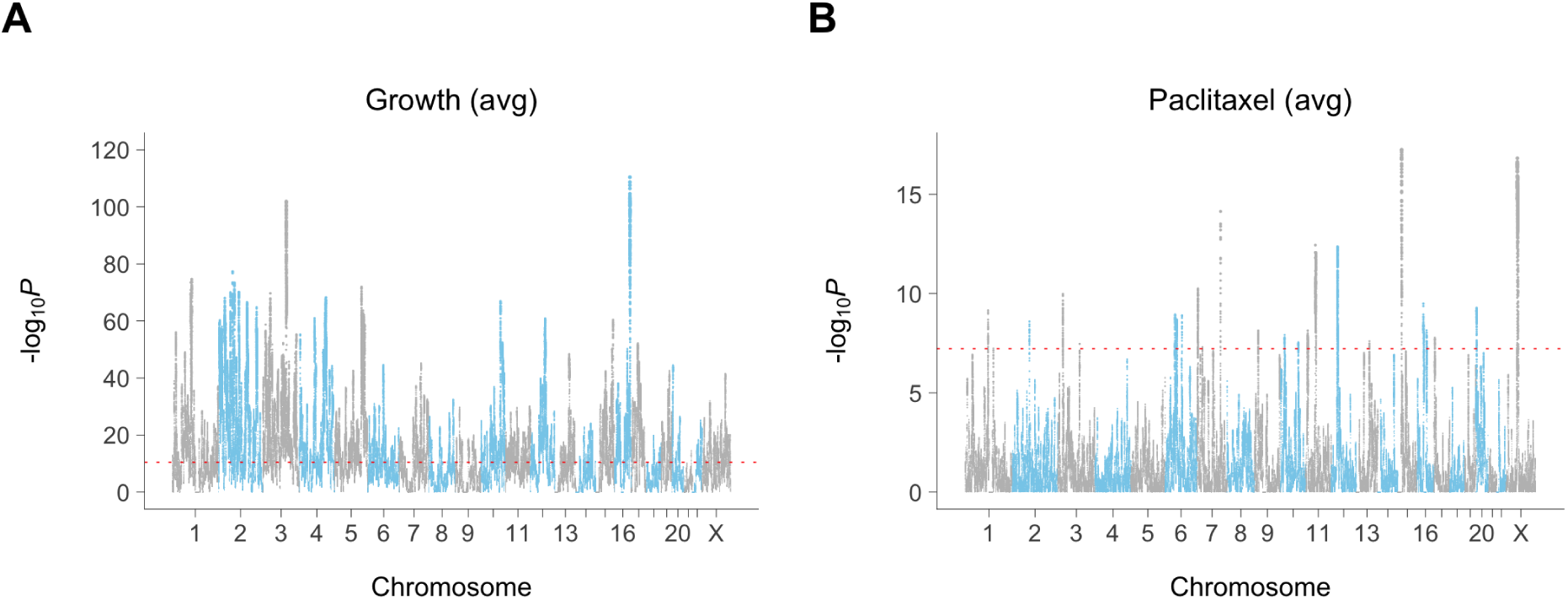
Loci for average conditional effects of growth and paclitaxel. (**A**) Significance values for growth. (**B**) Significance values for paclitaxel. Red dotted line, permutation significance threshold. clearly play a substantial role in cell proliferation. Close-up views and copy number changes for growth loci are shown in **Figures 5** and **S22** (**Supporting Information**).

#### Coding and non-coding growth loci

Four of the growth loci mapped to centromeres (**Supporting Information**). Of the remaining loci, the nearest genes were coding for 442 (52%) and non-coding for 413 (48%). Although the RH coding growth genes were significantly enriched compared to non-coding (**Supporting Information**), non-coding genes

#### Nearly all growth loci show DNA loss

Consistent with the loss of human DNA in the RH samples (**Figures 3, S7–S10**) only seven of the 859 unique growth loci, including SEMA3A and TOR1A, had positive coefficients (**Figure S23A**). The predominance of gene loss is consistent with the frequent loss of human DNA in RH cells and echoes the finding that inactivation of tumor suppressor genes is more common in cancer than oncogene activation (Harris 1993; Vogelstein et al. 2013).

#### RH growth genes and complex human disease

The RH growth genes were linked with common human disease. The genes were significantly enriched in the catalog of human genome-wide association studies (GWAS) (false discovery rate, FDR = 5.8 × 10^−12^) (Chen et al. 2013) and also significantly over-represented in 16 of 203 disease-related terms in the Database of Genotypes and Phenotypes (dbGaP) (FDR *<* 0.05) (Chen et al. 2013; Tryka et al. 2014). In contrast, there was no significant enrichment of essential loss-of-function genes identified from CRISPR screens in common human disease (FDR *>* 0.76, GWAS; FDR *>* 0.25, dbGaP) (Hart et al. 2015; Wang, Birsoy, et al. 2015).

Most common human disease variants represent polymorphisms in regulatory regions, which have more subtle effects than null alleles (Gallagher and Chen-Plotkin 2018). The enrichment of RH growth genes in human disease may thus be related to the restrained effects of an extra copy compared to a knockout. The RH growth genes may be good candidates for causative genes in complex human disease.

#### The balance hypothesis

The “balance” hypothesis postulates that the effects of altered gene dosage occur as a result of unequal synthesis of gene products (Papp et al. 2003). The hypothesis yields two predictions: (**1**) genes that show a phenotype when dosage is increased should also show a phenotype when dosage is decreased, (**2**) genes with dosage effects should be more likely to participate in multisubunit complexes. The balance hypothesis has been falsified in yeast (Sopko et al. 2006), but whether it is true in mammalian cells is unknown.

We tested the first prediction by asking whether there was increased overlap between the growth genes identified using RH-BSA and those identified in loss-of-function CRISPR (Hart et al. 2015; Wang, Birsoy, et al. 2015) and CRISPRi screens (Gilbert et al. 2014). Contrary to expectations, the RH growth genes had significantly decreased effect scores, decreased cell line hit rates and significant under-representation among the CRISPR growth genes (**Figures 6A–6F, S24 and S25**). These observations refute the first prediction of the balance hypothesis.

**Figure 5.**
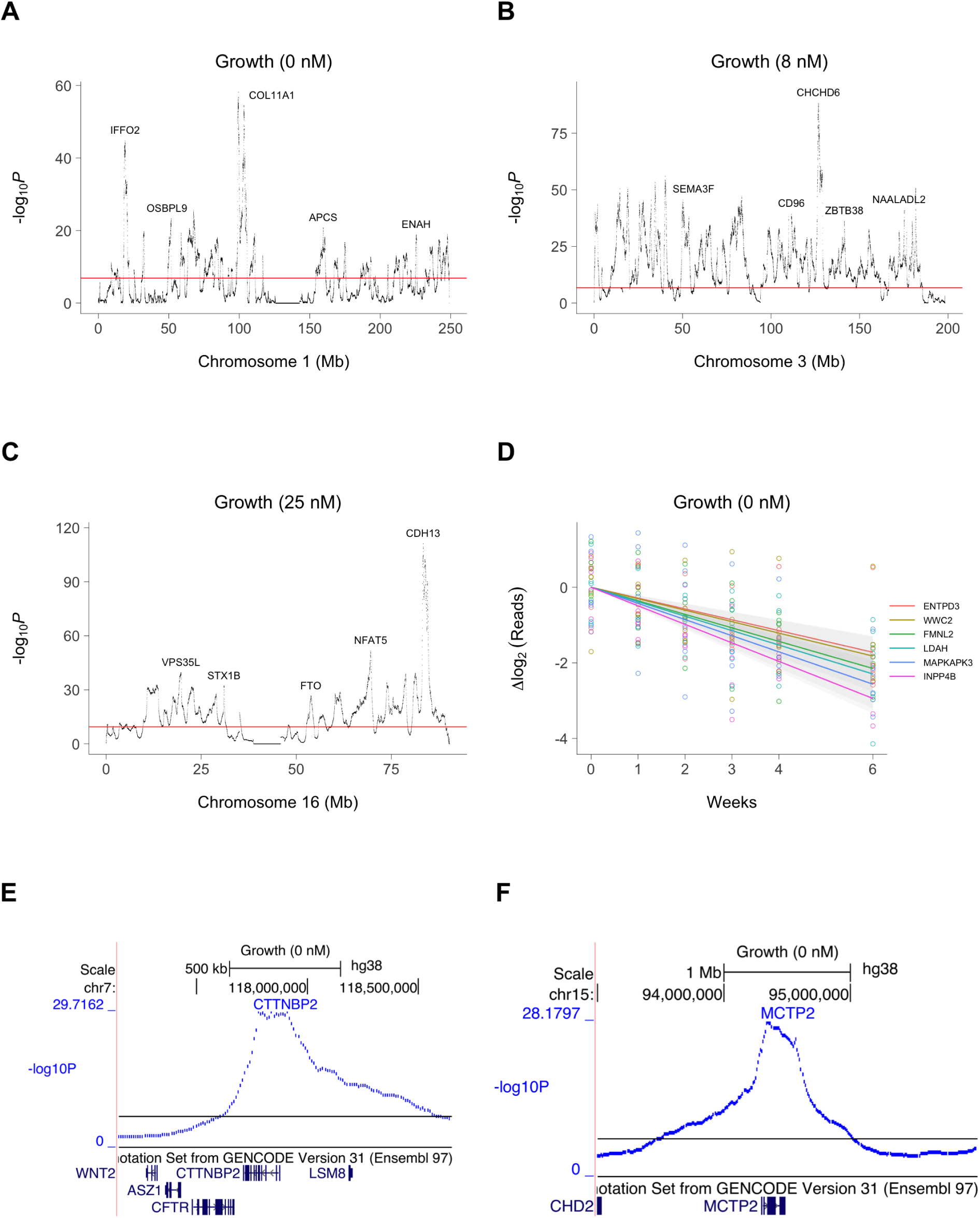
Close up views of growth loci. (**A**) Chromosome 1. (**B**) Chromosome 3. (**C**) Chromosome 16. (**D**) Normalized sequence reads on log_2_ scale for six significant growth loci. Colored lines, best fit. Grey bands, 95% confidence intervals. (**E**) Locus for CTTNBP2 on chromosome 7. (**F**) Locus for MCTP2 on chromosome 15. Horizontal red and black lines, permutation significance thresholds.

**Figure 6.**
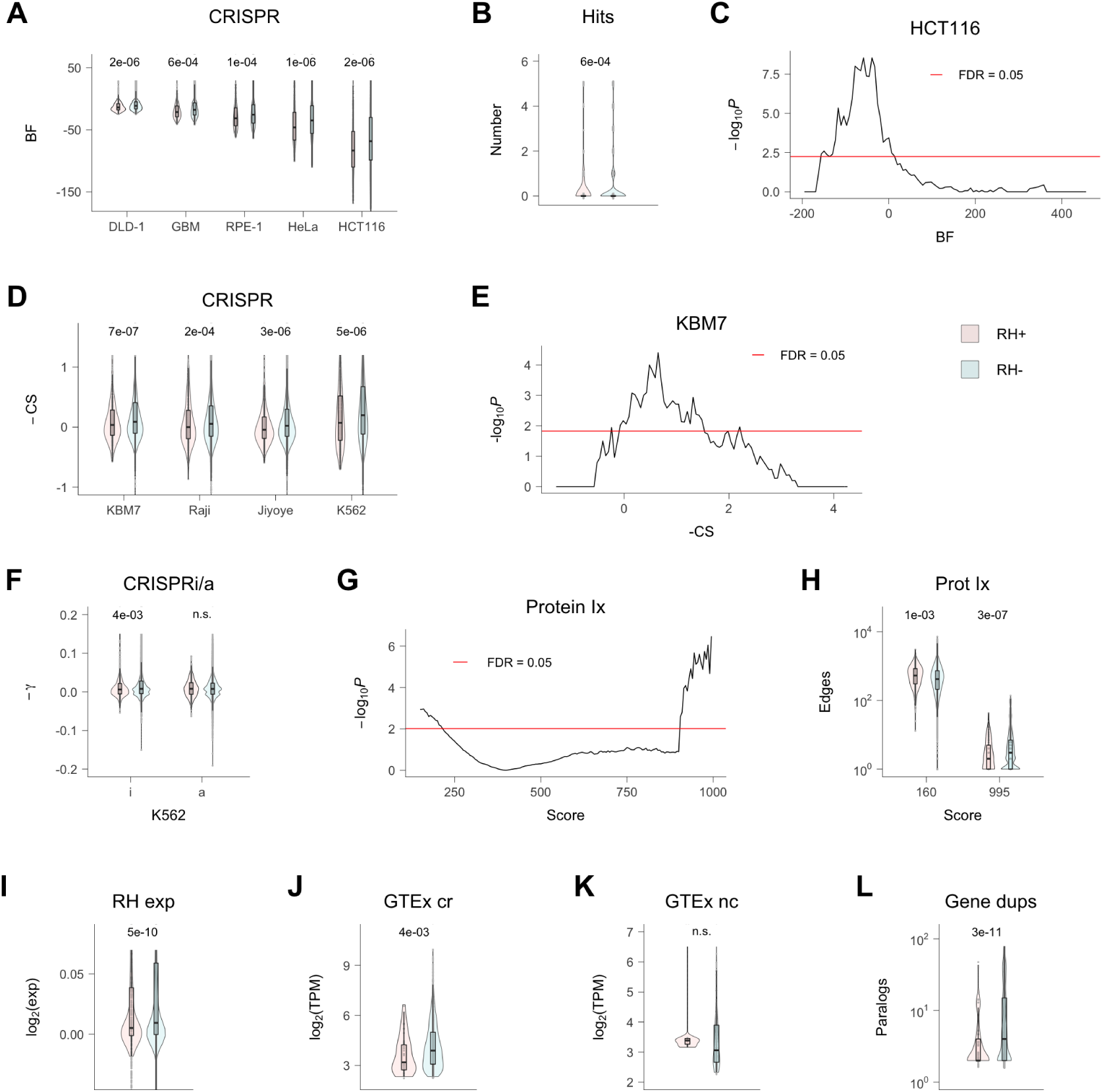
RH growth genes. (**A**) RH growth genes (RH+) have weaker CRISPR effects (Hart et al. 2015) than non-growth RH genes (RH-). BF, Bayes factor for growth effects in each cell line; *P* values shown above comparisons. (**B**) Mean number of cell lines with a CRISPR hit is lower for RH growth genes. (**C**) Lack of overlap between RH and CRISPR growth genes in HCT116 cells. Abscissa shows threshold BF used to calculate overlap. (**D**) CRISPR scores (CS) for growth effects (Wang, Birsoy, et al. 2015) multiplied by −1, so that higher scores mean stronger effects. (**E**) Lack of overlap between RH and CRISPR growth genes in KBM7 cells. (**F**) RH growth genes show weaker CRISPRi, but not CRISPRa, effects (−*γ*) in K562 cells (Gilbert et al. 2014). (**G**) RH growth genes have increased numbers of low confidence protein-protein interactions and decreased numbers of high confidence interactions. Score threshold (abscissa) is a measure of confidence. (**H**) Number of protein-protein interactions at most significant low confidence score threshold (160) and most significant high confidence threshold (995). (**I**) RH coding region growth genes have decreased expression compared to non-growth genes in microarray data from a human RH panel. (**J**) Coding region (cr) RH growth genes have decreased expression compared to cr non-growth genes in GTEx RNA-Seq data from substantia nigra. (**K**) Non-coding (nc) RH growth and non-growth genes show no significant expression differences in same tissue. (**L**) Decreased number of human paralogs (duplicates) for RH growth genes. *P* values, Welch Two Sample t-test except (C) and (E), Fisher’s exact test. FDR, false discovery rate (Benjamini and Hochberg 1995). Transcripts per million (TPM) thresholded at *≥* 5.

To test the second prediction we asked whether the RH growth genes had increased likelihood of protein-protein interactions, similar to the CRISPR essential genes (Hart et al. 2015; Wang, Birsoy, et al. 2015). Using the STRING v11 database (Szklarczyk et al. 2019), the RH growth genes showed decreased numbers of high confidence protein-protein interactions, and a corresponding increase in low confidence interactions (**Figures 6G and 6H**). Further, the RH growth genes showed no significant enrichment in protein interactions, molecular pathways or multisubunit protein complexes using Enrichr or DAVID (FDR *>* 0.05) (**Supporting Information**) (Chen et al. 2013; Huang et al. 2009). These findings contradict the second prediction of the balance hypothesis.

Based on our results and those in yeast (Sopko et al. 2006), it appears that gene dosage alterations do not exert their effects through unbalanced synthesis of gene products, even in evolutionarily distant cells. Other mechanisms are likely responsible for gene dosage effects.

#### Further divergent properties of RH-BSA and loss-of-function growth genes

As well as the divergent properties of the RH and CRISPR growth genes discussed in the previous section (**The balance hypothesis**), there were additional discrepancies. Essential coding region genes identified in loss-of-function CRISPR screens showed increased expression (Hart et al. 2015; Wang, Birsoy, et al. 2015). In contrast, the RH-BSA growth genes had the opposite behavior with decreased expression of coding, but not non-coding, genes in both a microarray dataset from a human RH panel (Wang, Ahn, et al. 2011) (**Figure 6I**) and in RNA-Seq data from human tissues (GTEx Consortium 2015) (**Figures 6J and 6K, S26**). The decreased expression of the RH growth genes may be the result of evolutionary selection for this property in genes with high dosage sensitivity.

CRISPR growth genes were less likely to have human paralogs because duplicated genes offer functional redundancy (Wang, Birsoy, et al. 2015). The RH growth genes also had fewer duplicates, but perhaps for a different reason; additional copies of the RH growth genes could result in adverse phenotypes as these genes inhibit proliferation when present in an extra dose (**Figure 6L**).

#### Similar properties of RH-BSA and loss-of-function growth genes

In other respects, the RH and CRISPR growth genes had comparable properties. Both sets of genes were intolerant of loss-of-function mutations, suggesting similar functional importance (**Figure S27A**) (Wang, Birsoy, et al. 2015). In addition, both the RH and CRISPR growth genes showed decreased human/mouse sequence divergence indicating evolutionarily conserved roles (**Figure S27B**) (Hart et al. 2015; Wang, Birsoy, et al. 2015). The RH and CRISPR growth genes also showed increased numbers of orthologs across species (“phyletic retention”), further suggesting evolutionary conservation of these two gene classes (**Figure S27C**) (Wang, Birsoy, et al. 2015).

The CRISPR screens for essential genes were restricted to coding genes and showed longer transcripts, increased exon numbers and longer coding sequences (Hart et al. 2015; Wang, Birsoy, et al. 2015). The coding and non-coding RH growth genes also showed increased gene length, transcript size and exon numbers, as well as longer open reading frames for coding genes (**Figures S27D–S27J**).

Receiver operating characteristic (ROC) curves showed that gene length was the best predictor of RH growth genes (area under the curves or AUC = 0.76 and 0.70 for coding and non-coding genes, respectively), followed by CRISPR essentiality scores, protein interactions and RH expression levels (**Figure S27K**). The ROCs were significantly better predictors than random (*P <* 4.9 × 10^−4^, Wilcoxon rank sum test).

Surprisingly, the RH growth genes showed no significant increase in effect scores, or overlap, with growth genes from a CRISPRa screen (**Figures 6F** and S25F) (Gilbert et al. 2014). The constitutive overexpression of the CRISPRa technology may be responsible for the discrepancy.

#### Novel RH growth genes

The coding region RH growth genes were relatively poorly studied, having fewer literature citations in the GeneRIF and Reactome databases (**Figure S28, Supporting Information**). In addition, of the 442 coding region RH growth genes, 148 had neither literature citations in GeneRIF nor entries in the Reactome database. Study of these poorly understood genes may provide insights into mammalian cell growth.

Unconventional RH growth loci included six that mapped to olfactory gene clusters (**Figure S29A, Supporting Information**) and two that mapped to gene deserts, with the nearest gene *>* 250 kb away (**Figures S29B and S29C, Supporting Information**).

### Paclitaxel loci

Paclitaxel loci were identified by detecting significant copy number changes as a result of rising drug concentration at the various growth times. The number of loci increased significantly with longer growth periods, suggesting that the various exposure times recruit different genes (**Figures 4B, S20, S21B and S30, Tables S10, S11, S14, S15 and S17–S19, Supporting Information**). This observation is consistent with our previous findings using a panel of human RH clones (Khan et al. 2016), as well as findings in yeast (Wang and Kruglyak 2014). There were 38 unique paclitaxel loci in total.

Compared to the growth loci, a significantly greater proportion of unique paclitaxel loci (7 out of 38) had positive coefficients indicating increased copy number as a result of drug exposure (**Figure S23B, Supporting Information**).

Of the 34 non-centromeric unique paclitaxel loci, the nearest genes were coding for 19 (56%) and non-coding for 15 (44%), not significantly different from random (**Supporting Information**). Coding and non-coding genes make important contributions to the action of paclitaxel as well as growth.

Close-up views and copy number changes for significant paclitaxel loci are shown in **Figure 7**. Consistent with the activity of paclitaxel as an antitubulin drug, there were two genes among the paclitaxel loci known to play a role in microtubule function; NEK10 (Porpora et al. 2018; Yi et al. 2018) and PDE4DIP (Bouguenina et al. 2017; Yang, Wu, et al. 2017). The role of PDE4DIP may explain the clinical utility of phosphodiesterase 4 inhibitors as antiproliferative and immunosuppressive agents (Keating 2017). There were 10 genes in common between the 38 unique paclitaxel loci and 859 unique growth loci, including LSAMP, SEMA3D and GATAD2A (**Figure S31, Supporting Information**). The dual role of these genes in growth and microtubule function suggests they may be attractive targets for the development of antiproliferative drugs.

**Figure 7.**
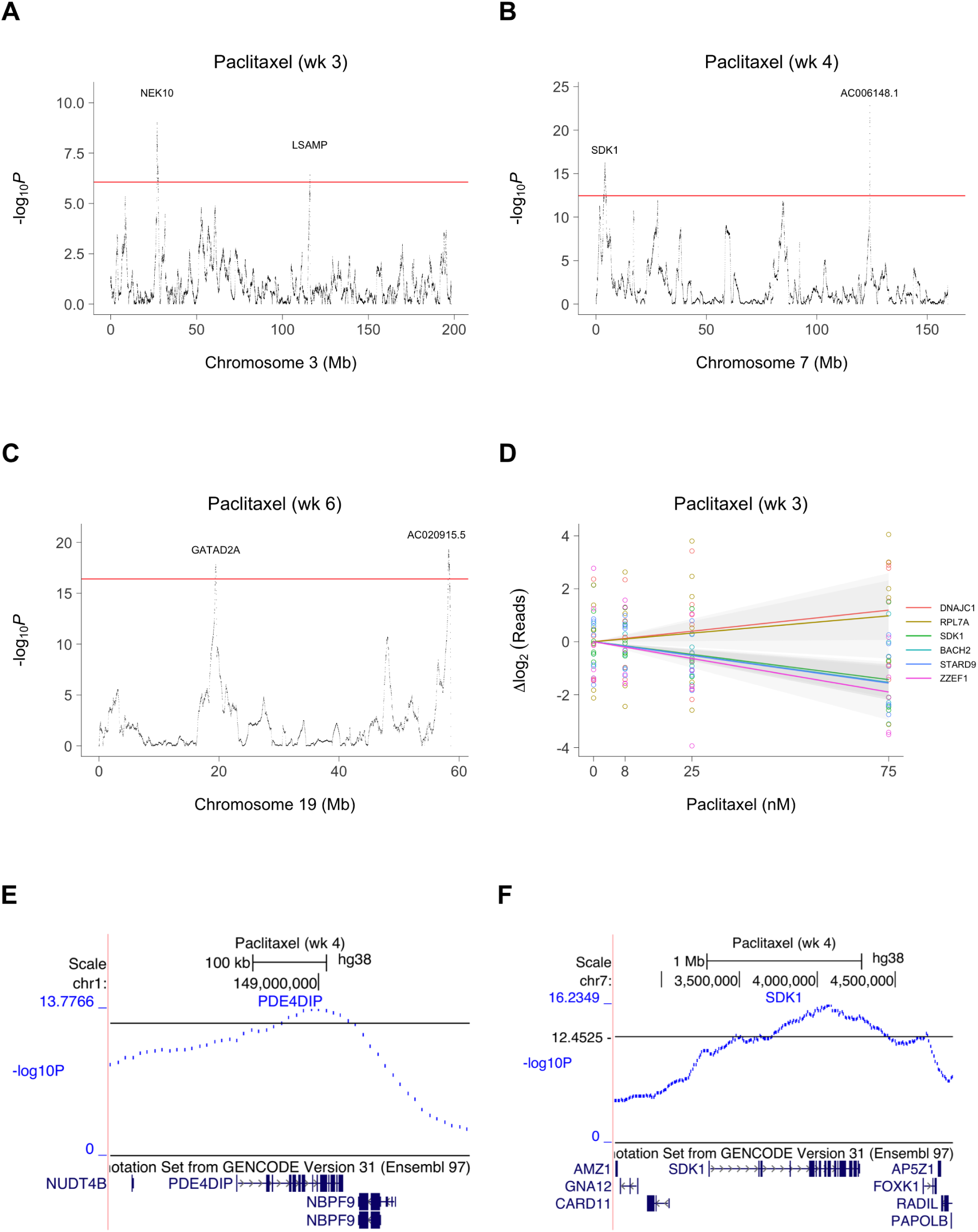
Close up views of paclitaxel loci. (**A**) Chromosome 3. (**B**) Chromosome 7. Chromosome 19. (**D**) Normalized sequence reads on log_2_ scale for six paclitaxel loci. Colored lines, best fit. Grey bands, 95% confidence intervals. (**E**) Locus for PDE4DIP on chromosome 1. (**F**) Locus for SDK1 on chromosome 7. Horizontal red and black lines, permutation significance thresholds.

A number of the paclitaxel loci mapped to novel genes, suggesting fresh entry points for understanding the mechanisms of action of this drug (**Supporting Information**).

### Interaction loci

A total of 62 interaction loci were identified based on human copy number changes due to growth depending significantly on paclitaxel concentration and vice versa. Of the 57 non-centromeric interaction loci, the nearest genes were coding for 26 (46%) and non-coding for 31 (54%), not significantly different from random (**Supporting Information**). Coding and non-coding genes make important contributions to the action of the interaction genes.

Of the interaction loci, 15 overlapped with the 859 unique growth loci and 14 overlapped with the 38 unique paclitaxel loci (**Figures S21C, S23C and S31–S33, Tables S10, S11, S14, S15 and S20, Supporting Information**). The interaction loci included genes previously demonstrated to co-regulate growth and tubulin function. For example, HGF promotes microtubule assembly and shows complex cell growth effects when combined with paclitaxel (Dugina et al. 1995; Rasola et al. 2004; Tian et al. 2014; Ying et al. 2015). Similarly, FSIP1 regulates growth of triple-negative breast cancer cells and also resistance to docetaxel, a drug in the same class as paclitaxel (Liu, Sun, et al. 2018). SIKE1 (Sonnenschein et al. 2018) and CPT1A (Li, Zhao, et al. 2013) also regulated both cell growth and microtubule activity. The interaction genes may thus provide useful clues to the relationship between growth inhibition and paclitaxel action (**Supporting Information**).

### Empirical evaluation of mapping resolution using RH-BSA

The −2log_10_ *P* width of the RH loci (**Figures 5** and **7**) suggested a mapping resolution of *<*100 kb. To obtain an empirical estimate of mapping accuracy, we used the increased retention frequency of TK1 and the centromeres as a proxy for resolution (**Figures 2, S5, S6 and S34, Supporting Information**). The mapping accuracy estimated using both approaches was similar, together suggesting a resolution ~30 kb.

Despite its high mapping resolution, the RH strategy could not always resolve adjacent loci, particularly in areas of high gene density (**Figure S35, Supporting Information**). In the future, increased radiation doses and pool numbers will help resolve these complex regions.

### Mitochondrial copy number

#### Mitochondria in HEK293 and A23 cells

The sequence data revealed a mitochondrial copy number of 2145 for the donor HEK293 cells and 843 for the recipient A23 cells (**Table S4, Supporting Information**), similar to published values for mammalian mitochondria (van Gisbergen et al. 2015; Phillips et al. 2014; Reznik et al. 2016; Wachsmuth et al. 2016).

#### Human loci regulating hamster mitochondrial copy number

The hamster mitochondrial copy number in the six RH pools was 757±63, not significantly different from A23 cells (t[1,5] = 0.51, *P* = 0.63; Two Sample t-test) (**Figures S36A and S36B, Table S21, Supporting Information**).

We identified three human loci that significantly regulated hamster mitochondrial copy number in the RH samples, DARS2, GORAB and GRID2 (**Figure S37, Tables S22 and S23, Supporting Information**). Increased dosage of all three loci was associated with higher hamster mitochondrial copy number (**Figures S37A–S37D**). DARS2 encodes human mitochondrial aspartyl-tRNA synthetase (Scheper et al. 2007), while GRID2 encodes the glutamate ionotropic receptor delta type subunit 2 which has previously been shown to promote mitochondrial fission (Liu and Shio 2008).

#### Human loci regulating human mitochondrial copy number

Human mitochondria were only present at low copy number (1.0±0.4, mean copy number all RH samples; t[1,5.3] = 2.7*, P* = 0.04, Kenward-Roger degrees of freedom; average conditional effect), suggesting that these organelles are either donated inefficiently or replicate poorly in hamster cells (**Figures S36C–S36H, Table S21, Supporting Information**).

Consistent with the low levels of human mitochondria, we found no permutation significant human loci that regulated their copy number (**Tables S24 and S25**). However, one locus on chromosome 3 surpassed the less stringent FDR threshold of 0.05 (−log_10_ *P* = 8.6, FDR = 8.1 × 10^−4^, **Figures S37E and S37F**). This locus mapped to CAPN7 and was 14.9 kb from SH3BP5, whose product is localized to the outer mitochondrial membrane and is involved in regulation of apoptosis (Win et al. 2018). Nevertheless, this locus should be viewed with considerable caution; deeper sequencing may provide stronger evidence for loci that regulate human mitochondrial copy number.

### Hamster loci

Significant hamster loci due to pre-existing copy number alterations (CNAs) (**Figures 2A, 2B and S2**) were identified for growth, paclitaxel and their interaction (**Figures S38 and S39, Tables S26 and S27, Materials and Methods, Supporting Information**).

The hamster loci had weaker −log_10_ *P* values than the human, consistent with the greater stability of the hamster genome (**Figures 3 and S10, Supporting Information**). In addition, hamster CNAs and their corresponding loci ranged over Mb which, combined with random fluctuations in −log_10_ *P* values, precluded gene identification (**Figure S40**).

## Discussion

Our rationale for developing RH-BSA was three-fold. (**1**) To advance a genome-wide method that employed extra gene dosage to evaluate gene function and which was complementary to loss-of-function approaches. (**2**) Develop RH-BSA as a method to assess coding and non-coding genes on an equal footing. (**3**) Improve the statistical power of the RH approach, while enhancing its already impressive mapping resolution.

We created six indpendent pools of RH cells, each consisting of ~2500 RH clones. Low-pass sequencing was used to identify human genes with significant changes in copy number as a result of growth or paclitaxel. We used cross-validation to demonstrate the high reliability of the approach.

Significant loci were identified using permutation, with a null expectation for each genome-scan of 0.05 false positives, or 0.7 loci from 13 genome scans. We found 859 unique growth loci, 38 paclitaxel loci, 62 interaction loci and 3 loci regulating mitochondrial copy number, far exceeding expectations and suggesting a low false positive rate.

We used the increased retention of TK1 and the centromeres to estimate the mapping resolution of RH-BSA as ~30 kb. Unlike association mapping (Edwards et al. 2013), the resolution of RH-BSA is not limited by recombination hotspots and linkage disequilibrium. In fact, the resolution and power of RH-BSA can be further increased by using higher doses of radiation and more pools. Ten-fold enhancements are readily attainable.

While the measured resolution was sufficient for single gene identification, gene rich regions in particular could not be clearly resolved. Nevertheless, a number of conclusions could be drawn from our study.

The RH-BSA growth genes showed significant non-overlap with genes identified from loss-of-function screens, as well as decreased participation in multi-subunit complexes. These findings repudiated the balance hypothesis, agreeing with findings from yeast. In addition, the RH-BSA growth genes had lower expression levels than average, consistent with the genes exerting adverse phenotypic effects when present in an extra copy.

In addition to their divergent properties, the RH-BSA and loss-of-function growth genes also shared some attributes. In particular, both sets of genes showed higher evolutionary conservation and increased gene lengths.

The RH growth genes were over-represented in the GWAS catalog and dbGaP. Complex trait variants are often found in regulatory regions, exerting their effects through altered gene expression. The extra human gene copies in the RH pools may simulate these mild allelic effects and hence represent attractive candidate genes for complex traits. In addition, both coding and non-coding RH genes made substantial contributions to cell growth, paclitaxel action and their interaction.

Three significant human loci that regulated hamster mitochondrial copy number were identified, two of which had known functions in mitochondria. Further study of these genes may provide useful therapeutic insights into mitochondrial disorders. RH-BSA may also allow better mapping of the functional sequences of centromeres based on increased retention.

Changes in human DNA levels in the RH pools could be due either to DNA loss or altered clone abundance. Pooled analysis cannot distinguish between these possibilities. However, the majority of the growth and paclitaxel loci showed decreased copy number, which is consistent with the known tendency of RH cells to lose human DNA fragments. Thus, DNA loss may be the predominant mechanism of copy number change in the RH pools. Single cell analysis can discriminate unequivocally between the two alternatives.

RH cells are a unique tool. These cells are not entirely normal. Their genome does not exist in nature, representing hybrids of two closely related mammalian species, human and hamster. They are also aneuploid, with ~4% of the genome in the triploid state. Nevertheless, the cells grow and divide predictably, and contain all the standard features found in mammalian cells (eg cytoskeleton, mitosis, etc.). For a small number of genes, the evaluated phenotypes may be due to species differences, but such occurrences will be infrequent and will still be valuable in highlighting phenotypic mechanisms.

Other cell types and species, including human, can be employed as recipients in the cell fusion process, widening the variety of tissue types beyond fibroblasts (Hiratsuka et al. 2015; Liskovykh et al. 2016; Suzuki et al. 2016). In addition, selectable markers such as neomycin resistance can be used instead of TK1, alleviating concerns about the perturbation of nucleotide synthesis pathways by HAT medium (Aoki et al. 2014).

We previously identified ZNRF3, a regulator of Wnt signaling, as a target for paclitaxel action using individual clones from the human G3 RH panel (Khan et al. 2016). This locus was not replicated using the RH pools. In our original study, the RH clones were weaned from HAT, unlike the RH-BSA experiments. In addition, the ability to detect growth factor signaling pathways is likely to be attenuated in a pooled setting. These differences may explain the lack of replication.

The RH-BSA approach can be harnessed to uncover genes for any phenotype where a convenient selection can be devised (Miyake et al. 2019). The approach is complementary to loss-of-function screens and provides information on both coding and non-coding genes. RH-BSA should be a valuable addition to the suite of tools for mammalian genetics.

## Materials and Methods

### Sequencing

Libraries were prepared using the Illumina TruSeq Nano DNA library prep kit employing 2 *µ*g of DNA from each RH sample. We performed the sequencing using 1 × 64 bp single reads on an Illumina Hiseq 2500 machine in rapid run mode with OLB (off-line basecaller) software and on-board cluster generation.

Reads were aligned to the indexed GRCh38/hg38 human genome assembly (hg38.fa) or the Chinese hamster (*Cricetulus griseus*) genome assembly (RAZU01) (Rupp et al. 2018) using bwa-0.7.12 mapping software (Li and Durbin 2009). The reference genomes were downloaded from the UCSC genome browser prior to use (https://genome.ucsc.edu) (Kent et al. 2002), unzipped and relevant files concatenated and indexed using BWA software. Sequence reads were aligned to the human and hamster genomes at high stringency, allowing only one mismatch. Aligned reads were then cross-aligned to the other species and reads that aligned to both species discarded, effectively discriminating between human or hamster sequences (**Results**).

Aligned sequences were converted to SAM format and sorted and indexed using samtools-1.2 software (Li, Handsaker, et al. 2009). To read and view sequence information, SAM files were converted to BAM files employing SAMtools. Indexed sorted BAM files were converted to BED files using BEDTools-2.15.0 software (Quinlan and Hall 2010).

### Read quantitation

The human and hamster nuclear reference genomes were indexed into 1 Mb windows using BEDTools, with 10 kb steps for both human and for hamster. BED files were then mapped against the indexed reference genome and the number of reads per 1 Mb window obtained. We chose the 1 Mb window size as being small enough to reflect the mapping accuracy of RH-BSA, but large enough to keep the sampling variance within bounds. The 10 kb step size was small enough to accurately delineate the mapped loci, while keeping the computing load manageable. The reads per non-overlapping 1 Mb window in the 115 RH samples was 237 ± 12 for human and 13 798 ± 234 for hamster.

### Statistical models

Changes in human or hamster reads were used to identify loci for growth, paclitaxel and their interaction. Since the data was both overdispersed and had batch effects of RH pool (**Supporting Information**), we used the negative binomial mixed model

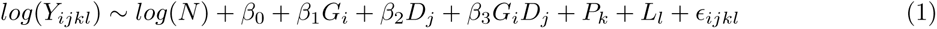

where *Y_ijkl_* is the number of species-specific human or hamster reads in each 1 Mb window, *N* is the total number of species-specific aligned human or hamster reads in each sample, respectively, with its logarithm representing the offset to correct for differences in sequence ascertainment across samples, *β*_0_ is the fixed effect of intercept, *β*_1_ is the fixed effect of growth *G* at time *i*, *β*_2_ is the fixed effect of drug *D* at concentration *j*, *β*_3_ is the fixed effect of the growth and drug interaction *G_i_D_j_*, *P* is the random intercept due to RH pool *k*, *L* is a random intercept nested inside *P_k_* to account for possible differences in drug histories of the RH pools, where *l* = *j·n* with *n* = 0 for week 0 and 1 for other times, and *E_ijkl_* is the residual variance. The addition of *L_l_* was precautionary, since the samples were well balanced with respect to drug history.

To identify additional human regulatory loci, reads from hamster mitochondria, human mitochondria and M. fermentans were added individually to the model as fixed effects.

Genome-wide significance thresholds were set by permutation to give a 5% family-wise error rate (FWER), equivalent to one expected false positive signal every 20 genome scans (Churchill and Doerge 1994).

Genome scans, related statistical analyses and graphical representations used overlapping 1 Mb windows with 10 kb steps. When noted, other statistical tests employed non-overlapping 1 Mb windows to be conservative. All errors in text are standard errors of the mean (s.e.m.).

### Mycoplasma contamination

The sequence data revealed appreciable levels of mycoplasma contamination in the A23 parental cells, as well as in the RH samples (**Tables S28–S31, Supporting Information**). However, including the *M. fermentans* copy number in our statistical model left the growth and paclitaxel loci essentially unchanged. In addition, no significant human loci were found that regulated the levels of the bacterium (all FDR = 1). Together, these results indicate that the mycoplasma contamination had little effect on the conclusions of our study.

### Accession numbers

The sequencing data is available from the Sequence Read Archive (SRA) (https://www.ncbi.nlm.nih.gov/sra/) under BioProject accession number PRJNA592253. Additional data and computer scripts are available from NIH figshare (https://nih.figshare.com/; http://dx.doi.org/10.35092/yhjc.11370204).

## Author contributions

AHK: Acquisition of data, analysis and interpretation of data, drafting or revising the article. AL: Analysis and interpretation of data. RTW: Analysis and interpretation of data. JSB: Analysis and interpretation of data. KL: Analysis and interpretation of data. DJS: Conception and design, analysis and interpretation of data, drafting or revising the article.

## Acknowledgments

We acknowledge the UCLA Semel Institute Neurosciences Genomics Core for sequencing. This work used computational and storage services associated with the Hoffman2 Shared Cluster provided by the UCLA Institute for Digital Research and Education Research Technology Group. We thank Jake Lusis for helpful comments on the manuscript. This study was supported by the National Institutes of Health (R21 HG007405), the Bennet Mills Allen Trust, the Edna Jones Endowment and the Department of Molecular and Medical Pharmacology, UCLA (to DJS).

## Supporting Information

### Cells

Human embryonic kidney 293 (HEK293) cells are female (Lin et al. 2014) and were obtained from the American Type Culture Collection (Manassas, VA) around 2006. We validated these cells in a previous study using simple tandem repeat genotyping (Khan et al. 2016).

Additional confirmation was obtained from the low pass genome sequence data generated during this study. A total of 1366 known common homozygous coding region single nucleotide polymorphisms (SNPs) had previously been identified in the HEK293 genome (Lin et al. 2014). Of these SNPs, a total of 640 were identified by our low pass sequencing (**Supporting Information**) and 637 (99.5%) agreed with the allelic variants in the HEK293 genome (*χ*^2^[640] = 982*, P <* 2.2 × 10^−16^, likelihood ratio test).

Similarly, 107 novel homozygous coding region SNPs had been previously identified in the HEK293 genome (Lin et al. 2014), of which 57 were identified in our low pass data. A total of 32 of these 57 SNPs (56%) agreed with the variants from the HEK293 genome (*χ*^2^[57] = 89*, P* = 4.5 × 10^−3^, likelihood ratio test).

Further, there was high concordance between the copy number alterations (CNAs) in our HEK293 isolate and the published HEK293 genome sequence (Lin et al. 2014) (*R* = 0.52, t[1,28173] = 101.1, *P <* 2.2 × 10^−16^) (**Figures 2C** and **S4**).

A23 hamster cells were kindly donated by Dr Christine Farr (University of Cambridge, Cambridge, UK), around 2004. These cells are a Chinese hamster lung fibroblast-derived cell line, a descendant of the DON cell line (Park et al. 2008) and are male. We had previously confirmed the A23 cells were of Chinese hamster origin by simple tandem repeat genotyping (Khan et al. 2016). The hamster origin of the cells was further verified by the stringent alignment to the published hamster genome (Rupp et al. 2018) of the low pass sequence data produced from this study.

HEK293 cells were grown in Dulbecco’s modified Eagle’s medium (DMEM) with 10% fetal bovine serum (FBS) and 1 × penicillin/streptomycin (100 units ml*^−^*^1^ penicillin, 100 *µ*g ml*^−^*^1^ streptomycin; Thermo Fisher Scientific®). The A23 cells were grown in *α* minimum essential medium (MEM), supplemented with 10% fetal bovine serum (FBS), 1 × penicillin/streptomycin. The RH pools were propagated in the same medium as the A23 cells, with the addition of 1 × HAT (100 *µ*M hypoxanthine, 0.4 *µ*M aminopterin, 16 *µ*M thymidine; Thermo Fisher Scientific®) (Park et al. 2008; Wang, Ahn, et al. 2011).

Cells were evaluated for mycoplasma contamination by aligning sequence data to the relevant bacterial genomes (**Materials and Methods**).

### Paclitaxel

Paclitaxel (Taxol®) was obtained from Tocris®. Dimethyl sulfoxide (DMSO) was from Thermo Fisher Scientific®. To aid solubilization, paclitaxel was dissolved in DMSO and then diluted in medium. The final concentration of DMSO in all samples was 0.37%.

### Cell fusion

We performed cell fusions as described (Schafer and Farr 1998), with modifications. Four separate fusion reactions of equal size were used to constitute each pool, with irradiation employing either 30 Gray (Gy) or 100 Gy from a Shepherd Mark I ^137^_55_Cs Irradiator (**Table S1**). The reactions were seeded into HAT selective medium using either five or ten 75 cm^2^ tissue flasks (1:5 and 1:10 dilutions, respectively), with the conditions applied in balanced fashion.

The expected fragment size for the two radiation doses was 10 Mb and 4 Mb, respectively (Cox et al. 1990; Hudson et al. 2001; Kwitek et al. 2004; McCarthy 1996; McCarthy et al. 2000; Olivier et al. 2001; Walter et al. 1994). The mean fragment size weighted by clone number at each dose was 7.4 Mb ±0.1 Mb for each pool. To evaluate the frequency of spontaneous revertants, mock fused A23 cells were also plated. When the four fusion reactions for each of the six pools reached confluence, the cells were consolidated and a small portion reserved for genotyping. These samples constitute week 0 and 0 nM paclitaxel and are referred to as the “RH pools” (**Table S2**). The remainder of the pools were grown in HAT medium for 1, 2, 3, 4 or 6 weeks, supplemented with 0, 8, 25 or 75 nM paclitaxel, and analyzed using low pass sequencing.

### Clone, colony and revertant counts

#### RH clone counts

A negative binomial mixed model was used to evaluate the RH clone counts (**Supporting Information**), since these data were significantly overdispersed (dispersion = 25, *χ*^2^[23] = 570, *P <* 2.2 × 10^−16^, Poisson model) and there was a potential batch effect of RH pool (**Figures S1A and S11, Table S1**). Fixed and interaction effects of dilution and radiation dose were employed, together with a random intercept of pool. The overdispersion was normalized towards unity by the negative binomial model (dispersion = 0.70, *χ*^2^[22] = 15, *P* = 0.85). Consistent with a batch effect of RH pool, the random intercept significantly improved model fit (Akaike information criterion, AIC = 387 without random effect, 378 with random effect; *χ*^2^[1] = 11, *P* = 9.5 × 10^−4^, likelihood ratio test). The adjusted between- and within-pool variance accounted for 69% (intraclass correlation coefficient, ICC = 0.69) and 31% of the total variance components, respectively.

The higher radiation dose resulted in a significantly smaller number of RH clones (t[1,22] = 2.4*, P* = 0.02; average conditional effect) (**Figure S1A**). We explored possible cost savings in plasticware and culture medium by using both a 1:5 dilution as well as the 1:10 dilution described in published protocols (Schafer and Farr 1998). The recommended 1:10 dilution was superior to the 1:5 dilution, providing a highly significant 2.6 ± 0.3-fold increase in the number of RH clones (t[1,22] = 8.5*, P* = 2.3 × 10^−8^; average conditional effect) (**Figure S1A**). The interaction of radiation dose and dilution was non-significant (t[1,22] = 0.69*, P* = 0.50).

#### The RH clone, colony and revertant counts are correlated

The number of RH clones was highly correlated with both the total number of colonies (*R* = 0.99, t[1,22] = 34, *P <* 2.2 × 10^−16^) and the number of revertants (*R* = 0.85, t[1,22] = 7.5, *P* = 1.7 × 10^−7^). However, there were some differences in the effects of radiation dose and dilution on the colonies and revertants compared to the RH clones.

#### Colony counts

Similar to the RH clones, the colonies showed significant overdispersion (dispersion = 20, *χ*^2^[23] = 5 × 10^2^, *P <* 2.2 × 10^−16^, Poisson model), which was normalized towards unity by the negative binomial model (dispersion = 0.72, *χ*^2^[22] = 16, *P* = 0.82) (**Table S1**). In addition, the random effect of RH pool significantly improved model fit (AIC = 400 without random effects, 384 with random effects; *χ*^2^[1] = 18, *P* = 2.0 × 10^−5^, likelihood ratio test, ICC = 0.82) (**Figures S1B and S11**).

Like the RH clones, the higher radiation dose significantly decreased colony numbers (t[1,22] = 2.8*, P* = 0.01; average conditional effect) (**Figure S1B**). However, the dominant effect was again dilution, with the higher dilution giving larger colony counts (t[1,22] = 11*, P* = 1.9 × 10^−10^; average conditional effect) (**Figure S1B**). Interactions of dilution and radiation dose were insignificant (t[1,22] = 0.06*, P* = 0.96).

#### Revertant counts

There was also significant overdispersion for the revertants (dispersion = 3.5, *χ*^2^[23] = 80, *P* = 2.7 × 10^−8^, Poisson model), which was normalized towards unity by the negative binomial model (dispersion = 0.72, *χ*^2^[22] = 16, *P* = 0.82) (**Table S1**). The random effect of RH pool significantly improved model fit (AIC = 339 without random effects, 318 with random effects; *χ*^2^[1] = 23, *P* = 1.9 × 10^−6^, likelihood ratio test, ICC = 0.86) (**Figures S1C and S11**).

There were no significant fixed effects of radiation for the revertants (t[1,22] = 1.5*, P* = 0.15; average conditional effect) (**Figure S1C**). The most significant effect was dilution, with higher dilution giving larger numbers (t[1,22] = 11*, P* = 3.5 × 10^−10^; average conditional effect) (**Figure S1C**). Interactions of dilution and radiation dose were significant for the revertants (t[1,22] = 2.2*, P* = 0.04).

### Low pass sequencing

Genotyping employed Illumina sequencing with single-end reads of 64 bp. As background information, we obtained 41.7 M sequence reads from the HEK293 donor cells and 37.2 M reads from the A23 recipient cells (**Tables S3 and S4**). In addition, we obtained 41.1 ± 2.8 M reads for each of the six RH pools (**Table S5**). For all RH samples under the various conditions of growth and drug, including the six RH pools, we obtained 40.5 ± 0.7 M reads per sample, corresponding to a sequencing depth of 0.84 ± 0.01 times the human genome.

#### Human and hamster genomes

The human genome is a high quality finished assembly. The hamster genome is a high quality draft assembly with a scaffold N50 of 20.2 Mb, L50 of 32, 0.12% gaps and 96.6% of the sequence assigned to a chromosome (Rupp et al. 2018). In fact, the continuity of the hamster genome is superior to the rat and approaches that of the mouse. However, using selected human/hamster sequence variants would neglect genome regions that could not be aligned between the two species. We therefore decided to maximize information recovery by aligning all sequence reads. Stringent alignment parameters were employed, so that only a small proportion of reads mapped across species (**Tables S3 and S4, Materials and Methods**). Using the HEK293 and A23 sequence data, the misalignment rate for the human to hamster nuclear genomes was 0.13% and 0.27% in the other direction. For the mitochondrial genomes, the human to hamster misalignment rate was 0.19% and 0.14% reversed.

#### Alignment strategies for human DNA

Although only 0.27% of hamster reads also aligned to the human nuclear genome (**Table S3**), this low frequency became non-negligible in the context of the six RH pools, where there was a mean of 59 ± 14-fold more mapped hamster reads than human (**Tables S5** and **S6**). To evaluate the human DNA in the RH pools with greater specificity, we discarded reads that aligned to both species (**Figure 1**). A mean of 0.88 ± 0.11 M reads initially aligned to the human genome (**Table S5**). After discarding 0.13 ± 0.03 M reads that aligned to both species (a hamster-to-human misalignment rate of 16.8% ± 3.7%), 0.74 ± 0.12 M reads specifically mapped to the human genome in the six RH pools. The corresponding density of human reads was 0.015 ± 0.0024 times the human genome.

For all samples under the various conditions of growth time and drug exposure, including the six RH pools, there was a mean of 84 ± 16-fold more mapped hamster reads than human. A mean of 0.90 ± 0.04 M reads initially aligned to the human genome. Of these, 0.16 ± 0.005 M reads (21.0% ± 1.1%) also aligned to the hamster genome and were discarded, giving 0.75 ± 0.04 M reads specifically aligned to the human genome. The corresponding density of human reads was 0.015 ± 0.0008 times the human genome.

#### Alignment strategies for hamster DNA

A total of 0.13% of human reads from HEK293 cells also mapped to hamster (**Table S3**). However, the much smaller amount of human DNA in the RH samples meant that the impact of cross-species alignment was much less for the hamster genome than the human. Nevertheless, to accurately quantitate the levels of hamster DNA, we discarded hamster reads that cross-aligned to both species (**Figure 1**).

A mean of 36.4±2.5 M reads initially aligned to the hamster genome in the six RH pools, giving 36.2±2.5 M reads specific to hamster after discarding 0.13±0.03 M reads that aligned to both hamster and human (a cross-alignment rate of 0.36%±0.06%) (**Table S6**). The corresponding read density was 0.98 times the hamster genome ±0.07.

For all samples, a mean of 35.8±0.6 M reads initially aligned to the hamster genome. Of these, 0.16±0.005 M reads (0.45%±0.01%) also aligned to the human genome and were discarded, giving 35.6±0.6 M reads specifically aligned to the hamster genome. The corresponding density of hamster reads was 0.96±0.02 times the hamster genome. All analyses employed species-specific read alignments, unless otherwise noted.

#### Alignment strategies for mitochondrial genomes

A total of 0.19% of human reads cross-aligned with hamster mitochondria and 0.14% of hamster reads cross-aligned with human mitochondria (**Table S4**). After correcting for cross-species alignments, 172±99 reads mapped to the human mitochondrial genome in the six RH pools (**Table S21**). For all samples under the various conditions of growth time and drug concentration, including the six RH pools, 68±15 reads aligned to the human mitochondrial genome. The low numbers of human mitochondrial sequence reads suggest that these organelles were transferred with low efficiency to the hamster cells (**Results**).

After correcting for cross-species alignments, 92926±6797 reads mapped to the hamster mitochondrial genome for each of the six RH pools (**Table S21**). For all samples under the various conditions of growth time and drug concentration, including the six RH pools, 90434 ± 3033 reads aligned to the hamster mitochondrial genome.

### DNA retention in the RH pools

#### The hamster genome is unchanged in RH cells

There was strong similarity between copy number alterations (CNAs) in the parental A23 cells and the RH pools, suggesting that the hamster genome in the six RH pools was not substantially altered by the cell fusions (*R* = 0.98, t[1,2041] = 253, *P <* 2.2 × 10^−16^, non-overlapping 1 Mb windows) (**Figures 2A, 2B, S2 and S3A**).

#### Regions of enhanced human DNA retention in the RH cells

While the recipient hamster genome was largely unchanged by the cell fusion, human DNA donated to the RH cells was retained as a selection of extra DNA fragments (**Figures 2C, 2D and S5**). Since the probability of human gene transfer from the donor cells increases with copy number, there was nevertheless a significant correlation between human DNA retention in the RH pools and copy number in the HEK293 cells (*R* = 0.47, t[1,3007] = 29, *P <* 2.2 × 10^−16^, non-overlapping 1 Mb windows) (**Figure S3B**). The thymidine kinase (TK1) gene was used as a selectable marker and therefore has a retention frequency of one (**Figure 2F**).

Donor centromeres and telomeres confer increased stability on the human DNA fragments in RH cells (Wang, Ahn, et al. 2011). In fact, nearly all centromeres showed significantly increased retention in the RH pools (centromere to non-centromere read ratio = 1.1 in HEK293 cells; 5.5 ± 0.2 in the six RH pools; t[1,5] = 7.4, *P* = 7.0 × 10^−4^, Two Sample t-test with pooled variance) (**Figures 2C, 2D and S5**). The combination of pooling and low-pass sequencing employed in this study more clearly demonstrated augmented centromere retention in the RH cells than older technologies such as arrays or PCR, which had few centromeric markers. For chromosome 6, the centromere copy number was greater than 1, indicating highly efficient retention and replication (**Figures 2D and S5**). However, centromere sequences represent an average of tandem chromosome-specific *α*-satellite repeats (Contreras-Galindo et al. 2017; Schneider et al. 2017) and caution is warranted when interpreting their retention values.

#### Human DNA retention frequency

Non-centromeric regions showed relatively constant retention across the genome, as previously documented in RH panels (Avner et al. 2001; Hudson et al. 2001; Kwitek et al. 2004; McCarthy et al. 2000; Olivier et al. 2001) (**Figures 2D** and **S5**). We calculated the overall retention of the human genome in the RH pools in two ways (**Table S7**). The first compared the median number of human-specific sequence reads in non-overlapping 1 Mb windows to the median number of hamster-specific reads. A correction multiplier of two was used to account for the fact that the retained human fragments are overwhelmingly haploid, while the hamster genome is diploid. The retention frequency, *R*, is thus 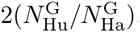, where 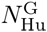 is the median number of human-specific reads in non-overlapping 1 Mb genome windows and 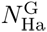 is the median number of hamster-specific reads.

The second approach assigned a retention of 100% to the human TK1 gene. Scaled values of the human-specific reads in each non-overlapping 1 Mb window were then used as a measure of retention frequency. A correction factor was applied to account for the revertants. Thus, 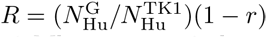, where 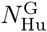 is the median number of human-specific reads in non-overlapping 1 Mb genome windows, 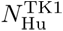 is the maximum number of human-specific reads in a 1 Mb window encompassing TK1, and *r* is the reversion frequency (**Table S1**).

Using the first method, the mean of the median fraction of the human genome that was retained in the six RH pools was 2.6% ± 0.5% and using the second 5.3%±0.3% (t[1,8.7] = 4.5, *P* = 1.6 × 10^−3^, Welch Two Sample t-test) (**Table S7**). The mean retention frequency using the two methods was 3.9%±0.3%.

The lowest euchromatic retention frequency was at 54 530 000 bp on chromosome 19 and was significantly greater than zero (0.4%±0.1%, t[1,5] = 4.2, *P* = 8.2 × 10^−3^ using sequence alignments; 0.9%±0.1%, t[1,5] = 7.8, *P* = 5.7 × 10^−4^ using TK1; 0.7% ± 0.1%, t[1,5] = 6.7, *P* = 1.1 × 10^−3^, using mean of sequence alignments and TK1; One Sample t-tests).

### Further details on copy number changes in the RH samples

After 1, 2, 3, 4 and 6 weeks of growth in HAT medium supplemented with 0, 8, 25 or 75 nM paclitaxel, the RH samples were analyzed by low pass sequencing. There were a total of 115 samples, including the six RH pools (**Table S2**). We obtained data for all permutations of time and drug, except one, corresponding to 91.3% ± 5.3% of all possible samples for each table cell.

#### Overall human copy number

A Gaussian linear mixed model was used to evaluate changes in overall human DNA retention in the RH samples. We employed fixed effects of growth time, paclitaxel concentration and their interaction, together with random effects of RH pool and nested drug history (**Figures S7–S9, Supporting Information**). There were significant decreases in retention as a result of growth (2.6% ± 0.5% at week 0 to 1.0% ± 0.2% at week 6, t[1,91.2] = 8.8*, P* = 6.8 × 10^−14^, using sequence alignments; 5.3% ± 0.3% at week 0 to 1.3% ± 0.2% at week 6, t[1,91.5] = 15*, P <* 2.2 × 10^−16^, using TK1; 3.9% ± 0.3% at week 0 to 1.1% ± 0.2% at week 6, t[1,90.8] = 15*, P <* 2.2 × 10^−16^, using mean of sequence alignments and TK1; Kenward-Roger degrees of freedom, df; average conditional effects), but not paclitaxel (t[1,24.5] = 1.7*, P* = 0.1, sequence alignments; t[1,24.7] = 0.07*, P* = 0.9, TK1; t[1,24.4] = 0.8*, P* = 0.5, mean; Kenward-Roger df; average conditional effects). The interaction was also insignificant (t[1,93.2] = 0.8*, P* = 0.5, sequence alignments; t[1,93.6] = 0.3*, P* = 0.7, TK1; t[1,92.8] = 0.7*, P* = 0.5, mean; Kenward-Roger df) (**Figures S7–S9**).

There was inconsistent significance of the random effect of pool (*χ*^2^[1] = 13*, P* = 4.1 × 10^−4^, ICC = 0.49, sequence alignments; *χ*^2^[1] = 4.4*, P* = 0.04, ICC = 0.26, TK1; *χ*^2^[1] = 1.8*, P* = 0.17, ICC = 0.18, mean; likelihood ratio tests) and also drug history (*χ*^2^[1] = 2.9*, P* = 0.09, ICC = 0.18, sequence alignments; *χ*^2^[1] = 4.9*, P* = 0.03, ICC = 0.23, TK1; *χ*^2^[1] = 4.7*, P* = 0.03, ICC = 0.33, mean; likelihood ratio tests).

#### Human and hamster copy number changes across genome

The hamster genome was relatively stable, as revealed by sequence alignments (**Figures 3** and **S10**). In contrast, the human genome showed significant decreases in average copy number and increases in variance as a result of growth or paclitaxel, consistent with selective pressure.

Congruent with the overall decrease in human DNA retention in the RH samples (**Figures S7–S9**), we found significant genome-wide decreases in relative normalized human copy number at week 4 due to both growth (mean log_2_ copy number change = −0.54 ± 0.01, t[1,2911] = 48, *P <* 2.2 × 10^−16^, One Sample t-test, 4 weeks compared to 0 weeks, 0 nM paclitaxel) (**Figures 3A and 3B**) and also paclitaxel (mean log_2_ copy number change = −2.14 ± 0.02, t[1,2723] = 87, *P <* 2.2 × 10^−16^, One Sample t-test, 75 nM compared to 0 nM paclitaxel, 4 weeks) (**Figures 3C** and **3D**).

Significant genome-wide decreases in relative normalized human copy number were also found at week 6 for both growth (mean log_2_ copy number change = −0.78 ± 0.01, t[1,2911] = 54.1, *P <* 2.2 × 10^−16^, One Sample t-test, 6 weeks compared to 0 weeks, 0 nM paclitaxel) (**Figures S10A and S10B**) and paclitaxel (mean log_2_ copy number change = −1.46 ± 0.02, t[1,2781] = 64, *P <* 2.2 × 10^−16^, One Sample t-test, 75 nM compared to 0 nM paclitaxel, 6 weeks) (**Figures S10C and S10D**).

The variance of relative normalized copy number changes for human DNA was significantly greater than hamster for both growth and paclitaxel at 4 weeks. The standard deviation (s.d.) of log_2_ copy number changes for 4 weeks of growth compared to 0 weeks (0 nM paclitaxel) was 0.05 (±4.1 × 10^−7^) for the hamster genome and 0.60 (±2.6 × 10^−6^) for the human (F[1,4953] = 3277, *P <* 2.2 × 10^−16^, Levene’s test) (**Figures 3A and 3B, Tables S8 and S9**). Similarly, the s.d. of log_2_ copy number changes at 75 nM compared to 0 nM paclitaxel (4 weeks) was 0.10 (±7.6 × 10^−7^) for the hamster genome and 1.3 (±6.1 × 10^−6^) for the human (F[1,4765] = 2754, *P <* 2.2 × 10^−16^, Levene’s test) (**Figures 3C and 3D**).

The variance of relative normalized copy number changes for human DNA was also significantly greater than hamster at 6 weeks. The s.d. of log_2_ copy number changes for 6 weeks of growth compared to 0 weeks (0 nM paclitaxel) was 0.07 (± 5.7 *×* × 10^−7^) for the hamster genome and 0.78 (± 3.3 *×* × 10^−6^) for the human (F[1,4953] = 3079, *P <* 2.2 *×* × 10^−16^, Levene’s test) (**Figures S10A and S10B, Tables S8 and S9**). Similarly, the s.d. of log_2_ copy number changes was 0.11 (± 8.2 *×* × 10^−7^) for the hamster genome at 75 nM compared to 0 nM paclitaxel (6 weeks growth) and 1.2 (± 5.5 *×* × 10^−6^) for the human (F[1,4823] = 2758, *P <* 2.2 *×* × 10^−16^, Levene’s test) (**Figures S10C and S10D**).

### Further details on statistical models

#### Mixed models

Genes that confer a survival advantage as a result of growth or paclitaxel will show increased copy number and vice versa. We identified human and hamster loci for growth and paclitaxel by evaluating the significance of copy number changes across the genome using a negative binomial mixed model to control for both overdispersion (**Supporting Information**) and batch effects of RH pool (**Figure S11**).

Fixed effects of growth time, paclitaxel concentration and their interaction were employed, together with random intercepts of RH pool and a nested term to account for potential treatment differences between pools (**Equation 1, Materials and Methods**). Genome scans used overlapping 1 Mb windows with 10 kb steps (**Supporting Information**).

The mixed model was analyzed using R 3.5.1 (R Core Team 2018) and the gam function of the mgcv package (Wood 2017) employing restricted maximum likelihood (REML) estimation. We obtained *P* values using the glht function of the multcomp package (Hothorn et al. 2008). Growth loci were identified employing the null hypotheses of no copy changes at 0, 8, 25 and 75 nM paclitaxel and an average conditional effect at 27 nM paclitaxel. Paclitaxel loci were identified using null hypotheses of no copy changes at 1, 2, 3, 4 and 6 weeks and an average conditional effect at 3.2 weeks. Interaction *P* values and loci were also calculated.

To identify additional human regulatory loci, reads from hamster mitochondria, human mitochondria and *M. fermentans* were added individually to the model as fixed effects. Each set of genome scans took ~2 h on a computer cluster.

Additional analyses of count data using negative binomial mixed models employed glmmTMB (Brooks et al. 2017), while Gaussian linear mixed model analyses of non-count data used the lmer function of the lme4 package without an offset (Bates et al. 2015). Post hoc analyses to obtain *P* values employed emmeans (Lenth 2019).

#### Model fit for human loci

There was significant overdispersion of the data used to identify human growth and paclitaxel loci (dispersion = 31.7 ± 0.4 Poisson model, non-overlapping 1 Mb windows), which was normalized towards unity by the negative binomial mixed model (dispersion = 0.96 ± 4.0 × 10^−3^ negative binomial; t[1,3113] = 69, *P <* 2.2 × 10^−16^, non-overlapping 1 Mb windows).

The fit of the negative binomial mixed model was significantly improved by the random effects (Akaike information criterion, AIC = 1311 ± 6 without random effects, 1182 ± 5 with random effects, t[1,6182] = 17, *P <* 2.2 × 10^−16^, non-overlapping 1 Mb windows) (**Figure S11**). Both the pool (*χ*^2^ = 874 ± 14, reference df = 5, −log_10_ *P* = 18.0 ± 0.3, non-overlapping 1 Mb windows) and drug history random effects were significant (*χ*^2^ = 226 ± 4, reference df = 28, −log_10_ *P* = 2.5 ± 0.03, non-overlapping 1 Mb windows).

Visual inspection of residuals in eight randomly selected genome positions suggested satisfactory performance of the model, including control of overdispersion and RH pool batch effects (**Figures S12–S17**). These conclusions were supported by a goodness-of-fit analysis for the whole dataset, which showed no significant departure of the residuals from normal (*χ*^2^ = 92.9 ± 0.4, df = 93.6 ± 0.1, −log_10_ *P* = 0.43 ± 2.1 *×* × 10^−3^, likelihood ratio test, non-overlapping 1 Mb windows).

#### Significance thresholds

Permutation significance thresholds for human and hamster loci were obtained by co-shuffling *Y_ijkl_* and *N* within growth (*G_i_*), drug (*D_j_*) and pools (*P_k_*) in **Equation 1 (Materials and Methods)**, thus keeping the relationship between *Y_ijkl_* and *N* intact within the fixed and random effects. More than 1000 shuffles of the genome were performed for all overlapping 1 Mb windows, taking ~25 days on a computer cluster for each of the two genome scans. The maximum −log_10_ *P* values were chosen from each shuffle and genome-wide significance thresholds set by choosing the 95^th^ percentile of the 1000 maximum values. This threshold is expected to give one false −log_10_ *P* value in 5% of genome scans, or one false positive every 20 scans (Churchill and Doerge 1994).

Significance thresholds for hamster mitochondria, human mitochondria and *M. fermentans* were obtained by permutation of *Y_ijkl_* and *N* within *P_k_*, taking *∼*15 days for each of the three genome scans.

#### Peak finding

Loci were identified using an automated peak finding algorithm that allowed the minimum distance between peaks to be adjusted (**Tables S12 and S17, Results**). Comparison with handpicked growth loci on chromosomes 1–3 (0 nM paclitaxel) showed an optimum concordance of 68% ± 6% when the minimum distance between peaks was set at 2 Mb. The algorithm was more conservative (35 ± 0.6 loci, first three chromosomes) than manual peak finding (46 ± 3.5 loci), but this difference was not significant t[1,2.1] = 3.2, *P* = 0.08, Welch Two Sample t-test).

### Reproducibility

To assess replicability, we used a cross-validation procedure in which non-overlapping halves of the six RH pools were compared using the intersection of significant 1 Mb windows at various false discovery rate (FDR) thresholds (**Figure S18**) (Benjamini and Hochberg 1995). Highly significant overlaps were found, suggesting good reproducibility (summary logarithm of the odds ratio = 0.56 at FDR = 0.01, 95% confidence intervals = 0.37 and 0.75, t[1,10.0] = 5.7, *P* = 2.0 × 10^−4^, Kenward-Roger df).

### Further details on growth loci

#### Growth loci tally

The number of growth loci at the five paclitaxel levels ranged between 55 and 460, giving a total of 1836 loci (**Figures 4 and S19, Tables S10–S13**). However, there was significant overlap between the growth loci at the various paclitaxel concentrations (**Figure S20, Tables S14 and S15, Supporting Information**). There were 859 unique growth loci after selecting representatives with the highest −log_10_ *P* values, ~1.4% of all known genes (**Tables S12 and S16**).

Seven of the 859 unique growth loci (SEMA3A, TOR1A, ATXN8OS, RN7SL584P, DNAH17, DNAH17-AS1 and AL050309.1) had positive growth coefficients (0.11±0.02); the remaining coefficients were negative (−0.24±2.4 × 10^−3^, compared to −0.15±0.0 in all non-overlapping 1 Mb windows) (**Figure S23A**). These observations are consistent with the loss of human DNA in the RH samples as a result of growth (**Figures 3, S7–S10**).

#### Overlap of growth loci

There were significant correlations between the −log_10_ *P* values for growth at different paclitaxel concentrations (weakest correlation, 0 nM and 75 nM; *R* = 0.21, t[1,3052] = 12.134, *P <* 2.2 × 10^−16^; mean *R* = 0.73 ± 0.09; non-overlapping 1 Mb windows) (**Figure S20, Table S14**). However, the average correlation was decreased for the 75 nM samples (*R* = 0.44±0.10) compared to the others (*R* = 0.93±0.02; t[1,3.3] = 4.8, *P* = 0.01, Welch Two Sample t-test).

The generally high similarity of −log_10_ *P* values for growth at different paclitaxel concentrations was reflected in the sharing of significant non-unique growth loci (least significant overlap, 0 nM and 75 nM; odds ratio = 8.5*, P* = 6.2 × 10^−3^, Fisher’s Exact Test) (**Figure S20, Table S15**). There was decreased overlap of loci at 75 nM (odds ratio = 33 ± 13) compared to the other levels (odds ratio = 1799 ± 1441), but the difference was insignificant (t[1,5.0] = 1.2, *P* = 0.28, Welch Two Sample t-test).

#### Growth centromeres

Four of the unique growth loci (0.5%) mapped to centromeres on chromosomes 1, 11, 15 and X. These loci had significantly lower −log_10_ *P* values than the unique non-centromeric loci (centromere −log_10_ *P* = 19.3 ± 1.3, non-centromere = 23.7 ± 0.5, t[1,4.0] = 3.2, *P* = 0.03, Welch Two Sample t-test).

#### Coding and non-coding growth loci

Of the 855 unique non-centromeric RH growth loci, 442 (52%) were coding and 413 (48%) were non-coding, a significant enrichment in coding genes compared to the null of all genome positions (44% coding, 56% non-coding; *χ*^2^[1] = 19.5*, P* = 9.9 × 10^−6^). However, there was no significant difference in −log_10_ *P* values for the unique coding and non-coding loci (mean −log_10_ *P* = 23.5 ± 0.6 for coding, 24.0 ± 0.8 for non-coding; t[1, 825] = 0.4, *P* = 0.7, Welch Two Sample t-test). These observations suggest that while coding genes are enriched, non-coding genes still make appreciable contributions to cell growth.

Close-ups of growth loci at the chromosome level are shown in **Figures 5A–5C**. Copy number changes are shown for six significant loci in **Figure 5D**. Loci displayed sharp mapping resolution, with the −2log_10_ *P* distance being,:S 100 kb (**Figures 5E–5F**). However, loci that mapped to large genes (≥ 1 Mb) tended to display a plateau at the top of their −log_10_ *P* peak, a testament to a high resolution mapping technique that can “trace” the profile of a lengthy gene (**Figure S22**).

#### Chromosomal distribution of growth loci

The number of unique growth loci (excluding centromeres) compared to the number of genes on each chromosome is shown in **Figure S21A**. The distribution of loci was non-uniform (*χ*^2^[22] = 75*, P* = 9.8 × 10^−8^), with significant surfeits on chromosomes 2, 4–5 and X (least significant, chromosome X; *χ*^2^[1] = 4.0*, P* = 0.046) and significant deficits on chromosomes 14, 17 and 19–21 (least significant, chromosome 20; *χ*^2^[1] = 4.7*, P* = 0.03).

#### Functional enrichment of RH growth genes

The unique RH growth genes were enriched in ten categories of the functional annotation chart of DAVID including alternative splicing, splice variant and phosphoprotein (FDR *<* 0.05) (Benjamini and Hochberg 1995; Huang et al. 2009). Further, 31 terms of the biological process category of the gene ontology (GO) were enriched, including neurogenesis, embryonic development and cellular biosynthesis (FDR *<* 0.05) (Mi et al. 2019; The Gene Ontology Consortium 2019). Cell adhesion was significant in both DAVID (observed = 24, expected = 10, odds ratio = 2.5, *P* = 2.5 × 10^−4^, EASE modified Fisher’s Exact Test, FDR = 0.02) and GO (observed = 43, expected = 19, odds ratio = 2.5, *P* = 1.7 × 10^−6^, Fisher’s Exact Test, FDR = 4.4 × 10^−3^), a possible consequence of selection for the adherent RH cells. Essential coding genes identified in two loss-of-function CRISPR screens were enriched in housekeeping terms related to cell division, transcription, splicing and translation (Hart et al. 2015; Wang, Birsoy, et al. 2015), overlapping with the RH growth genes in the splicing terms.

#### Novel RH growth genes

The non-redundant coding RH growth genes had significantly fewer literature citations in the GeneRIF database (mean citations = 398 ± 15) than coding non-growth genes (896 ± 72; t[1,164] = 6.8, *P* = 2.3 × 10^−10^, Welch Two Sample t-test at peak significance threshold of 364 publications, FDR = 2.3 × 10^−9^) (**Figures S28A and S28B**) (Jimeno-Yepes et al. 2013). Similarly, there were fewer entries in the Reactome database for unique coding RH growth genes (mean citations = 249 ± 20) than coding non-growth genes (400 ± 20), which reached nominal but not FDR corrected significance (t[1,4.3] = 5.4, *P* = 4.7 × 10^−3^, Welch Two Sample t-test at peak significance threshold of 113 entries, FDR = 0.09) (**Figures S28C and S28D**) (Fabregat et al. 2018).

Of the 442 unique coding region RH growth genes, 148 had neither literature citations in GeneRIF nor entries in the Reactome database. These genes showed significant enrichment of one protein domain, the WW domain (observed = 6, expected = 0.4, odds ratio = 18.9, *P* = 3.3 × 10^−5^, EASE modified Fisher’s Exact Test, FDR = 9.6 × 10^−3^), as judged by the InterPro database (Huang et al. 2009; Mitchell et al. 2019).

None of the 413 unique non-centromeric non-coding RH genes had entries in either the GeneRIF or Reactome databases.

#### Olfactory receptor genes

Reports have suggested that olfactory receptors may play a role in cell proliferation and differentiation (Kang and Koo 2012; Tsai et al. 2017). The unique RH growth loci were found in olfactory receptor gene clusters on six chromosomes: 3, 11, 12, 14, 19 and X. One locus with −log_10_ *P* = 21.5 mapped within the chromosome 19 cluster to AC005255.1 (OR7C1) (**Figure S29A**), an olfactory receptor which has been shown to play a role in maintenance of colon cancer cells (Morita et al. 2016). Although 19 out of 298 olfactory receptor genes displayed a growth phenotype in at least one of the five cell lines in a recent CRISPR screen (Hart et al. 2015), none overlapped with the RH growth genes. This observation suggests complementary gain- and loss-of-function roles for olfactory receptors in growth.

#### Gene deserts

“Gene deserts” are defined as regions of the genome *>*500 kb that lack known genes (Salzburger et al. 2009). The majority of unique RH growth loci mapped close to a gene (mean distance = 1.7 kb 1.4 kb). However, two growth loci were located in gene deserts, with the nearest gene *>* 250 kb away (**Figures S29B and S29C**).

The RH gene desert growth locus at 104 730 000 bp on chromosome 1 (**Figure S29B**) was 11 897 bp away from variant rs10494021 (*R*^2^ ∼ 1) (1000 Genomes Project Consortium et al. 2015). This polymorphism had suggestive significance (odds ratio = 1.14, P = 6.61 × 10^−6^) in a GWAS for childhood ear infection (Tian et al. 2017). Thus, a possible candidate for this clinical trait may be a novel RH gene rather than a known, but distant, gene regulated by rs10494021.

No significant difference was found in the −log_10_ *P* values for the unique growth loci in gene deserts (mean = 45.5±29.5) compared to the other non-centromeric growth loci (mean = 23.7±0.5; t[1,1.0] = 0.74, *P* = 0.59, Welch Two Sample t-test). Although there were only two RH gene desert loci and test power was low, the similarity of −log_10_ *P* values lends some credence to these unusual loci.

### Further details on paclitaxel loci

#### Paclitaxel loci tally

There were a total of 97 paclitaxel loci (**Figures 4B, S20 and S30, Tables S10, S11, S14, S15, S17 and S18**). After accounting for overlap at the various growth times and selecting representatives with the highest −log_10_ *P* values, there were 38 non-redundant paclitaxel loci (**Tables S17 and S19**).

Seven of the 38 unique paclitaxel loci (NEK10, RPL7A, DNAJC1, TBC1D12, ZBED5-AS1, GALNT18, AC013762.1) had positive coefficients (1.7 × 10^−2^±2.4 × 10^−3^) indicating increased copy number as a result of drug exposure. The remaining non-redundant paclitaxel loci had negative coefficients (−2.1 × 10^−2^±1.7 × 10^−3^; null of all non-overlapping 1 Mb windows, 2.0 × 10^−4^±4.5 × 10^−5^) (**Figure S23B**). The proportion of unique paclitaxel loci with positive coefficients was significantly greater than the unique growth loci (seven out of 859 unique growth loci; log-likelihood ratio = −15.8, *P* = 2.4 × 10^−8^, Multinomial Goodness-Of-Fit test).

#### Overlap of paclitaxel loci

There was a significant increase in the number of non-unique paclitaxel loci with time, increasing from zero at week 1 to 25 at week 6 (*χ*^2^[4] = 43.7, P = 7.4 × 10^−9^) (**Figures 4B and S30, Tables S17 and S18**). There was also significant overlap of the paclitaxel loci at the different times (least significant overlap, weeks 2 and 3, odds ratio =, *P* = 1.8 × 10^−7^, Fisher’s Exact Test) (**Figure S20, Tables S14 and S15**), although there were also non-overlapping loci, suggesting that the various drug exposure times bring different genes into play (Khan et al. 2016; Wang and Kruglyak 2014).

The overlap of paclitaxel loci was reflected in significant correlations between −log_10_ *P* values for paclitaxel at different times (**Figure S20, Table S14**) (weakest significant correlation, week 1 and average conditional effect, *R* = 0.075, t[1,3052] = 4.2, *P* = 3.2 × 10^−5^; mean *R* = 0.62 ± 0.10, including insignificant comparisons between week 1 and weeks 4 or 6; non-overlapping 1 Mb windows).

#### Shared RH growth and paclitaxel loci

We evaluated the correlations between the −log_10_ *P* values for growth and paclitaxel (**Figure S32A, Table S14**). As expected, the correlations were significantly higher for growth at 75 nM paclitaxel (*R* = 0.27 ± 0.06) than growth at other paclitaxel levels (*R* = 0.01 ± 0.02, non-overlapping 1 Mb windows, t[1,6.6] = 3.7, *P* = 8.8 × 10^−3^, Welch Two Sample t-test).

There was also significant sharing between the 38 unique paclitaxel loci and 859 unique growth loci with 10 loci in common (odds ratio = 25.1*, P* = 1.0 × 10^−10^, Fisher’s Exact Test) (**Figure S31**). As expected, the overlap between unique growth and paclitaxel genes was greatest at the highest growth time (week 6) and paclitaxel concentration (75 nM) (odds ratio = 711*, P* = 2.2 × 10^−16^, Fisher’s Exact Test). Similar results were obtained for the non-unique growth and non-unique paclitaxel loci, with the highest overlap at week 6 growth and 75 nM paclitaxel (odds ratio = 490*, P <* 2.2 × 10^−16^, Fisher’s Exact Test) (**Figure S32B, Table S15**).

#### Paclitaxel centromeres

Four of the 38 unique paclitaxel loci were centromeres, located on chromosomes 11, 16, 20 and X. There was no significant difference in −log_10_ *P* values for the unique centromeric (−log_10_ *P* = 26.3 ± 5.6) and non-centromeric paclitaxel loci (−log_10_ *P* = 16.6 ± 1.2; t[1,3.3] = 1.7, *P* = 0.18, Welch Two Sample t-test). The frequency of centromeres in the unique paclitaxel loci (four out of 38, 10.5%) was significantly higher than for the unique growth loci (four centromeres out of 859 loci, 0.5%; log-likelihood ratio = −8.8, *P* = 3.1 × 10^−5^, Multinomial Goodness-Of-Fit test).

There were two centromeres in common between the unique growth and unique paclitaxel loci, on chromosomes 11 and X. This overlap was insignificant likely because of the low numbers (odds ratio = 7.4, *P* = 0.12, Fisher’s Exact Test). There was no significant difference between the −log_10_ *P* values of the unique paclitaxel centromeres (−log_10_ *P* = 26.3 ± 5.6) and unique growth centromeres (−log_10_ *P* = 19.3 ± 1.3; t[1,3.3] = 1.2, *P* = 0.31, Welch Two Sample t-test).

#### Coding and non-coding paclitaxel loci

The nearest genes were coding for 19 (56%) and non-coding for 15 (44%) of the 34 non-centromeric unique paclitaxel loci. In contrast to the RH growth loci, which were enriched in coding genes, there was no significant difference in the numbers of unique coding and non-coding paclitaxel genes compared to randomly chosen locations in the genome (*χ*^2^[1] = 1.9*, P* = 0.17) (**Supporting Information**). Although the RH coding and non-coding growth genes had similar −log_10_ *P* values, the unique coding region paclitaxel loci had significantly lower −log_10_ *P* values (14.3±1.5) than the non-coding (19.5±1.9; t[1,28.5] = 2.1, *P* = 0.040, Welch Two Sample t-test). Together, our observations suggest that coding and non-coding genes make important contributions to the action of paclitaxel as well as growth.

#### Chromosomal distribution of paclitaxel loci

The number of unique paclitaxel loci compared to the number of genes on each chromosome is shown in **Figure S21B**. The distribution of loci was not significantly different from uniform (*χ*^2^[22] = 22.8*, P* = 0.4), although post-hoc testing revealed one significant result, a surfeit on chromosome 7 (*χ*^2^[1] = 6.7*, P* = 9.7 × 10^−3^).

#### Novel paclitaxel loci

Of the 19 unique coding region RH paclitaxel genes, eight had neither GeneRIF nor Reactome entries. Of the eight, AC020915.5 also lacked entries in the PubMed database. None of the 15 unique non-coding non-centromeric RH paclitaxel genes had entries in either GeneRIF or Reactome. Study of these novel genes may provide new insights into paclitaxel action.

### Further details on interaction loci

#### Interaction loci tally

The majority of the 62 interaction loci (60 out of 62) had negative coefficients (−5.2 × 10^−3^ ± 1.7 × 10^−4^; coefficients in all non-overlapping 1 Mb windows, 8.9 × 10^−4^ ± 3.5 × 10^−5^), while the other two loci (LINC01487 and AC111188.1) had positive coefficients (5.4 × 10^−3^ ± 3.4 × 10^−4^) (**Figures S23C, S30 and S33, Tables S10, S11 and S20**).

#### Overlap of interaction loci with growth and paclitaxel loci

The interaction −log_10_ *P* values showed significantly higher correlation with paclitaxel (*R* = 0.60 ± 0.14) than growth (*R* = 0.03±0.14, non-overlapping 1 Mb windows, t[1,8.9] = 2.8, *P* = 0.02, Welch Two Sample t-test) (**Figure S32A, Table S14**). Consistent with this observation, the correlation of the interaction and growth −log_10_ *P* values at 75 nM paclitaxel (*R* = 0.60) was significantly higher than at the other paclitaxel concentrations (*R* = 0.11 0.03, non-overlapping 1 Mb windows, t[1,3] = 9.1, *P* = 2.8 × 10^−3^, Two Sample t-test with pooled variance).

There were 15 genes that overlapped between the interaction loci and the 859 unique growth loci (odds ratio = 22.6*, P* = 8.3 × 10^−15^, Fisher’s Exact Test) and 14 genes that overlapped with the 38 unique paclitaxel loci (odds ratio = 731*, P <* 2.2 × 10^−16^, Fisher’s Exact Test) (**Figure S31**). As expected, the overlap between the interaction loci and the unique growth loci was greatest at the highest paclitaxel concentration, 75 nM, (odds ratio = 586, *P <* 2.2 × 10^−16^, Fisher’s Exact Test), while the overlap between the interaction loci and the unique paclitaxel loci was greatest at the highest growth time, 6 weeks (odds ratio = 1522, *P <* 2.2 × 10^−16^, Fisher’s Exact Test).

A total of 22 interaction loci were shared with either the unique growth or unique pacliatexel loci (odds ratio = 38, *P <* 2.2 × 10^−16^, Fisher’s Exact Test), while a total of 7 interaction loci were shared with both (odds ratio = 2447, *P <* 2.2 × 10^−16^, Fisher’s Exact Test) (**Figure S31**).

Similar overlap results were obtained for the interaction and non-unique loci. The highest overlap between the interaction and non-unique growth loci was at 75 nM paclitaxel (odds ratio = 482, *P <* 2.2 × 10^−16^, Fisher’s Exact Test) and the highest overlap between the interaction and non-unique paclitaxel loci was at 6 weeks of growth (odds ratio = 1522, *P <* 2.2 × 10^−16^, Fisher’s Exact Test) (**Figure S32B, Table S15**).

#### Interaction centromeres

Five interaction loci mapped to centromeres on chromosomes 11, 15, 16, 20 and X. There was no significant difference in −log_10_ *P* values between the centromeric (−log_10_ *P* = 16.2 ± 2.2) and non-centromeric interaction loci (−log_10_ *P* = 12.0 ± 0.5; t[1,4.3] = 1.9, *P* = 0.13, Welch Two Sample t-test).

Three of the five interaction centromeres overlapped with the four unique growth centromeres (chromosomes 11, 15 and X; odds ratio = 20, *P* = 0.021, Fisher’s Exact Test) and four with the four unique paclitaxel centromeres (chromosomes 11, 16, 20 and X; odds ratio =, *P* = 5.6 × 10^−4^,

Fisher’s Exact Test). There was no significant difference between the −log_10_ *P* values of the interaction centromeres (16.2 ± 2.2) and either the unique growth centromeres (19.3 ± 1.3, t[1,6.2] = 1.2, *P* = 0.26, Welch Two Sample t-test) or the unique paclitaxel centromeres (26.3 ± 5.6, t[1,3.9] = 1.7, *P* = 0.17, Welch Two Sample t-test).

#### Coding and non-coding interaction loci

The nearest genes were coding for 26 (46%) and non-coding for 31 (54%) of the 57 non-centromeric interaction loci. There was no significant difference in the proportion of coding and non-coding interaction genes compared to randomly chosen locations in the genome (*χ*^2^[1] = 0.05*, P* = 0.83) (**Supporting Information**). In addition, there were no significant differences in the −log_10_ *P* values between the coding (−log_10_ *P* = 12.4 ± 0.9) and non-coding interaction loci (−log_10_ *P* = 11.6 ± 0.4; t[1,36.5] = 0.8, *P* = 0.4, Welch Two Sample t-test). Together, our observations suggest that coding and non-coding genes make important contributions to the interaction between growth and paclitaxel.

#### Chromosomal distribution of interaction loci

The number of interaction loci compared to the number of genes on each chromosome is shown in **Figure S21C**. The distribution of loci was significantly different from uniform (*χ*^2^[22] = 50.9*, P* = 4.4 × 10^−4^), with significant surfeits on chromosomes 6 (*χ*^2^[1] = 18.2*, P* = 1.9 × 10^−5^) and 13 (*χ*^2^[1] = 5.5*, P* = 0.02).

#### Novel interaction loci

Of the 25 coding region RH interaction genes, five had neither GeneRIF nor Reactome entries. However, all of these five genes had entries in the PubMed database. None of the 31 non-coding non-centromeric RH interaction genes had entries in either GeneRIF or Reactome.

### Public data

The human gene catalog was the comprehensive annotation set of GENCODE v31 (GRCh38.p12) (Frankish et al. 2019). Loss-of-function growth genes identified in CRISPR screens (Hart et al. 2015; Wang, Birsoy, et al. 2015) and microarray expression data from the G3 human RH panel (Wang, Ahn, et al. 2011) were from published studies. Human tissue RNA-Seq data were retrieved from GTEx v8 (GTEx Consortium 2015).

Paralogs were identified using the duplicated genes database (DGD), employing Ensembl release 71 (Ouedraogo et al. 2012). The degree of intolerance to predicted loss-of-function (pLoF) variation in human genes was measured using the observed/expected ratio from the Genome Aggregation Database (gnomAD) release 2.1.1. (Karczewski et al. 2019). The ratio of nonsynonymous to synonymous substitutions (dN/dS) was calculated using mouse-human homologs in Ensembl release 97 (Cunningham et al. 2019).

Homologene release 68 was employed to evaluate the number of species with gene orthologs (Sayers et al. 2019). Protein-protein interactions were identified using STRING v11 (Szklarczyk et al. 2019).

Functional analysis of RH genes was performed using DAVID v6.8 (Huang et al. 2009). Enrichment in multisubunit protein complexes was assessed using the GO_CC_ALL category of DAVID, as described (Hart et al. 2015). GO analyses used the 2019-07-03 release and the PANTHER Overrepresentation Test (Mi et al. 2019; The Gene Ontology Consortium 2019).

InterPro was employed to assess protein domains (Huang et al. 2009; Mitchell et al. 2019). Coding genes of unknown function were classified as those lacking entries in GeneRIF (Jimeno-Yepes et al. 2013) and the Reactome Knowledgebase v70 (Fabregat et al. 2018).

### Further details on mapping resolution

#### Mapping resolution using TK1

A Gaussian linear mixed model was used to evaluate the distance between the location of the TK1 gene and its retention peak in all the RH samples. The model employed fixed and interaction effects of growth time and paclitaxel concentration, combined with random intercepts of RH pool and nested drug history (**Supporting Information**).

There were no significant fixed effects of growth (t[1,92.5] = 0.2*, P* = 0.9, Kenward-Roger df; average conditional effect) or drug concentration (t[1,25.6] = 1.8*, P* = 0.08, Kenward-Roger df; average conditional effect), but the interaction was significant (t[1,94.7] = 2.7*, P* = 9.4 × 10^−3^, Kenward-Roger df). These observations indicate that, as expected, growth time or drug concentration had minimal effects on TK1 localization. There was, however, a significant random effect of pool (*χ*^2^[1] = 7.7*, P* = 5.4 × 10^−3^, ICC = 0.31, likelihood ratio test) but not drug history (*χ*^2^[1] = 3.0*, P* = 0.09, ICC = 0.12, likelihood ratio test), consistent with the observed batch effect of RH pool (**Figure S11**).

The intercept distance between the peak retention of TK1 and the location of the gene was estimated as −27 kb ± 66 kb using the mixed model. This distance was statistically indistinguishable from zero (t[1,5.1] = 0.4*, P* = 0.7, Kenward-Roger df; average conditional effect) (**Figures S34A and S34C**).

#### Mapping resolution using centromeres

As an independent approach to evaluating mapping accuracy, we measured the distances between the observed retention peak for each centromere and their assigned positions in the genome (**Figures 2D–2F, S5, S6, S34B, S34D and S34E**). The functional locations of centromeres are poorly defined, and generous limits are provided on their positions in the human genome build GRCh38/hg38 (https://genome.ucsc.edu). For this analysis, we took the consensus locations of the centromeres to be the center of the contigs covering each centromere in hg38.

Some centromeres showed clearly defined retention peaks, while others showed a plateau, suggesting uniformly dispersed functional attributes (**Figures 2E, 2F and S6**). Nevertheless, we reasoned that certain centromere sequences will have greater functional importance than others, indicated by the maximum retention peak. To evaluate the distances between the observed retention peak for each centromere and their consensus assignations in the genome, we used the mixed effects model described for TK1 above (**Supporting Information**), with the addition of a random intercept of chromosome.

Similar to TK1, the fixed effects of growth (t[1,2293] = 0.7*, P* = 0.5, Kenward-Roger df; average conditional effect), paclitaxel (t[1,27.6] = 0.06*, P* = 0.95, Kenward-Roger df; average conditional effect) and their interaction (t[1,2053] = 0.05*, P* = 0.96, Kenward-Roger df) had no significant effect on centromere mapping accuracy. Also as for TK1, and consistent with the observed RH pool batch effects (**Figure S11**), the random effect of pool was significant (*χ*^2^[1] = 4.5*, P* = 0.03, likelihood ratio test), but the nested random effect of drug history was not (*χ*^2^[1] = 0.09*, P* = 0.76, likelihood ratio test). Nevertheless, the variance due to the RH pools and drug history was minor, accounting for only 0.50% and 0.10% of the variance components, respectively (ICC = 5.0 × 10^−3^, 1.0 × 10^−3^).

In contrast, the between-chromosome variance dominated the between-RH pool effect, accounting for 37% of the variance components (ICC = 0.37). Further, the random effect of chromosome (*χ*^2^[1] = 1077*, P <* 2.2 × 10^−16^, likelihood ratio test) was much more significant than that of the RH pools (**Figure S34B**). The significant chromosome random effect suggests that there are substantial differences between the functional locations of centromeres and their currently assigned positions. Our approach may allow more accurate localization of active centromere sequences.

The estimated distance between the centromere retention peaks and their assigned genomic location was statistically indistinguishable from zero (mean distance = −30 kb ± 105 kb; t[1,24] = 0.3*, P* = 0.8, Kenward-Roger df; average conditional effect) (**Figures S34D–S34E**). The mapping accuracy using centromeres was similar to that obtained using TK1, together suggesting a resolution ~30 kb. Despite the good mapping resolution of RH-BSA, regions of high gene density were not always clearly resolved (**Figure S35**). Higher radiation doses and pool numbers can help resolve such regions in the future.

### Further details on mitochondrial copy number

#### Calculating mitochondrial copy number of HEK293 and A23 cells

Mitochondrial copy numbers in the HEK293 and A23 cells were calculated by comparing the number of mitochondrial sequence reads with those of the host nuclear genome, normalized to the mitochondrial and nuclear genome lengths, respectively (**Table S4**). For HEK293 cells an additional correction factor of three was employed, as these cells are thought to be pseudotriploid (Lin et al. 2014), while mitochondria are haploid. For A23 cells the correction factor was two.

#### Calculating mitochondrial copy number of RH samples

For the RH samples, including the RH pools (**Table S21**), the human and hamster mitochondrial copy numbers were deduced by comparing the reads from the mitochondria of each species with the reads from the hamster nuclear genome, normalized to the lengths of the respective genomes. A further correction factor of two was employed to account for the haploid and diploid nature of the mitochondrial and hamster genomes, respectively.

Thus, the human mitochondrial copy number, 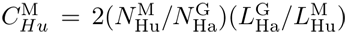, where 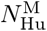 is the is the number of hamster-specific genome reads, 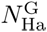 is the length of the hamster genome (2 368 906 908 bp) and 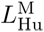 is the length of the human mitochondrial genome (16 569 bp). The hamster mitochondrial copy number, 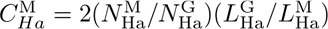, where 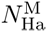 is the number of hamster-specific mitochondrial reads and 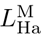 is the length of the hamster mitochondrial genome (16 283 bp).

Human mitochondrial copy numbers were also estimated by comparing human mitochondrial reads with those from the 1 Mb window giving maximum human TK1 reads, assuming that the TK1 gene had a copy number of 1. Normalization employed the respective lengths of the human mitochondria and the 1 Mb TK1 window. In addition, a correction factor for the reversion frequency was applied. Thus, 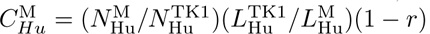, where 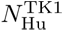 is the maximum number of human-specific reads in the 1 Mb window encompassing TK1, 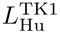 is the length of the TK1 window (1 × 10^6^ bp) and *r* is the reversion frequency (**Table S1**).

#### Hamster mitochondrial copy number changes in the RH cells

We evaluated changes of the hamster mitochondrial copy number in the RH samples using the previously described Gaussian linear mixed model with fixed effects of growth time, paclitaxel concentration and their interaction, combined with random effects of RH pool and drug history (**Supporting Information**).

Hamster mitochondrial copy numbers showed a significant decrease as a result of growth (t[1,93.5] = 2.3*, P* = 0.02, Kenward-Roger df; average conditional effect) (**Figure S36A**) and a significant increase as a result of paclitaxel (t[1,28.1] = 3.4*, P* = 2 × 10^−3^, Kenward-Roger df; average conditional effect) (**Figure S36B**). These changes are consistent with previous observations, which showed a decrease in mitochondria as a result of malignant growth (Reznik et al. 2016) and an increase in cells treated with paclitaxel (Karbowski et al. 2001). The interaction was insignificant (t[1,95.5] = 1.5*, P* = 0.1). In addition, there was a significant random effect of pool (*χ*^2^[1] = 6.6*, P* = 0.01, ICC = 0.18, likelihood ratio test), but not drug history (*χ*^2^[1] = 0*, P* = 1, ICC = 0, likelihood ratio test).

#### Details on human loci regulating hamster mitochondrial copy number

To identify human loci that regulated the levels of hamster mitochondria, we amended our negative binomial mixed model by adding hamster mitochondrial reads as a supplementary fixed effect (**Figures S37A–S37D, Tables S22 and S23, Materials and Methods**). The addition of hamster mitochondrial abundance to the model resulted in only minor changes to the −log_10_ *P* values for growth, paclitaxel and their interaction (weakest correlation, paclitaxel week 1, *R* = 0.84, t[1,3052] = 87, *P <* 2.2 × 10^−16^; mean *R* = 0.96 ± 0.02; non-overlapping 1 Mb windows).

#### Details on human loci regulating human mitochondrial copy number

Human mitochondrial copy numbers in the RH samples were calculated either by reference to the hamster nuclear genome or the human TK1 gene. The two methods gave comparable results.

Using reads from the hamster nuclear genome as a reference gave a copy number of 1.4 ± 0.8 for human mitochondria in the six RH pools, not significantly different from zero (t[1,5] = 1.7, *P* = 0.14; One Sample t-test) (**Figures S36C and S36D, Table S21**). Using TK1 as a reference gave a copy number of 2.8 ± 1.6 in the RH pools, also not significantly different from zero (t[1,5] = 1.8, *P* = 0.14; One Sample t-test) (**Figures S36E and S36F, Table S21**).

In addition, the mean of the two methods gave a copy number of 2.1 ± 1.2 in the RH pools, which was not significantly different from zero (t[1,5] = 1.8, *P* = 0.14; One Sample t-test) (**Figures S36G and S36H, Table S21**).

We used the previously described Gaussian linear mixed model with fixed effects of growth time, paclitaxel concentration and their interaction, combined with random effects of RH pool and drug history to evaluate the human mitochondrial copy number in all the RH samples (**Supporting Information**). The results were of varying and marginal significance, depending on the measurement method used.

The copy number was 0.6±0.3 using the hamster nuclear genome as a reference (t[1,5.2] = 2.1*, P* = 0.09, Kenward-Roger df; average conditional effect), 1.4 ± 0.5 using TK1 as a reference (t[1,5.4] = 2.7*, P* = 0.04, Kenward-Roger df; average conditional effect), and 1.0 ± 0.4 using the mean of the two methods (t[1,5.3] = 2.7*, P* = 0.04, Kenward-Roger df; average conditional effect).

There was no significant effect on human mitochondrial copy number of growth (t[1,93.5] = 0.1*, P* = 0.9, hamster alignments; t[1,93.4] = 0.8*, P* = 0.4, TK1; t[1,93.5] = 0.7*, P* = 0.5, mean of two methods; Kenward-Roger df and average conditional effects), paclitaxel (t[1,28.1] = 0.4*, P* = 0.7, hamster alignments; t[1,27.2] = 1.8*, P* = 0.09, TK1; t[1,28.2] = 1.5*, P* = 0.2, mean of two methods; Kenward-Roger df and average conditional effects) or their interaction (t[1,95.5] = 1.6*, P* = 0.1, hamster alignments; t[1,95.5] = 0.9*, P* = 0.4, TK1; t[1,95.5] = 1.2*, P* = 0.2, mean of two methods; Kenward-Roger df).

The random effects of pool were inconclusive (*χ*^2^[1] = 10.8*, P* = 9.9 × 10^−4^, ICC = 0.2, hamster alignments; *χ*^2^[1] = 1.1*, P* = 0.3, ICC = 0.07, TK1; *χ*^2^[1] = 3.3*, P* = 0.07, ICC = 0.1, mean of two methods; likelihood ratio tests), and drug history was insignificant (*χ*^2^[1] = 0*, P* = 1, ICC = 0, hamster alignments; *χ*^2^[1] = 0.2*, P* = 0.6, ICC = 0.05, TK1; *χ*^2^[1] = 0*, P* = 1, ICC = 4.9 × 10^−9^, mean of two methods; likelihood ratio tests).

Despite the low levels of human mitochondria, we nevertheless identified human loci that regulated the copy number of these organelles with FDR *<* 0.05 by including the mitochondrial sequence reads as an additional fixed effect in our negative binomial mixed model (**Figures S37E and S37F, Tables S24 and S25, Materials and Methods**). The additional fixed effect left the −log_10_ *P* values for growth, paclitaxel and their interaction essentially unchanged from those obtained previously (weakest correlation, paclitaxel week 1, *R* = 0.99, t[1,3052] = 538, *P <* 2.2 × 10^−16^; mean *R* = 0.998 ± 4.9 × 10^−4^; non-overlapping 1 Mb windows).

### Hamster genome changes

We evaluated whether hamster copy number alterations (CNAs) (**Figure 2A**) confer selective advantages for growth and paclitaxel using the same statistical model we employed for the human genome (**Tables S26 and S27, Materials and Methods**). Loci that exceeded genome-wide permutation significance thresholds were identified for growth, paclitaxel and their interaction (**Figures S38 and S39**).

Despite the high quality of the hamster genome, a total of 341 contigs, many relatively short, still remained after discarding unanchored contigs (Rupp et al. 2018). For greater clarity and continuity in the hamster genome scans, we removed the 209 shortest remaining contigs, 61% of the contigs (maximum length 3.6 Mb), leaving 132 contigs that encompassed 95% of the hamster genome. However, analyses of sequence alignment numbers in the hamster genome used all available contigs.

The greater abundance of mapped hamster sequence reads compared to human (59±14-fold, RH pools; 84 ± 16-fold, all samples; c.f. **Tables S5 and S6** for RH pools), provided higher statistical power to detect loci. However, the −log_10_ *P* values for significant hamster growth loci (9.2 ± 0.1) were significantly less than for human (21.9 ± 0.2; t[1,3237] = 68, *P <* 2.2 × 10^−16^, Welch Two Sample t-test; non-overlapping 1 Mb windows). A similar picture was evident for paclitaxel loci (−log_10_ *P* = 11.3 ± 0.2, hamster; 20.7 ± 1.0, human; t[1, 46] = 9.6, *P* = 1.7 × 10^−12^, Welch Two Sample t-test; non-overlapping 1 Mb windows). The weaker −log_10_ *P* values of hamster loci are consistent with the greater stability of the hamster genome compared to human (**Figures 3 and S10**).

The mapping resolution using the hamster genome depends on the length of the CNAs, typically encompassing many megabases and genes. Further, the hamster genome is of high quality but is still divided into 132 contigs. Hamster CNAs are thus artefactually divided into smaller intervals. Nevertheless, examples of hamster growth and paclitaxel loci due to pre-existing CNAs are shown in **Figure S40**. In contrast, CNAs in the donor HEK293 genome affect retention frequency but do not affect mapping resolution, which depends on the number of breakpoints caused by the radiation.

### Further details on mycoplasma contamination

#### Quantitation of mycoplasma copy number

Mycoplasma copy numbers were calculated by comparing the number of mycoplasma reads with those from the host nuclear genome in each sample. For HEK293 cells the reference genome was human; for A23 cells and the RH samples the reference genome was hamster. Normalization was used to adjust for the respective lengths of each nuclear and mycoplasma genome. A further correction factor of two accounted for the diploid and haploid nature of the nuclear and mycoplasma genomes, respectively.

Thus, the mycoplasma copy number, 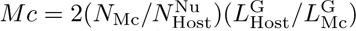, where *N*_Mc_ is the number of sequence reads aligned to the mycoplasma genome, 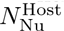 is the number of reads aligned to the host nuclear genome, 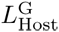 is the length of the host nuclear genome and 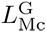 mycoplasma genome.

#### Effects of mycoplasma contamination

We examined the sequence data from the parental HEK293 and A23 cell lines to see if the cells were contaminated with mycoplasma. We aligned reads to the five species shown in **Table S28**, with 1 bp and 4 bp mismatches giving identical read numbers.

A recent paper using publicly available RNA-Seq datasets evaluated the prevalence of mycoplasma contamination in published studies (Olarerin-George and Hogenesch 2015). Of the datasets, 11% featured one or more contaminated samples, defined as ≥ 100 reads million*^−^*^1^ mapping to mycoplasma. None of our parental samples exceeded these levels (**Table S28**). However, a lower threshold is appropriate for our dataset, since mammalian genome sequences are more complex than RNA-Seq datasets.

The HEK293 cells were nearly devoid of mycoplasma sequences. The A23 cells, however, had appreciable levels. The most abundant species was *M. fermentans* at 0.26 copies per cell, followed by *M. hominis* and *M. hyorhinis*.

To assess the effects of mycoplasma contamination in the RH samples, reads were aligned to *M. fermentans* using 1 bp mismatches. The *M. fermentans* copy number was 0.11 ± 0.03 in the six RH pools (t[1,5] = 3.4, *P* = 0.02; One Sample t-test). A linear mixed model was used to evaluate *M. fermentans* copy number in all RH samples using fixed effects of growth and paclitaxel as well as their interaction, together with pool and nested drug history as random effects (**Supporting Information**). The *M. fermentans* copy number was 0.13 ± 0.03 (t[1,5.16] = 4.9*, P* = 4.1 × 10^−3^, Kenward-Roger df; average conditional effect).

The mixed model revealed no significant effect of growth (t[1,91.5] = 1.1*, P* = 0.3, Kenward-Roger df; average conditional effect), paclitaxel (t[1,24.7] = 0.8*, P* = 0.4, Kenward-Roger df; average conditional effect) or their interaction (t[1,93.6] = 1.0*, P* = 0.3, Kenward-Roger df). However, there was a significant random effect of pool (*χ*^2^[1] = 3.9*, P* = 0.047, ICC = 0.22, likelihood ratio test), and drug history (*χ*^2^[1] = 13.0*, P* = 3.2 × 10^−4^, ICC = 0.25, likelihood ratio test).

Although mycoplasma contamination is irrelevant to the original role of RH panels in high resolution genetic mapping, these bacteria could nevertheless affect the findings of our study. To evaluate the consequences of mycoplasma contamination, we modified our negative binomial mixed model (**Materials and Methods**) to include the *M. fermentans* copy number as an additional fixed effect (**Tables S29–S31**). The −log_10_ *P* values for growth and paclitaxel remained essentially unchanged (weakest correlation, paclitaxel week 1, *R* = 0.97, t[1, 3052] = 222, *P <* 2.2 × 10^−16^; mean *R* = 0.995 ± 2.4 × 10^−3^; non-overlapping 1 Mb windows). In addition, there were no human loci that regulated the levels of the bacterium and that also exceeded either the permutation significance threshold or the more liberal FDR threshold of 0.05 (in fact, all FDR = 1). These observations indicate that the mycoplasma contamination had minimal effects on our conclusions.

## Supporting Figures

**Figure S1.**
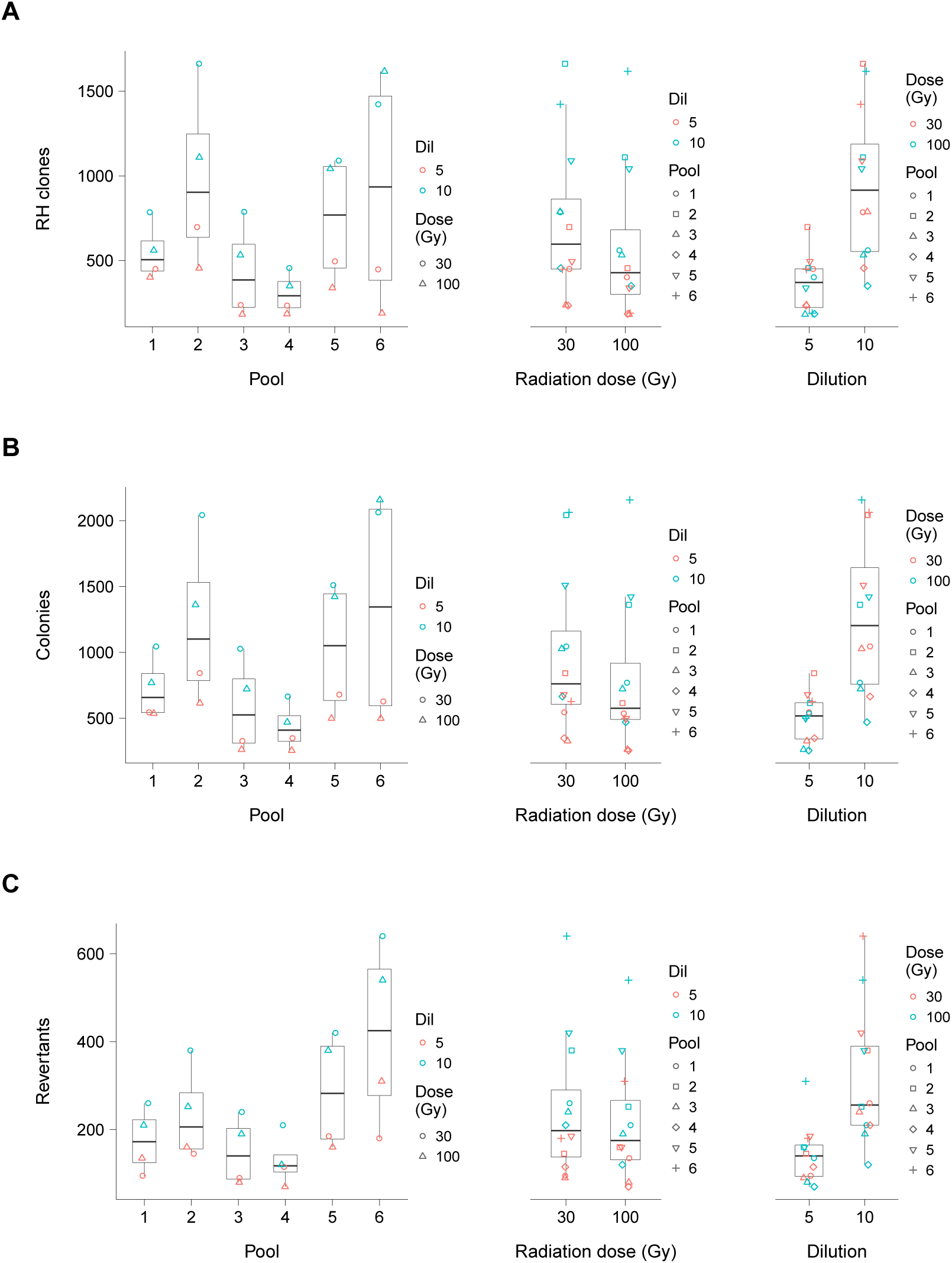
Cell fusions. (**A**) RH clones. (**B**) Total colonies. (**C**) Revertants. Numbers shown.

**Figure S2.**
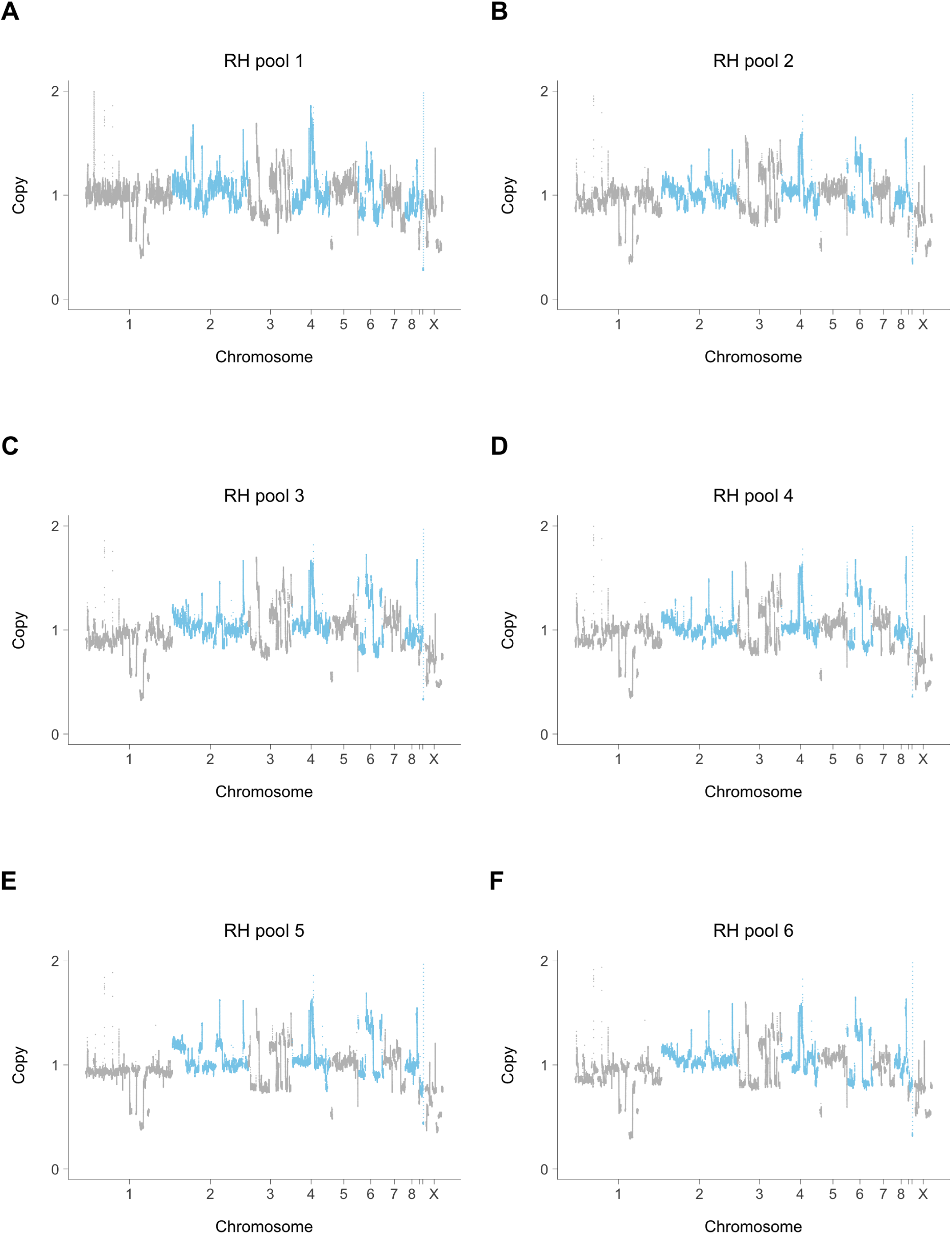
Hamster genome copy number in six RH pools. (**A**) Pool 1. (**B**) Pool 2. (**C**) Pool 3. (**D**) Pool 4. (**E**) Pool 5. (**F**) Pool 6. Sequence coverage normalized to a mean value of 1.

**Figure S3.**
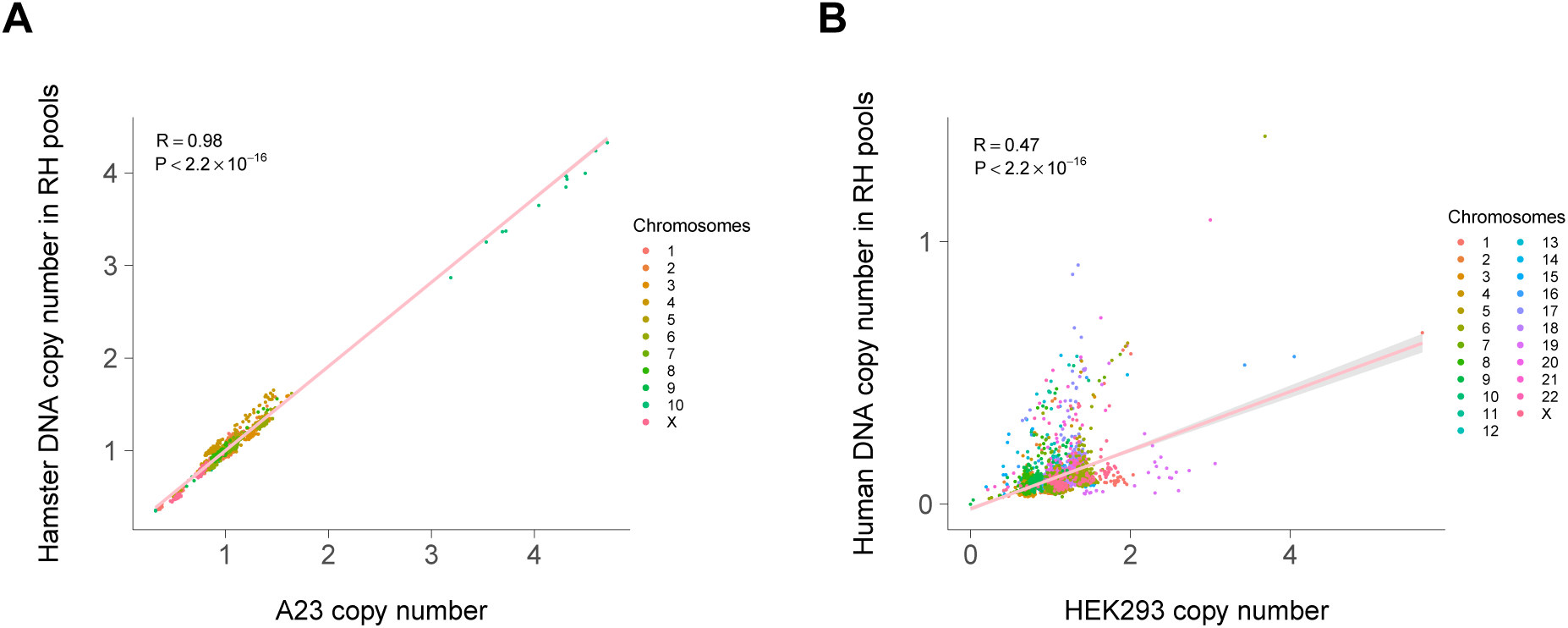
Copy number changes after cell fusion. (**A**) Hamster genome copy number in the six RH pools is little changed from the recipient A23 cells. (**B**) Human genome copy number shows changes as a result of cell fusion but is still correlated with initial copy number in the donor HEK293 cells. Pink lines, linear best fit. Grey bands, 95% confidence interval. Non-overlapping 1 Mb windows.

**Figure S4.**
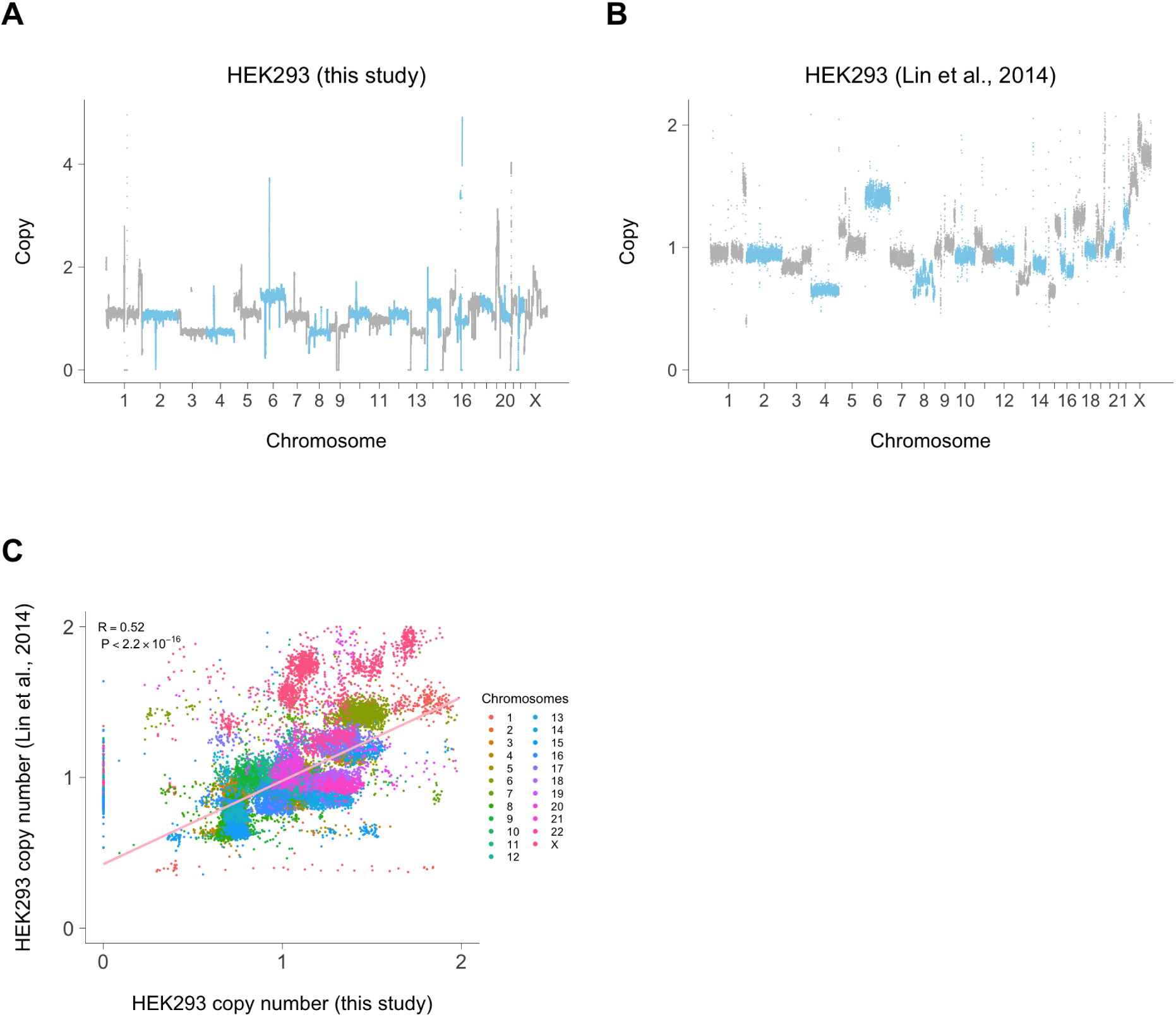
HEK293 CNAs. (**A**) Relative genome copy number in HEK293 cells from this study. (**B**) Relative copy number of HEK293 cells from published genome sequence (Lin et al. 2014). (C) Scatterplot showing relationship of HEK293 genome copy number in the two datasets. All data points from (Lin et al. 2014) used. Pink line, linear best fit. Grey band, 95% confidence interval (too small to be visible).

**Figure S5.**
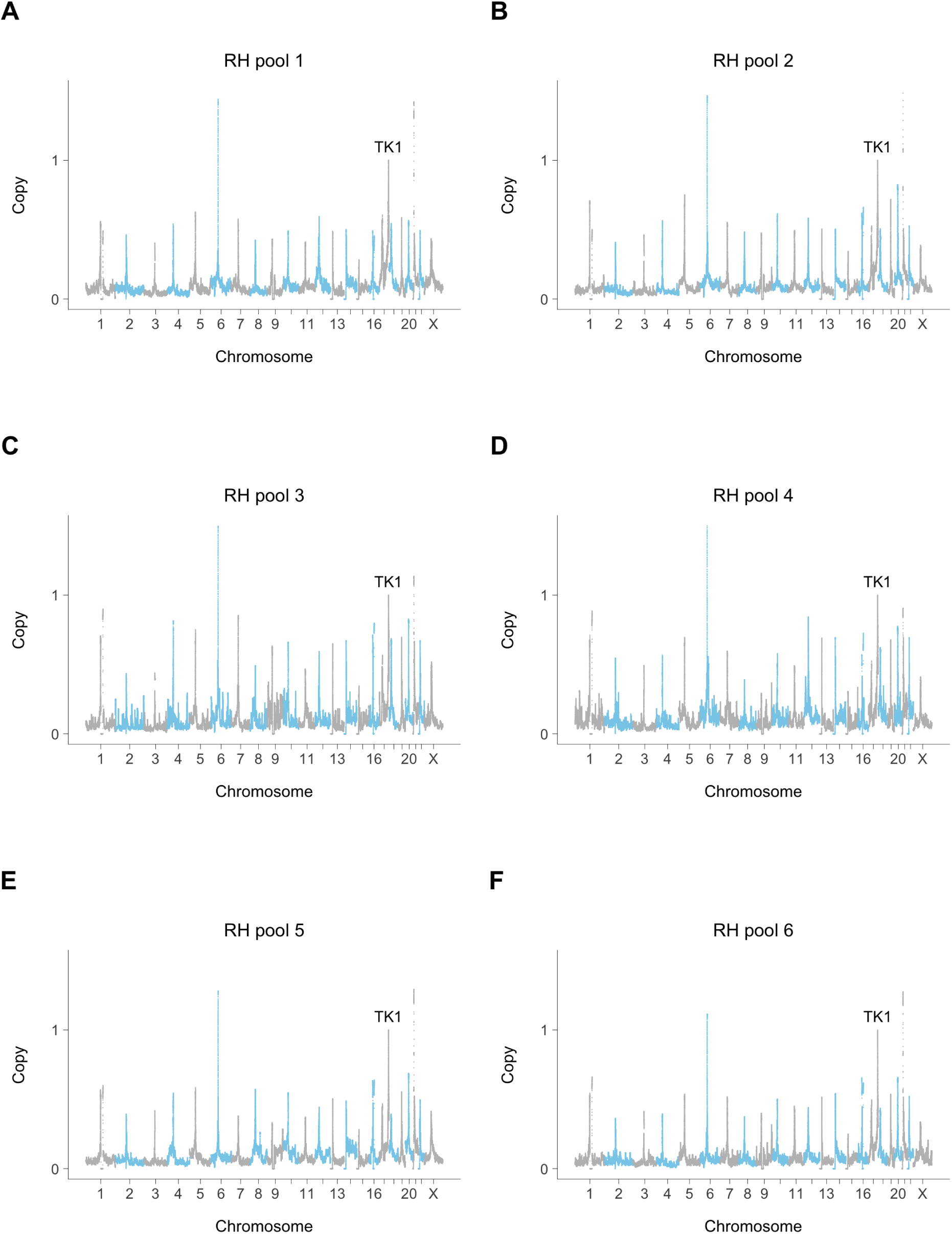
Retention of human DNA in six RH pools. (**A**) Pool 1. (**B**) Pool 2. (**C**) Pool 3. (D) Pool 4. (**E**) Pool 5. (**F**) Pool 6. Sequence coverage normalized to a mean value of 1 at TK1.

**Figure S6.**
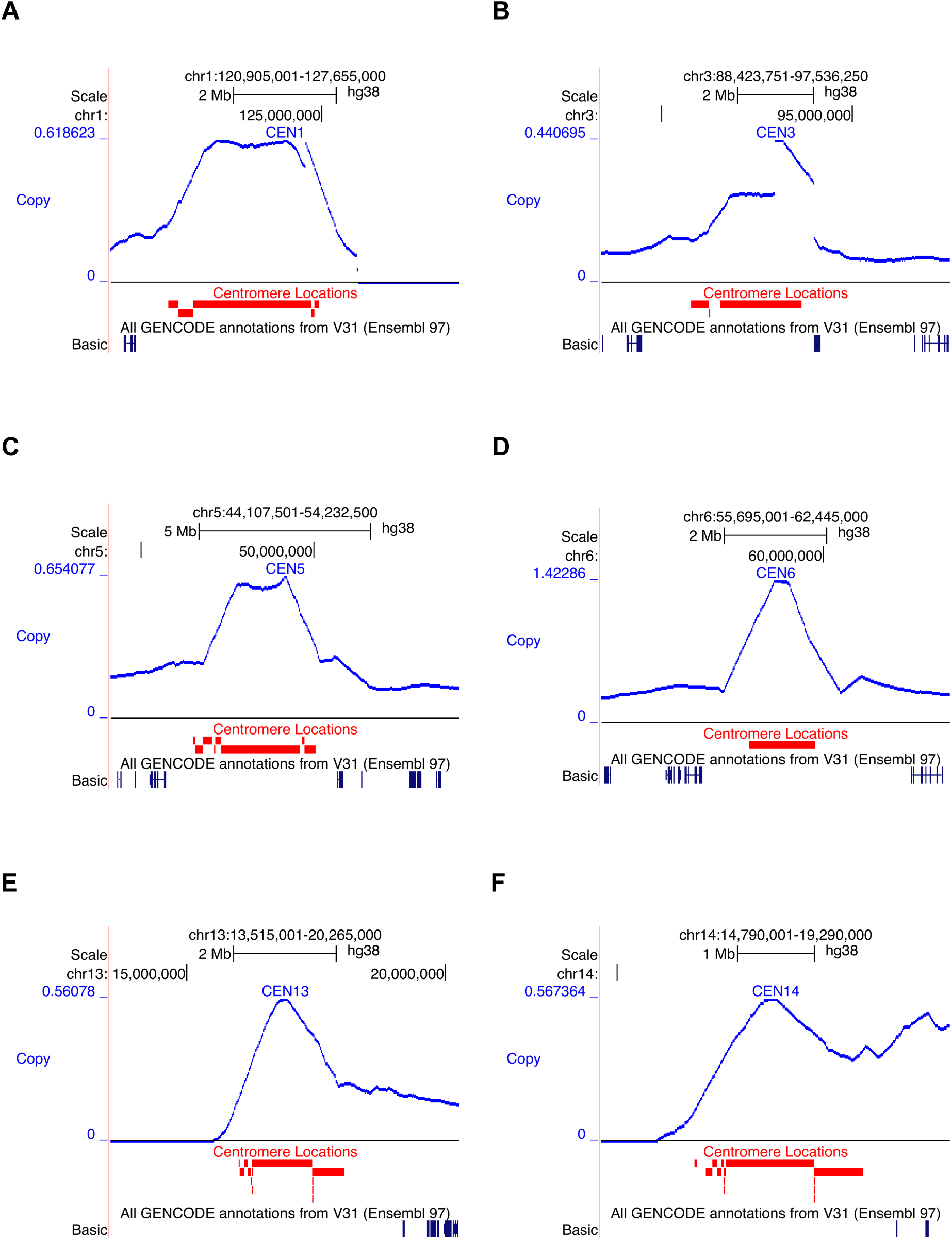
Close up views of human centromere retention in the RH pools. (**A**) Chromosome 1. (**B**) Chromosome 3. (**C**) Chromosome 5. (**D**) Chromosome 6. (**E**) Chromosome 13. (**F**) Chromosome 14. Human DNA retention averaged across the six RH pools. TK1 assigned retention of 1.

**Figure S7.**
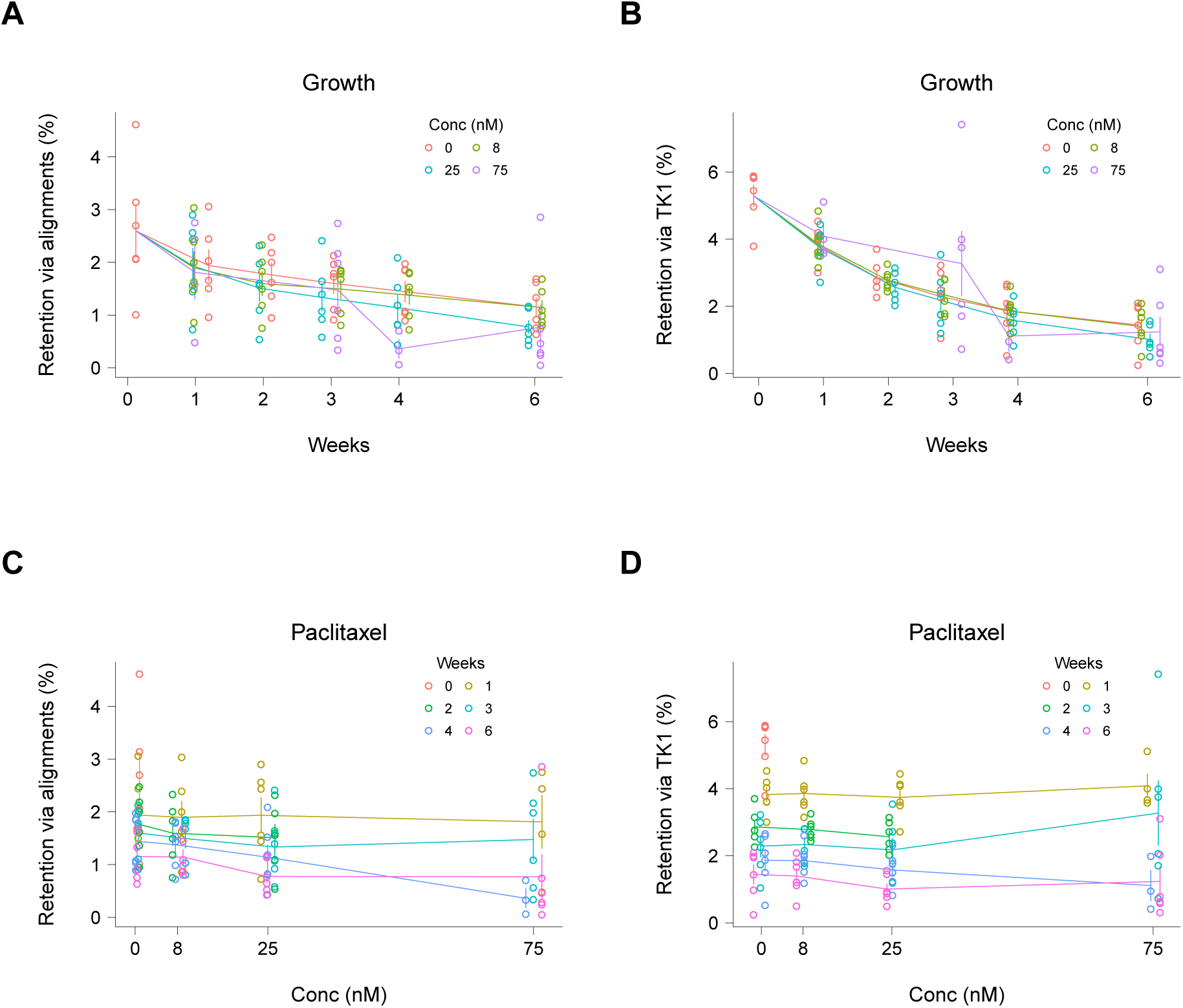
Overall human DNA retention stratified by growth time and paclitaxel concentration. (**A**) Changes in median retention as a result of growth, calculated using aligned sequence reads. (**B**) Retention changes as a result of growth, calculated assuming TK1 retention of 1 and corrected for revertants. (**C**) Retention changes as a result of paclitaxel, calculated using aligned sequence reads. (**D**) Retention changes as a result of paclitaxel, calculated assuming TK1 retention of 1 and corrected for revertants. Means ± s.e.m.

**Figure S8.**
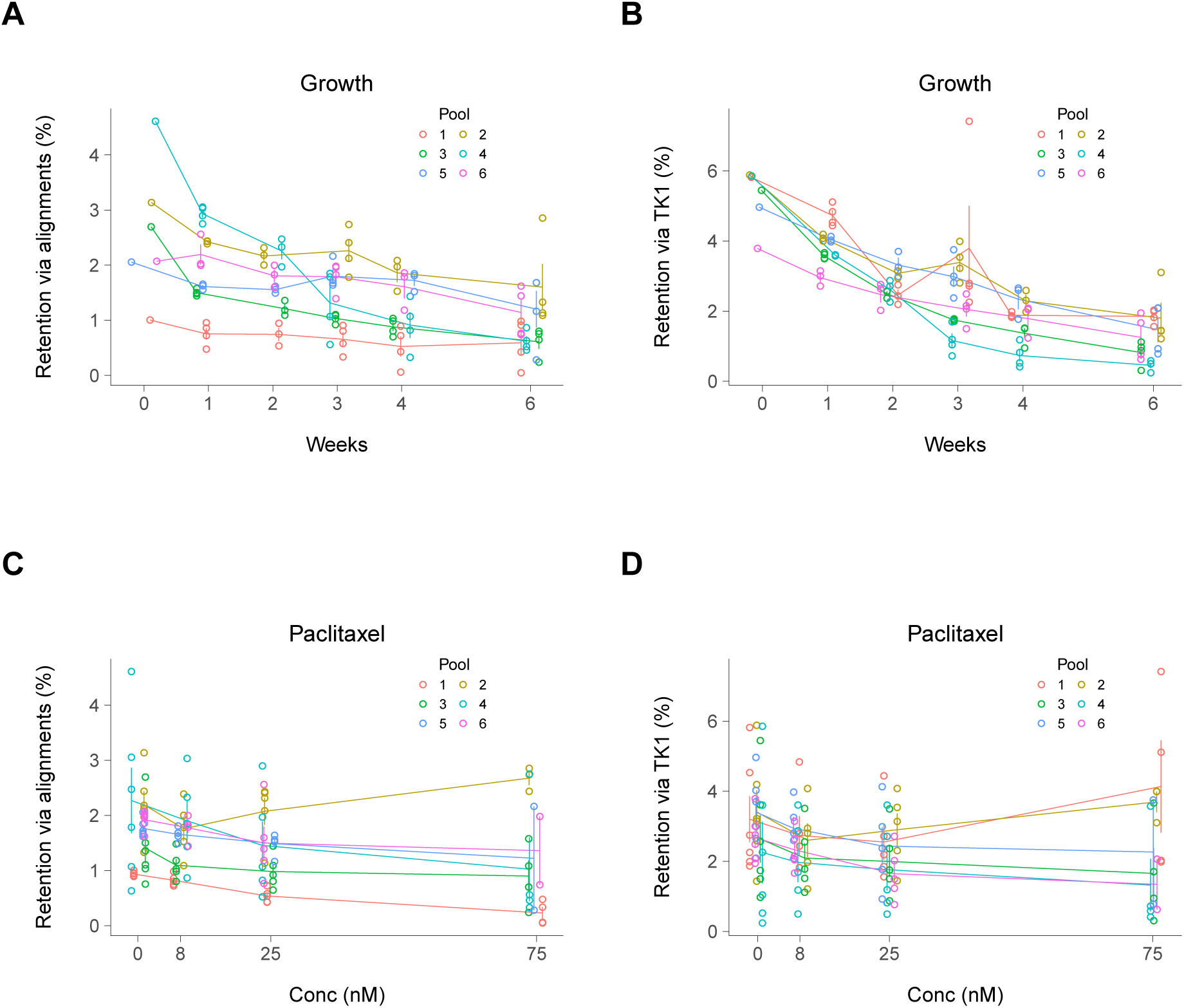
Overall human DNA retention changes stratified by pool. (**A**) Changes in median retention as a result of growth, calculated using aligned sequence reads. (**B**) Retention changes as a result of growth, calculated assuming TK1 retention of 1 and corrected for revertants. (**C**) Retention changes as a result of paclitaxel, calculated using aligned sequence reads. (**D**) Retention changes as a result of paclitaxel, calculated assuming TK1 retention of 1 and corrected for revertants. Means ± s.e.m.

**Figure S9.**
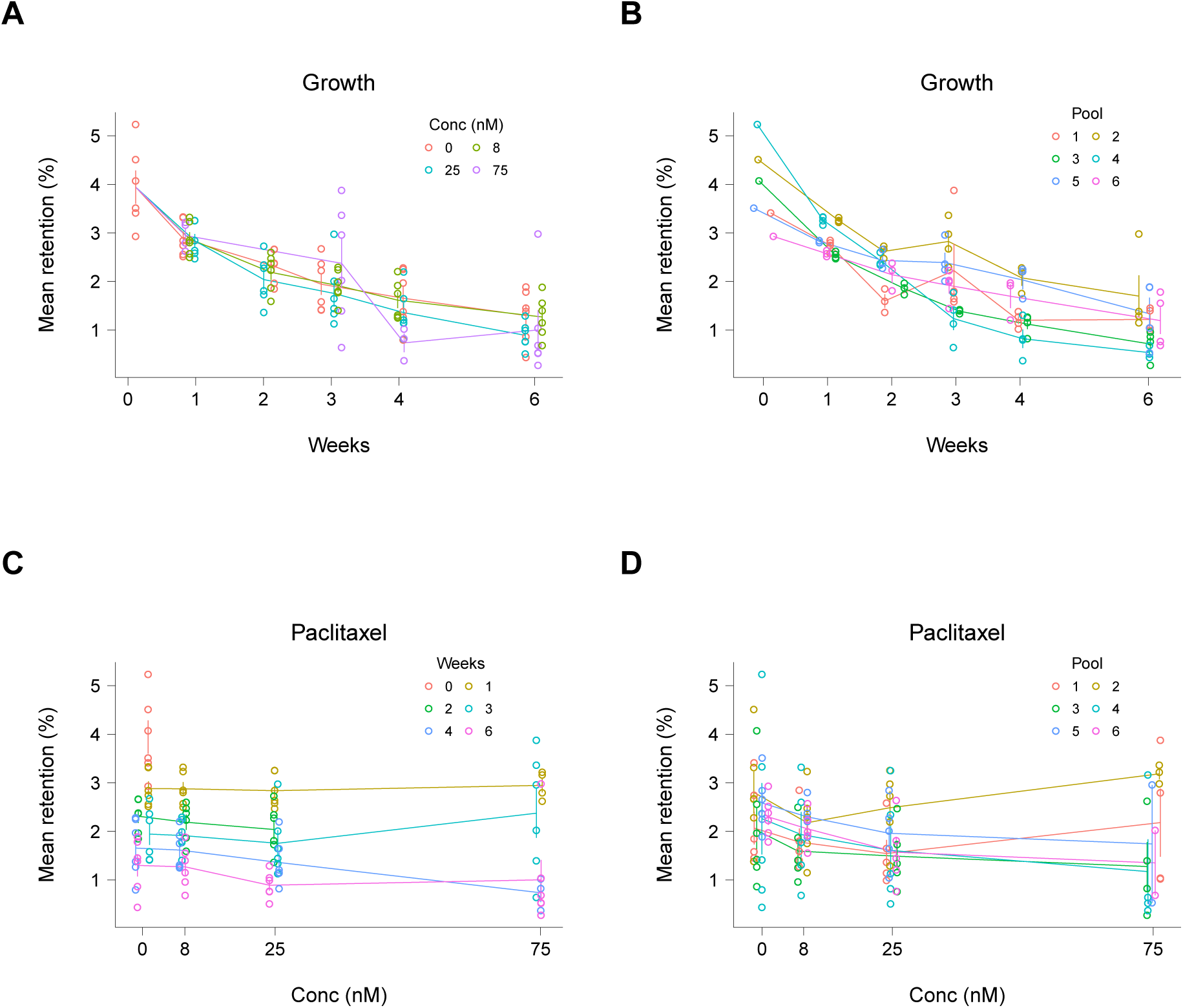
Mean human DNA retention changes. (**A**) Mean retention changes as a result of growth time, stratified by paclitaxel concentration. (**B**) Mean retention changes as a result of growth time, stratified by pool. (**C**) Mean retention changes as a result of paclitaxel concentration, stratified by growth time. (**D**) Mean retention changes as a result of paclitaxel concentration, stratified by pool. Retention based on mean of sequence alignments and TK1 corrected for revertants ± s.e.m.

**Figure S10.**
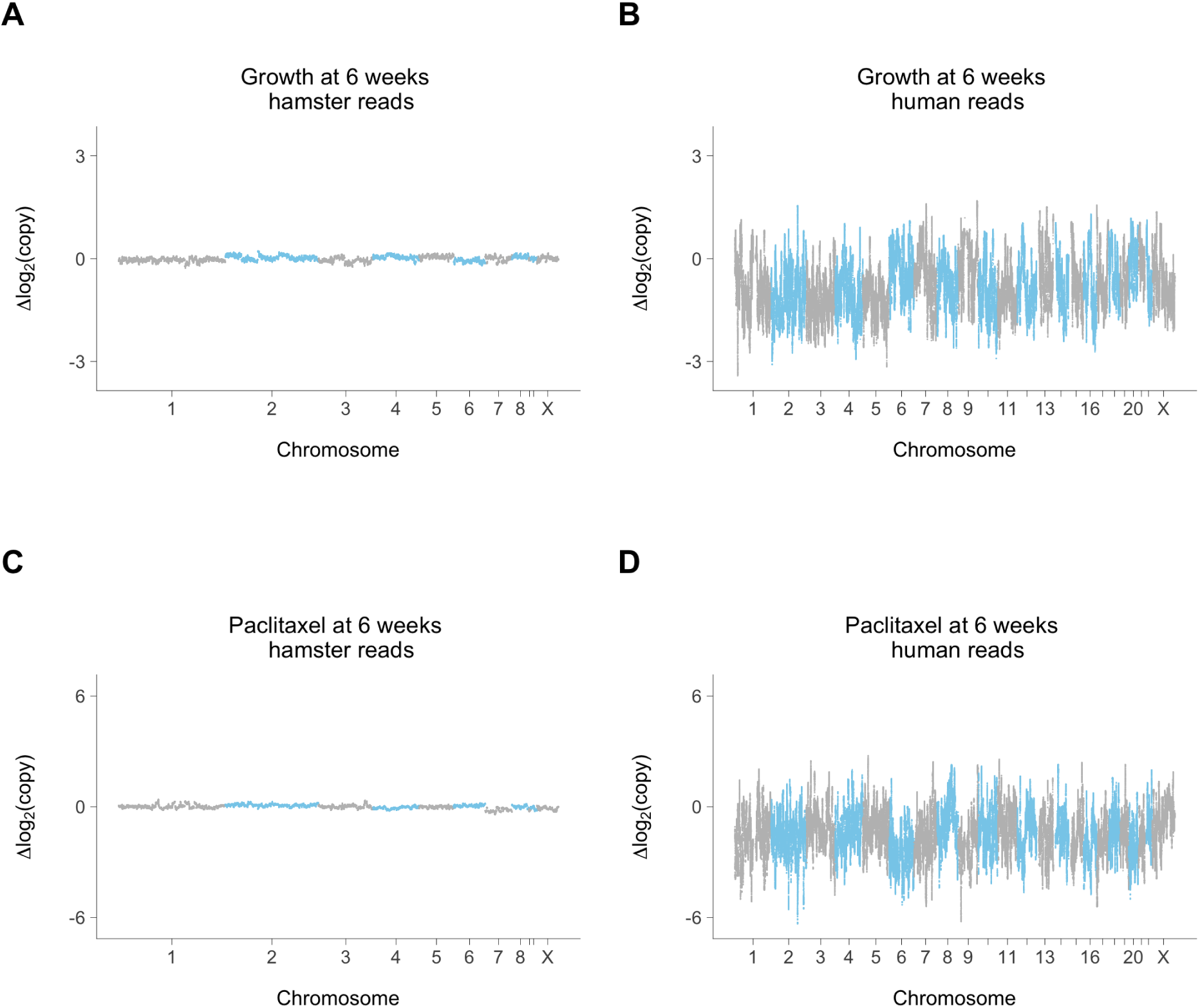
Copy number changes at week 6. (**A**) Hamster genome, growth at week 6 compared to week 0; 0 nM paclitaxel. (**B**) Human genome, growth at week 6 compared to week 0; 0 nM paclitaxel. (C) Hamster genome, 75 nM paclitaxel compared to 0 nM paclitaxel; week 6. (**D**) Human genome, 75 nM paclitaxel compared to 0 nM paclitaxel; week 6. Hamster and human copy number changes on same log_2_ scale, representing change in relative copy number normalized to hamster genome averaged across the six RH pools.

**Figure S11.**
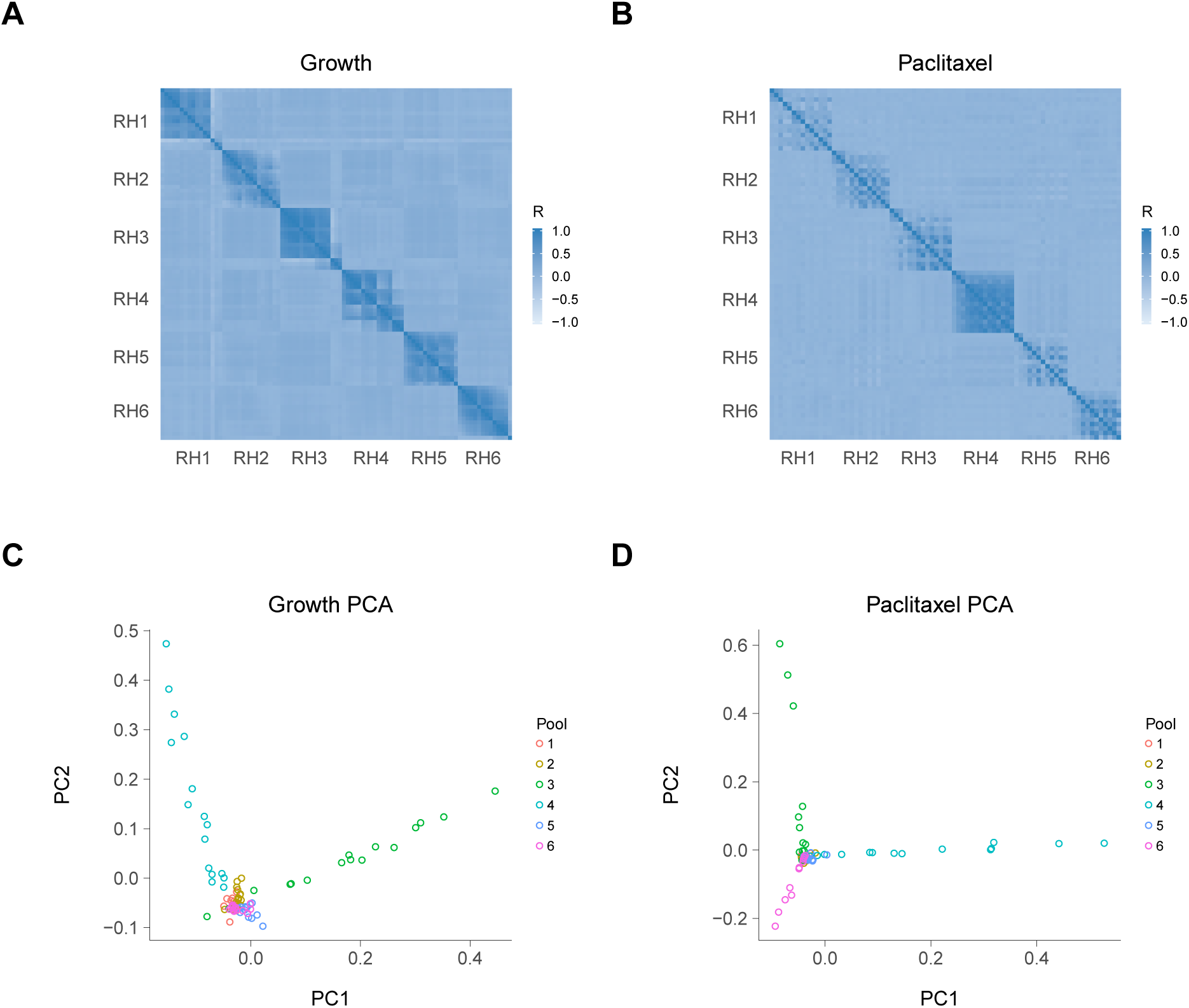
Human copy number changes reveal RH pool batch effects. (**A**) Correlations in human copy changes as a result of increasing growth times. (**B**) Correlations in human copy changes as a result of increasing paclitaxel concentrations. (**C**) Principal components analysis (PCA) of human copy number changes due to growth. PC1 explains 16% of the variance and PC2 explains 15%. (**D**) PCA of human copy number changes due to paclitaxel. PC1 explains 23% of the variance and PC2 explains 15%. Mean normalized human reads used, log_2_ scale. Non-overlapping 1 Mb windows.

**Figure S12.**
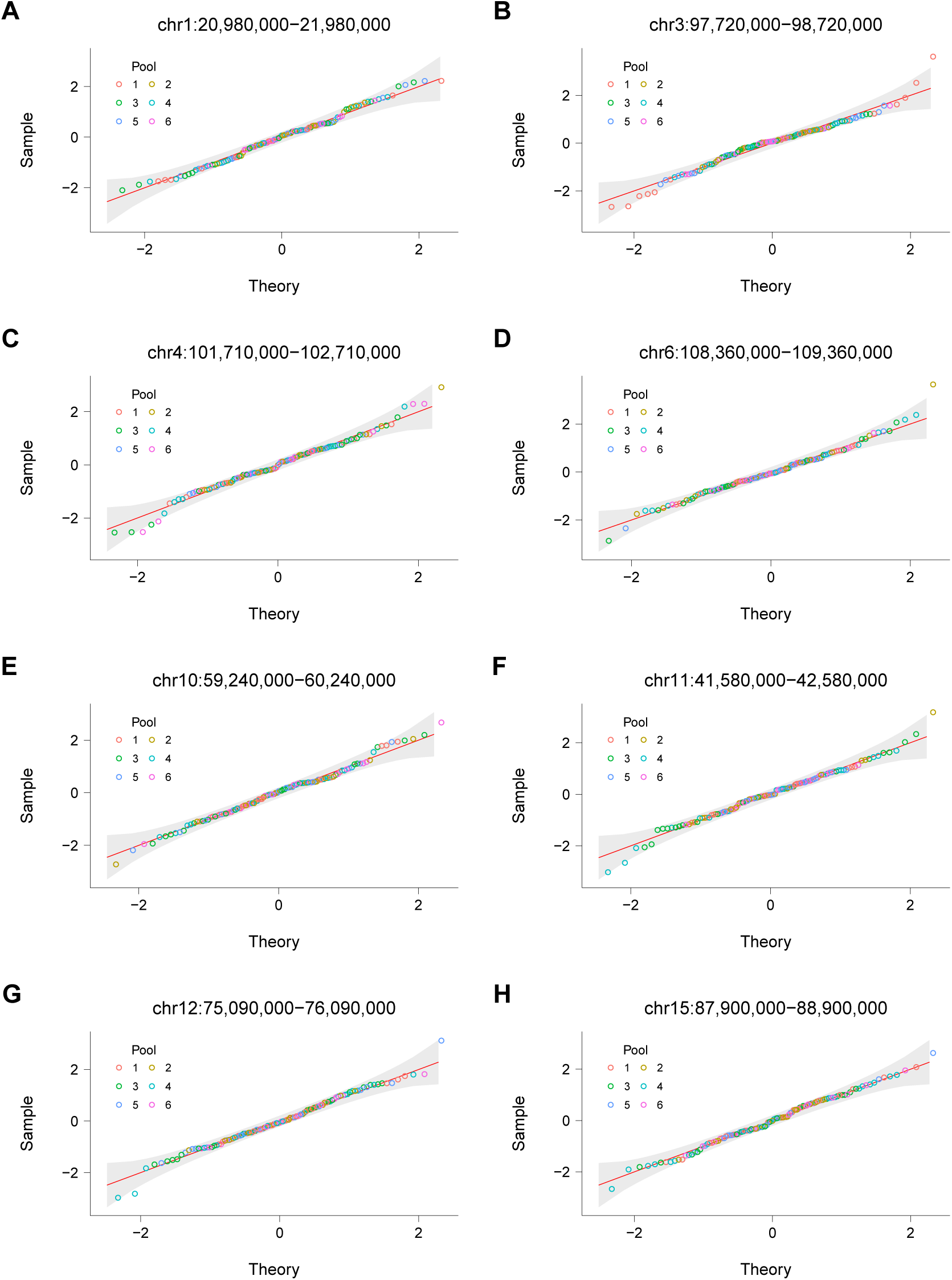
Quantile-quantile plots of residuals. (**A**) Chromosome 1, 20 980 000 bp to 21 980 000 bp. (**B**) Chromosome 3, 97 720 000 bp to 98 720 000 bp. (**C**) Chromosome 4, 101 710 000 bp to 102 710 000 bp. (**D**) Chromosome 6, 108 360 000 bp to 109 360 000 bp. (**E**) Chromosome 10, 59 240 000 bp to 60 240 000 bp. (**F**) Chromosome 11, 41 580 000 bp to 42 580 000 bp. (**G**) Chromosome 12, 75 090 000 bp to 76 090 000 bp. (**H**) Chromosome 15, 87 900 000 bp to 88 900 000 bp. Red line, linear best fit. Grey band, 95% confidence interval.

**Figure S13.**
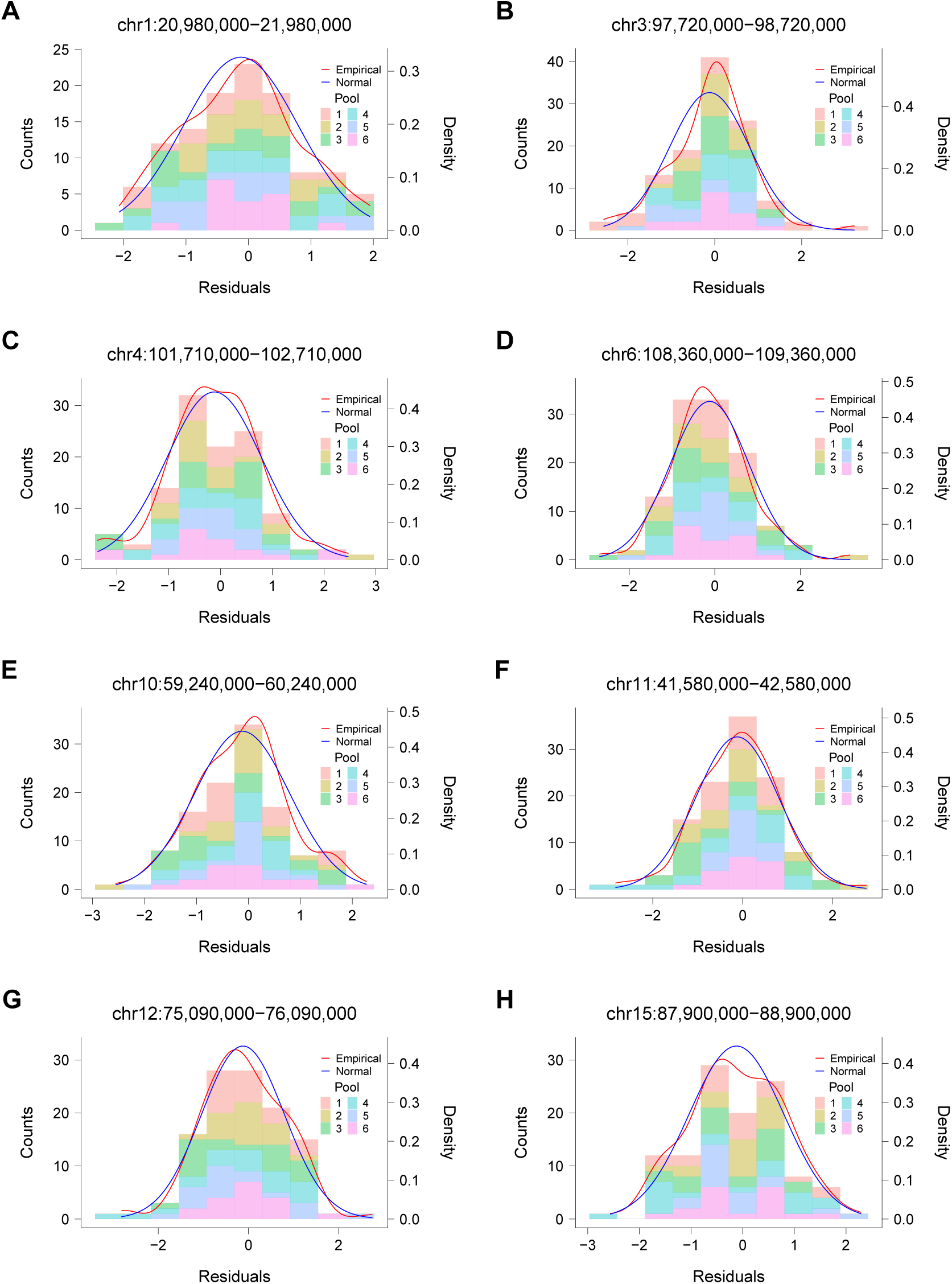
Sample counts plotted against residuals. (**A**) Chromosome 1, 20 980 000 bp to 21 980 000 bp. (**B**) Chromosome 3, 97 720 000 bp to 98 720 000 bp. (**C**) Chromosome 4, 101 710 000 bp to 102 710 000 bp. (**D**) Chromosome 6, 108 360 000 bp to 109 360 000 bp. (**E**) Chromosome 10, 59 240 000 bp to 60 240 000 bp. (**F**) Chromosome 11, 41 580 000 bp to 42 580 000 bp. (**G**) Chromosome 12, 75 090 000 bp to 76 090 000 bp. (**H**) Chromosome 15, 87 900 000 bp to 88 900 000 bp.

**Figure S14.**
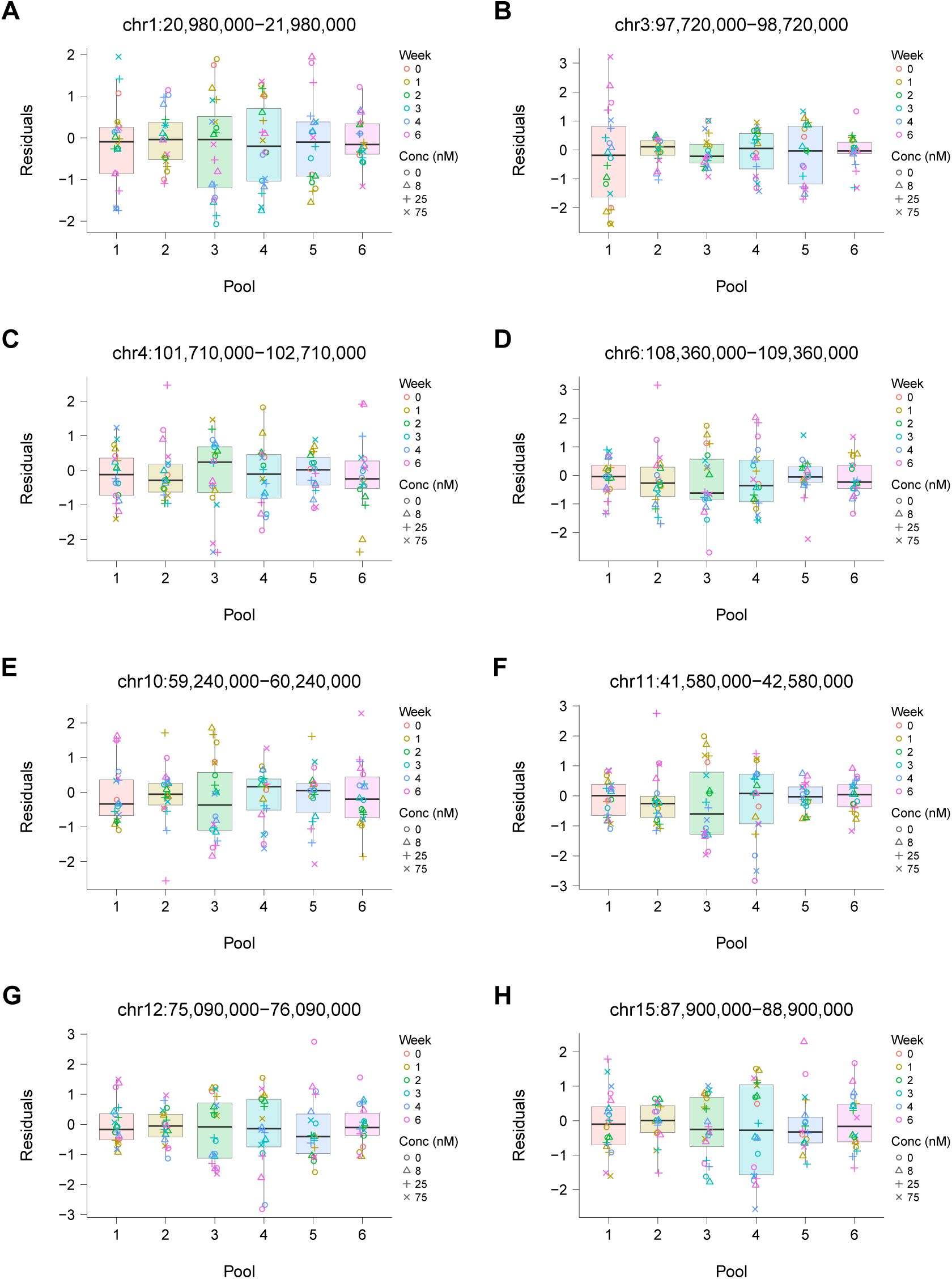
Residuals plotted against the RH pools. (**A**) Chromosome 1, 20 980 000 bp to 21 980 000 bp. (**B**) Chromosome 3, 97 720 000 bp to 98 720 000 bp. (**C**) Chromosome 4, 101 710 000 bp to 102 710 000 bp. (**D**) Chromosome 6, 108 360 000 bp to 109 360 000 bp. (**E**) Chromosome 10, 59 240 000 bp to 60 240 000 bp. (**F**) Chromosome 11, 41 580 000 bp to 42 580 000 bp. (**G**) Chromosome 12, 75 090 000 bp to 76 090 000 bp. (**H**) Chromosome 15, 87 900 000 bp to 88 900 000 bp.

**Figure S15.**
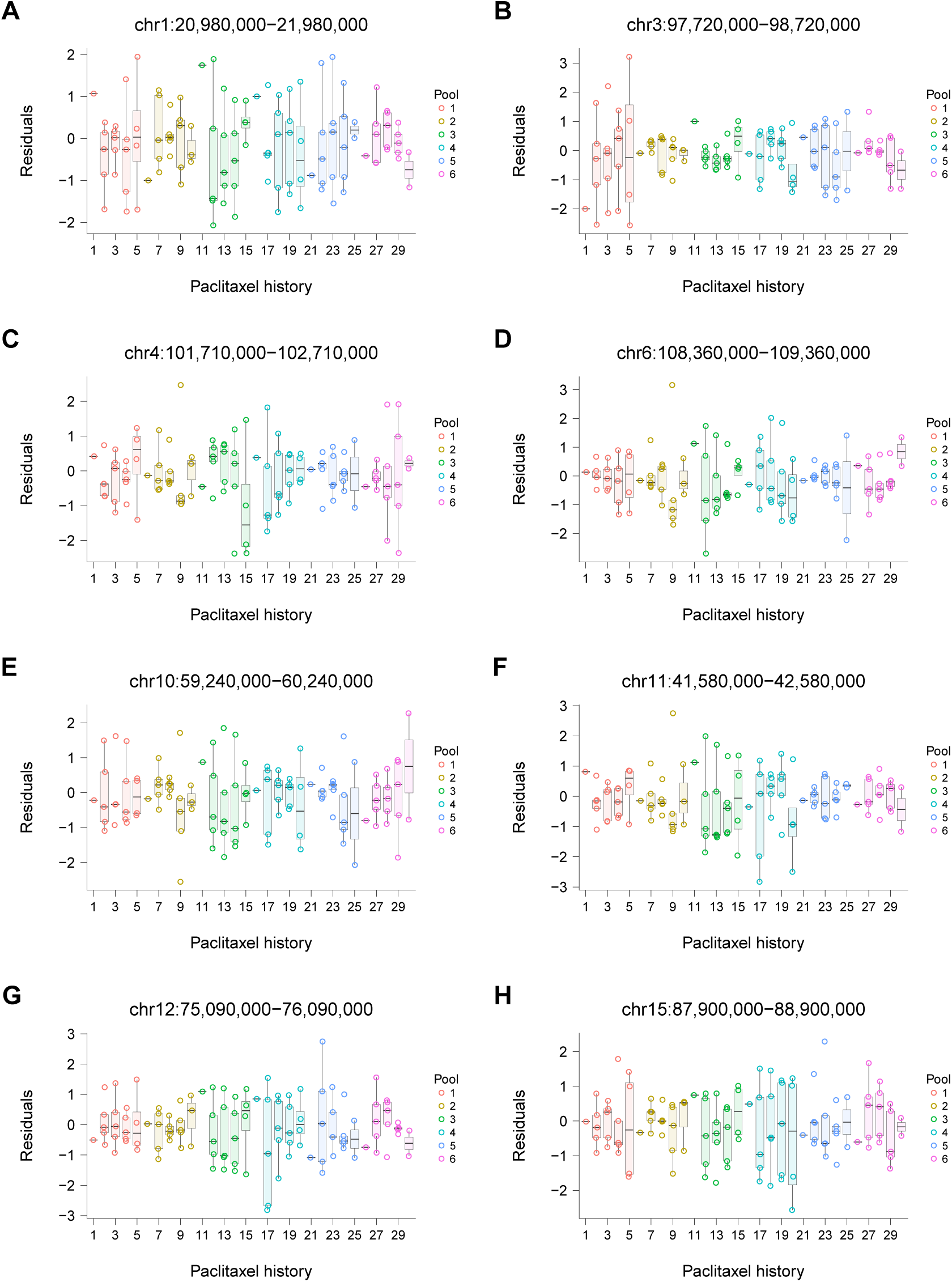
Residuals plotted against paclitaxel history. (**A**) Chromosome 1, 20 980 000 bp to 21 980 000 bp. (**B**) Chromosome 3, 97 720 000 bp to 98 720 000 bp. (**C**) Chromosome 4, 101 710 000 bp to 102 710 000 bp. (**D**) Chromosome 6, 108 360 000 bp to 109 360 000 bp. (**E**) Chromosome 10, 59 240 000 bp to 60 240 000 bp. (**F**) Chromosome 11, 41 580 000 bp to 42 580 000 bp. (**G**) Chromosome 12, 75 090 000 bp to 76 090 000 bp. (**H**) Chromosome 15, 87 900 000 bp to 88 900 000 bp. Paclitaxel history is a nested random effect term that accounts for possible differences in drug exposures.

**Figure S16.**
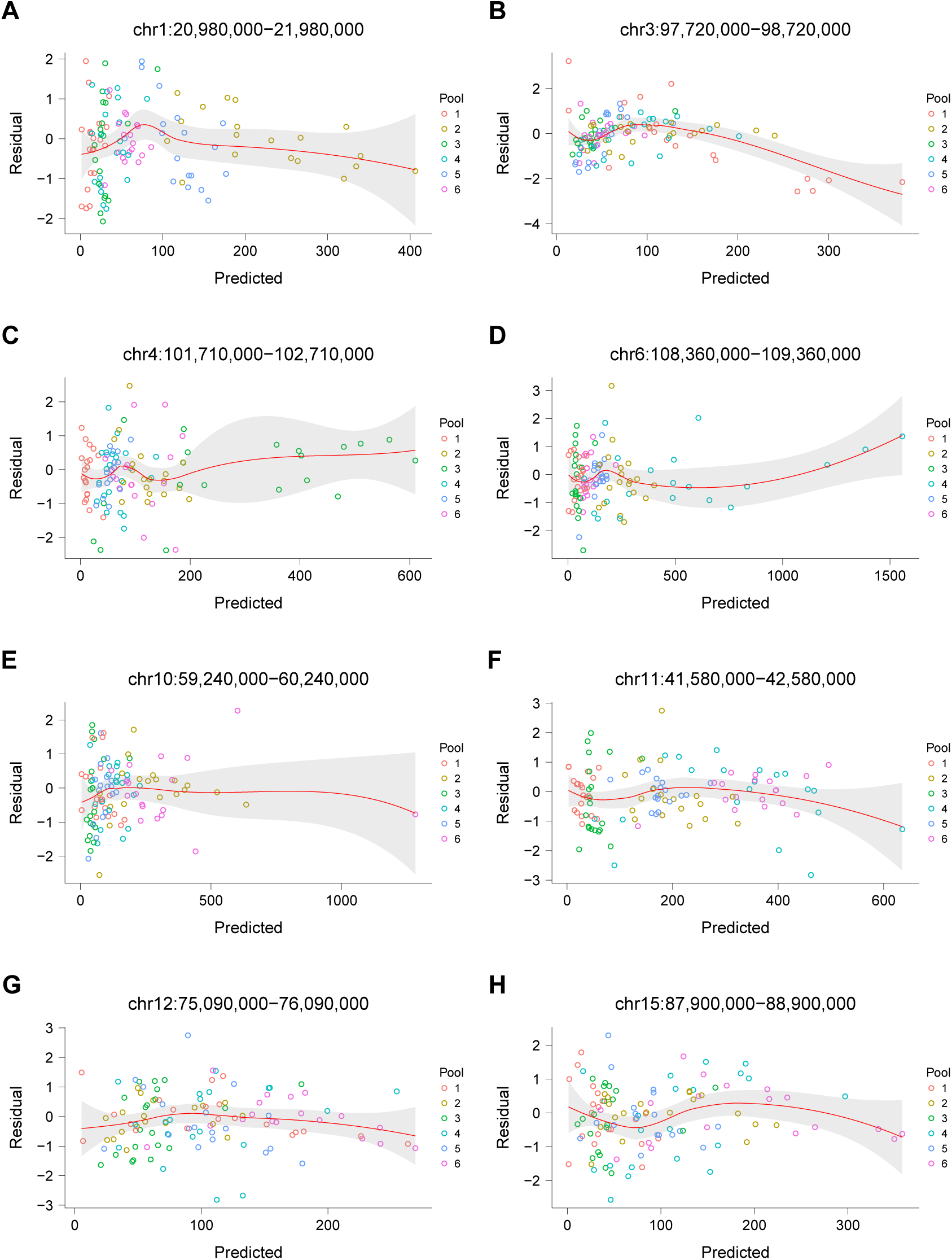
Residuals plotted against predicted values. (**A**) Chromosome 1, 20 980 000 bp to 21 980 000 bp. (**B**) Chromosome 3, 97 720 000 bp to 98 720 000 bp. (**C**) Chromosome 4, 101 710 000 bp to 102 710 000 bp. (**D**) Chromosome 6, 108 360 000 bp to 109 360 000 bp. (**E**) Chromosome 10, 59 240 000 bp to 60 240 000 bp. (**F**) Chromosome 11, 41 580 000 bp to 42 580 000 bp. (**G**) Chromosome 12, 75 090 000 bp to 76 090 000 bp. (**H**) Chromosome 15, 87 900 000 bp to 88 900 000 bp. Red line, loess best fit. Grey band, 95% confidence interval.

**Figure S17.**
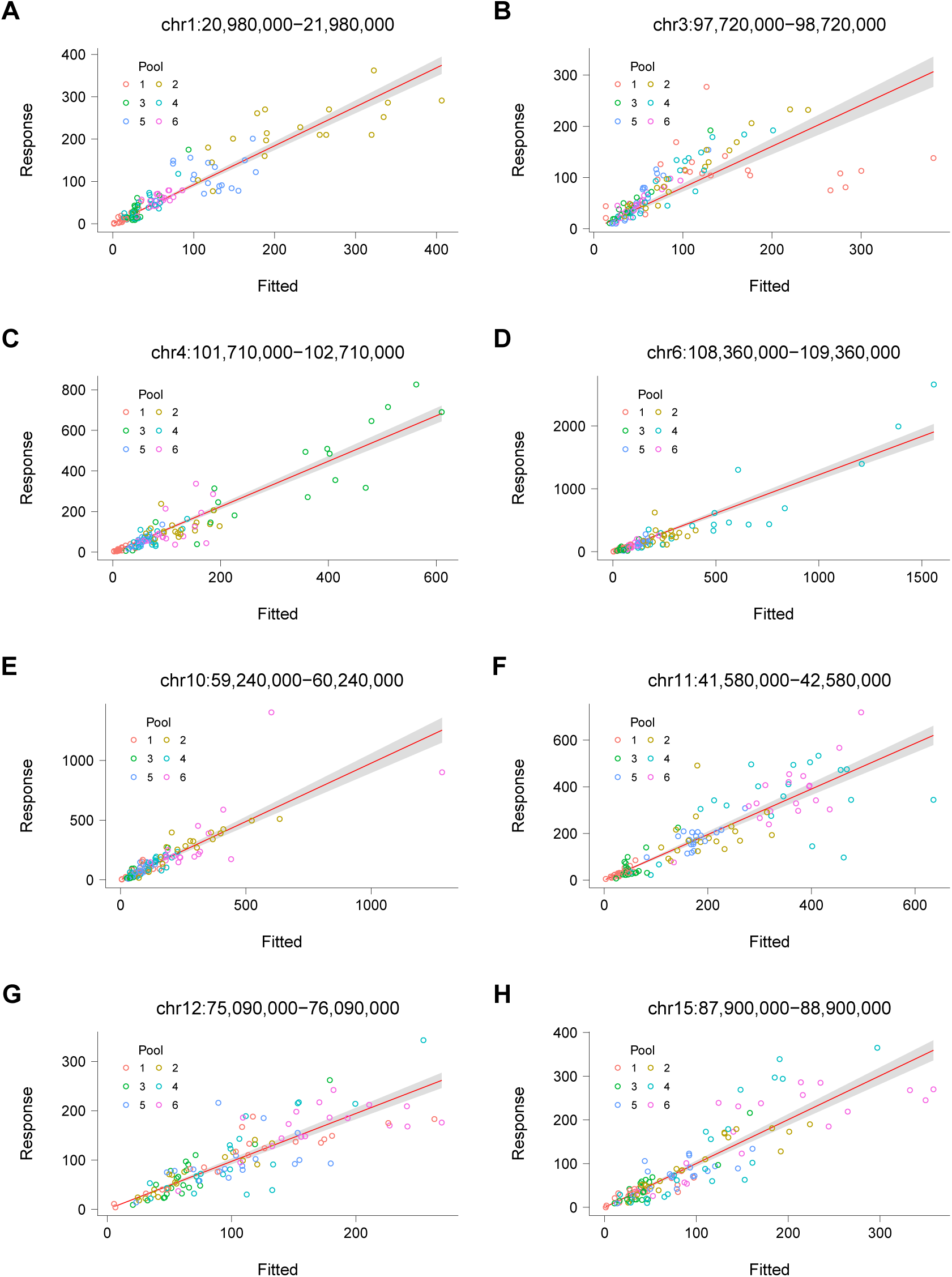
Response plotted against fitted values. (**A**) Chromosome 1, 20 980 000 bp to 21 980 000 bp. (**B**) Chromosome 3, 97 720 000 bp to 98 720 000 bp. (**C**) Chromosome 4, 101 710 000 bp to 102 710 000 bp. (**D**) Chromosome 6, 108 360 000 bp to 109 360 000 bp. (**E**) Chromosome 10, 59 240 000 bp to 60 240 000 bp. (**F**) Chromosome 11, 41 580 000 bp to 42 580 000 bp. (**G**) Chromosome 12, 75 090 000 bp to 76 090 000 bp. (**H**) Chromosome 15, 87 900 000 bp to 88 900 000 bp. Response shown as human specific sequence reads in each 1 Mb window. Red line, linear best fit. Grey band, 95% confidence interval.

**Figure S18.**
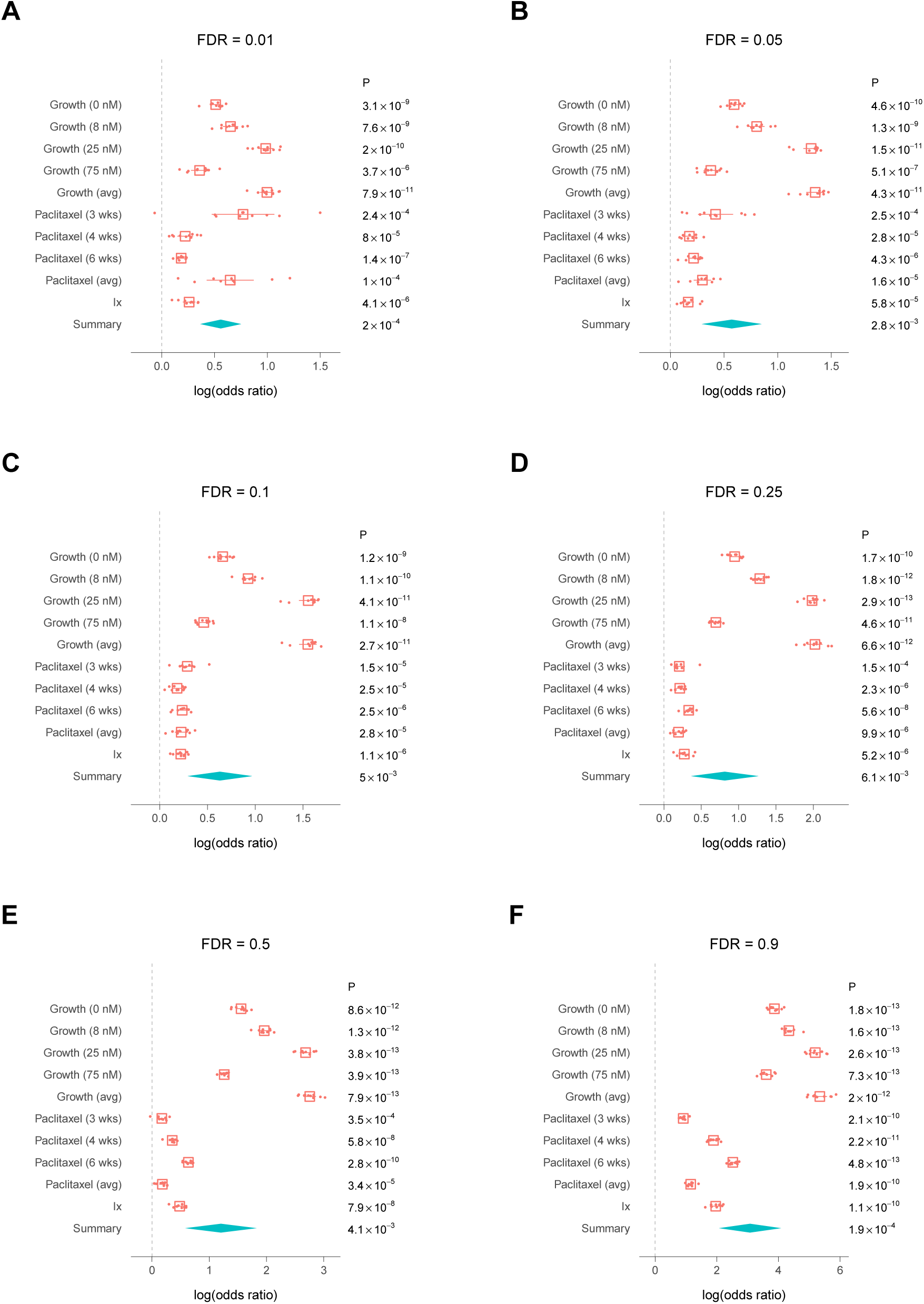
Cross validation. (**A**) False discovery rate (FDR) = 0.01. (**B**) FDR = 0.05. (**C**) FDR = 0.1. (**D**) FDR = 0.25. (**E**) FDR = 0.5. (**F**) FDR = 0.9. Means, center of boxes. Whiskers and summary diamonds, 95% confidence intervals.

**Figure S19.**
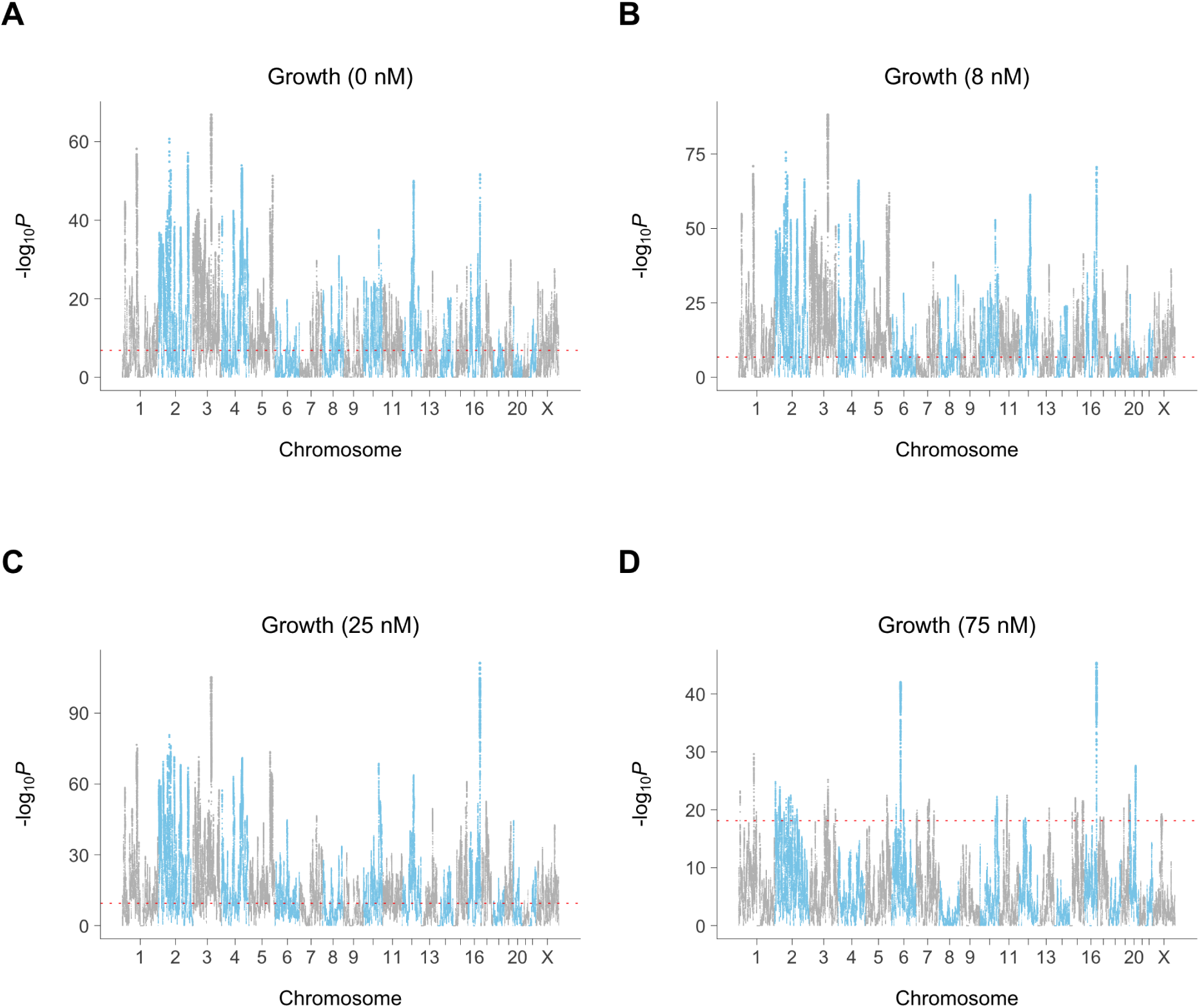
Growth loci. (**A**) 0 nM paclitaxel. (**B**) 8 nM paclitaxel. (**C**) 25 nM paclitaxel. (**D**) 75 nM paclitaxel. Red dotted line, permutation significance threshold.

**Figure S20.**
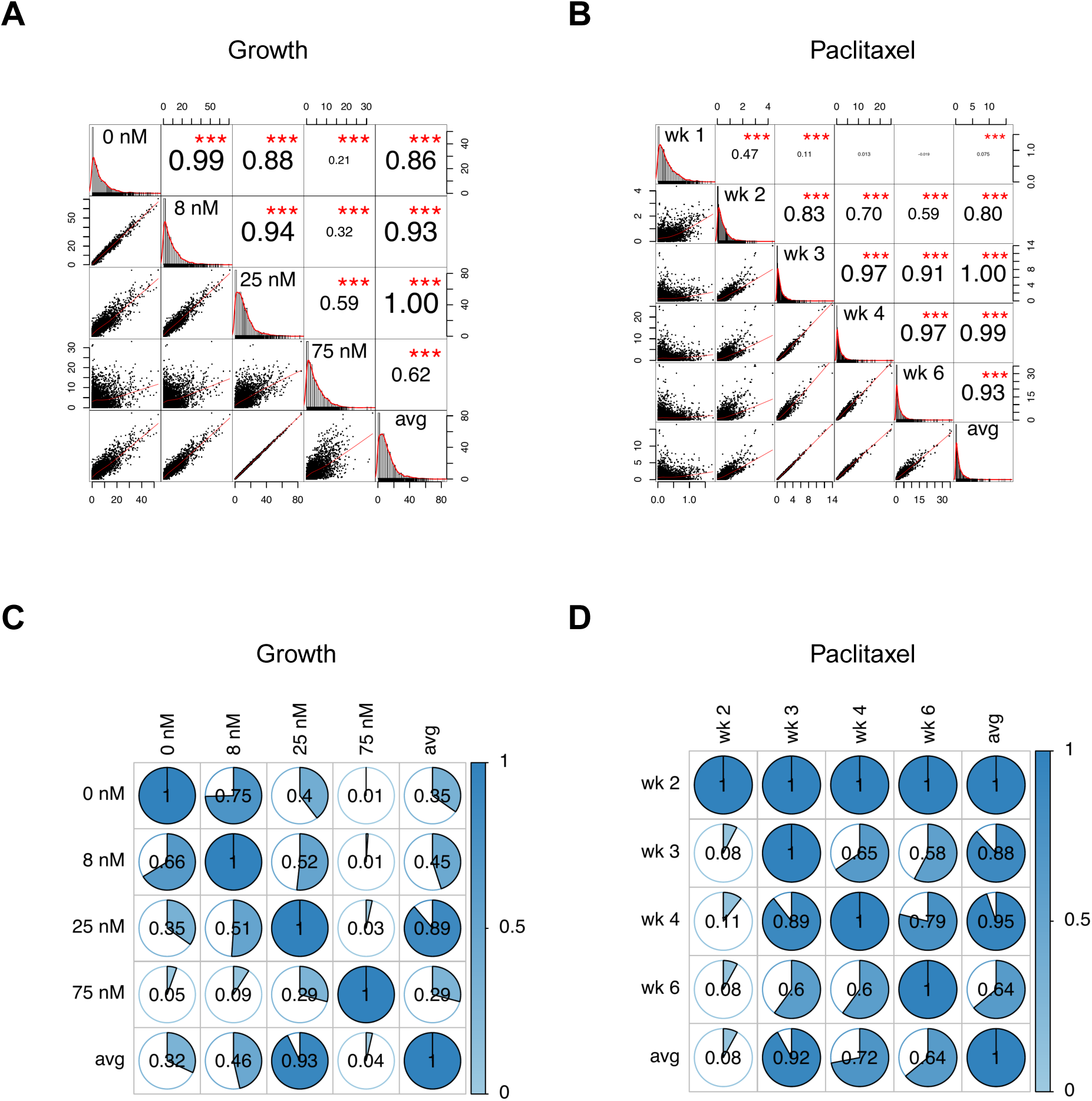
Overlap of non-unique loci. (**A**) Correlations of −log_10_ *P* values for growth at different paclitaxel concentrations. Numbers of varying size are *R* values for correlations between various genome scans. \**P <* 0.05, \*\**P <* 0.01, \*\*\**P <* 0.001. Non-overlapping 1 Mb windows. (**B**) Correlations of −log_10_ *P* values for paclitaxel at different timepoints. (**C**) Overlap of significant non-unique growth loci. Numbers inside circles represent overlap ratios using diagonal element of relevant row. (**D**) Overlap of significant non-unique paclitaxel loci. “avg”, average conditional effect of growth (A, C) or paclitaxel (B, D).

**Figure S21.**
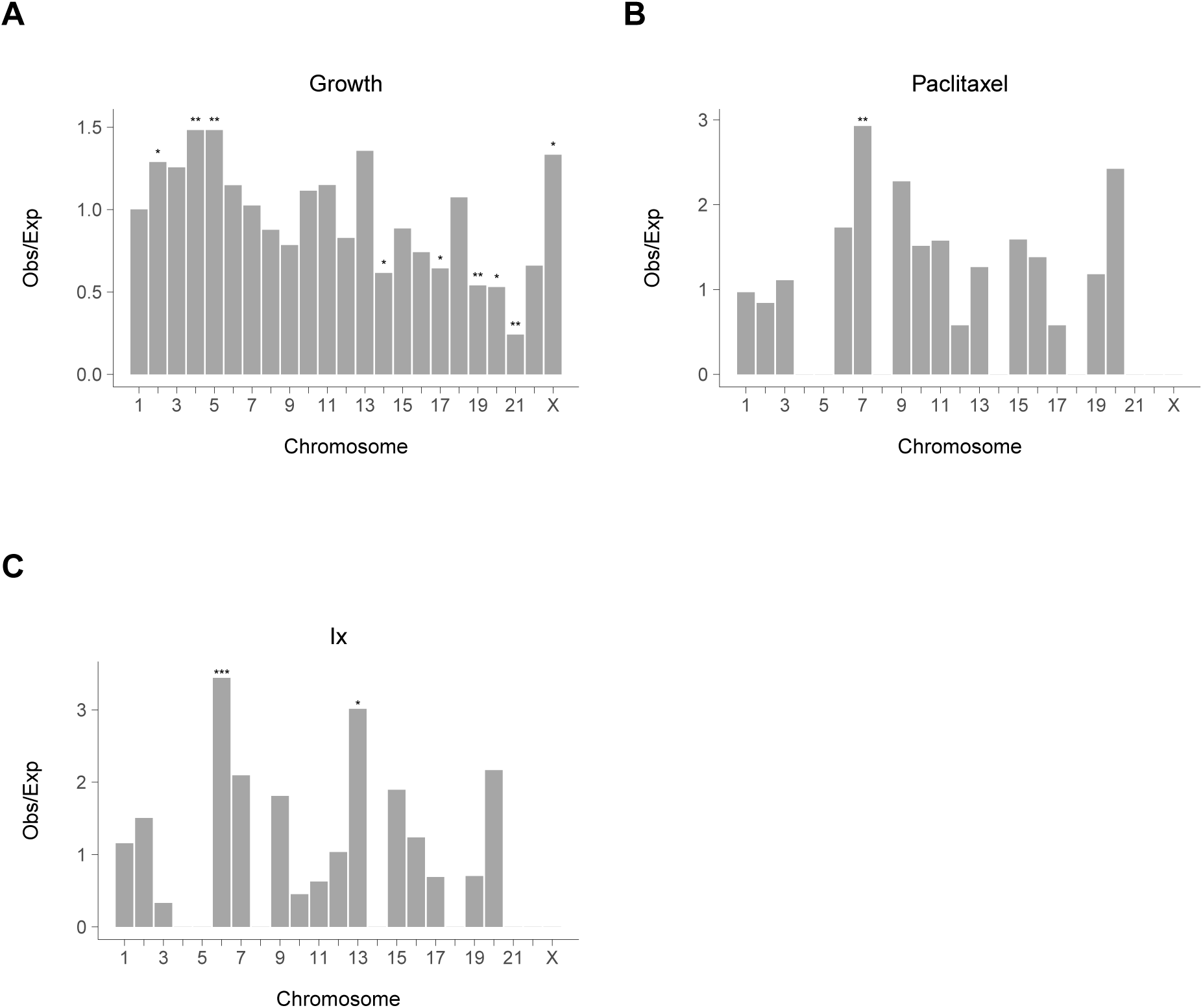
Number of loci on each chromosome. (**A**) Growth. (**B**) Paclitaxel. (**C**) Interaction. Observed vs expected ratio for unique RH loci shown, centromeres excluded. \**P <* 0.05, \*\**P <* 0.01, \*\*\**P <* 0.001, *χ*^2^ tests.

**Figure S22.**
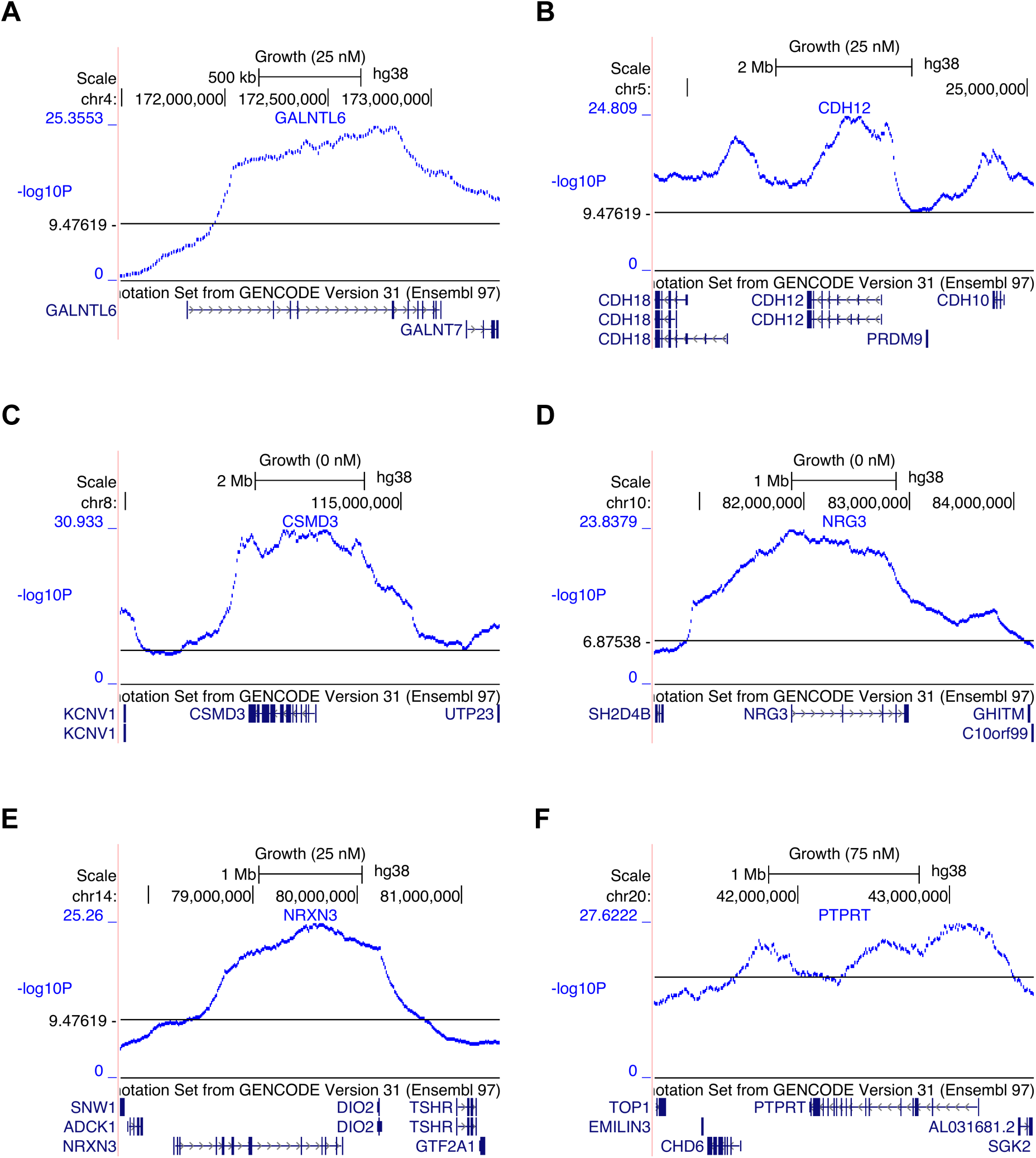
Long growth genes. (**A**) GALNTL6. (**B**) CDH12. (**C**) CSMD3. (**D**) NRG3. (**E**) NRXN3. (**F**) PTPRT. The −log_10_ *P* values follow the profiles of the long genes. Horizontal black lines, permutation significance thresholds.

**Figure S23.**
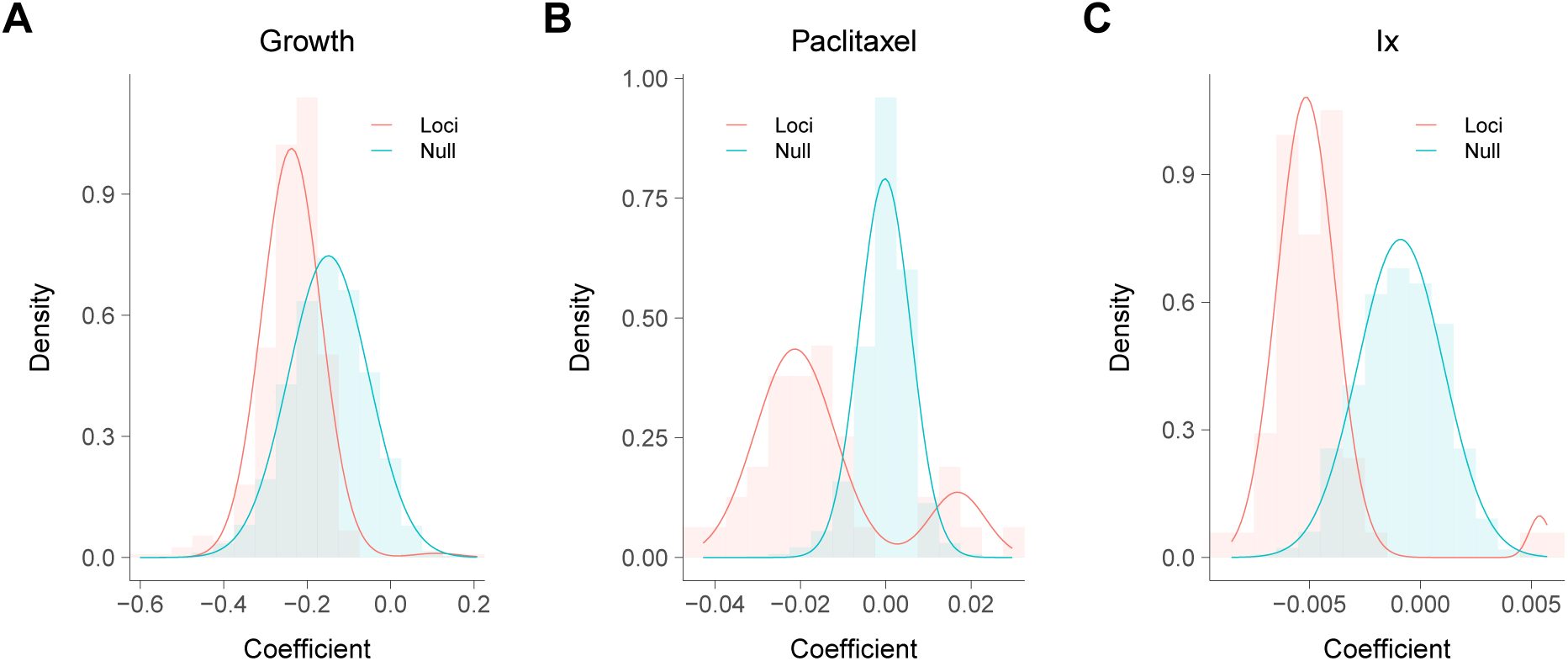
Loci coefficients. (**A**) Growth. (**B**) Paclitaxel. (**C**) Interaction. Empirical and best fit normal distributions for unique loci shown. Null based on non-overlapping 1 Mb windows.

**Figure S24.**
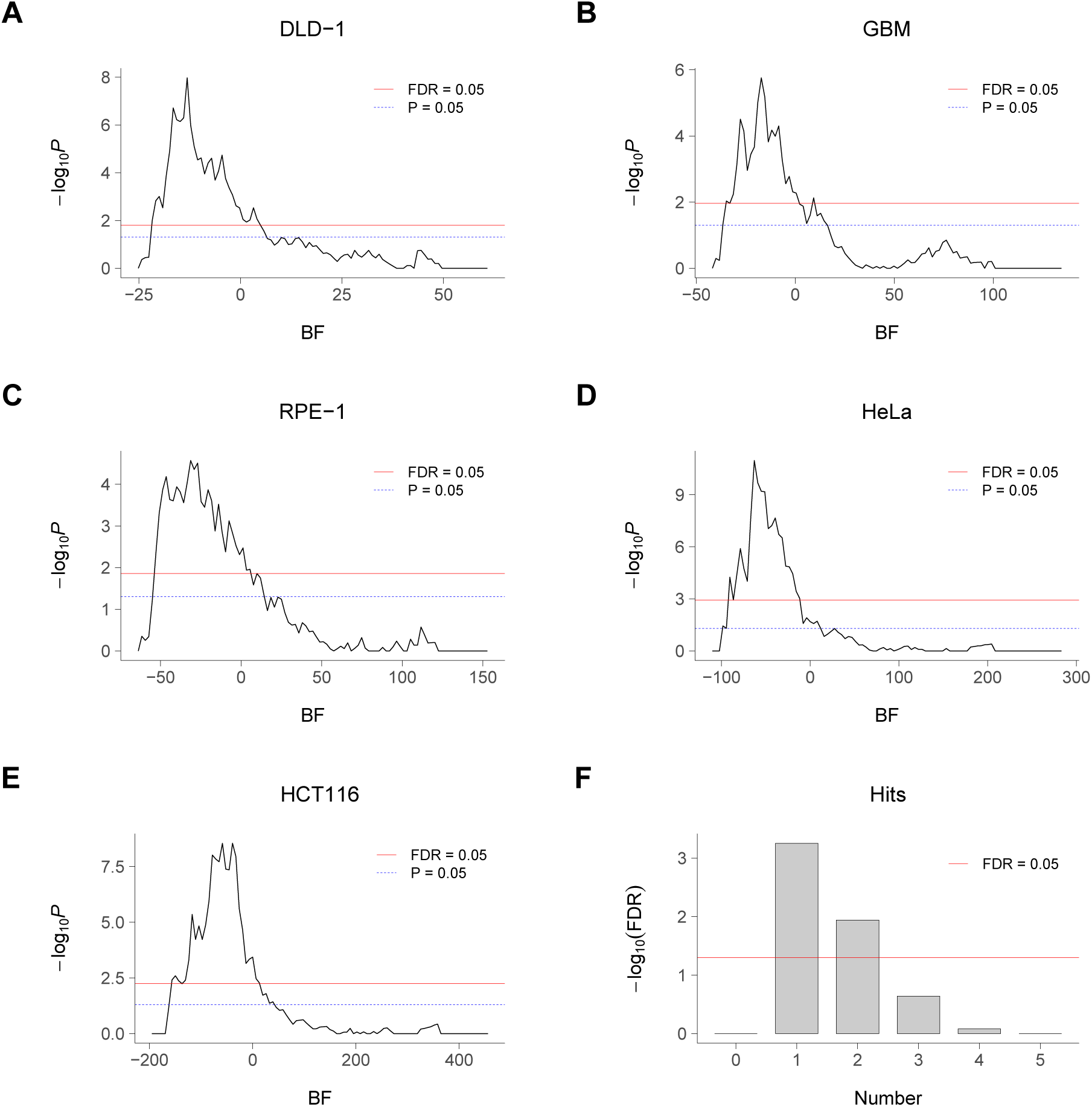
Significant non-overlap between RH and loss-of-function growth genes. (**A**) Comparison of RH and CRISPR growth genes in DLD-1 cells. Fisher’s exact test used to derive −log_10_ *P* values, thresholded on CRISPR growth effect measured by Bayes Factor (BF). (**B**) Patient derived glioblastoma (GBM) cells. (**C**) RPE-1 cells. (**D**) HeLa cells. (**E**) HCT116 cells. (**F**) Comparison thresholded on number of cell lines with CRISPR hits. Loss-of-function data from (Hart et al. 2015).

**Figure S25.**
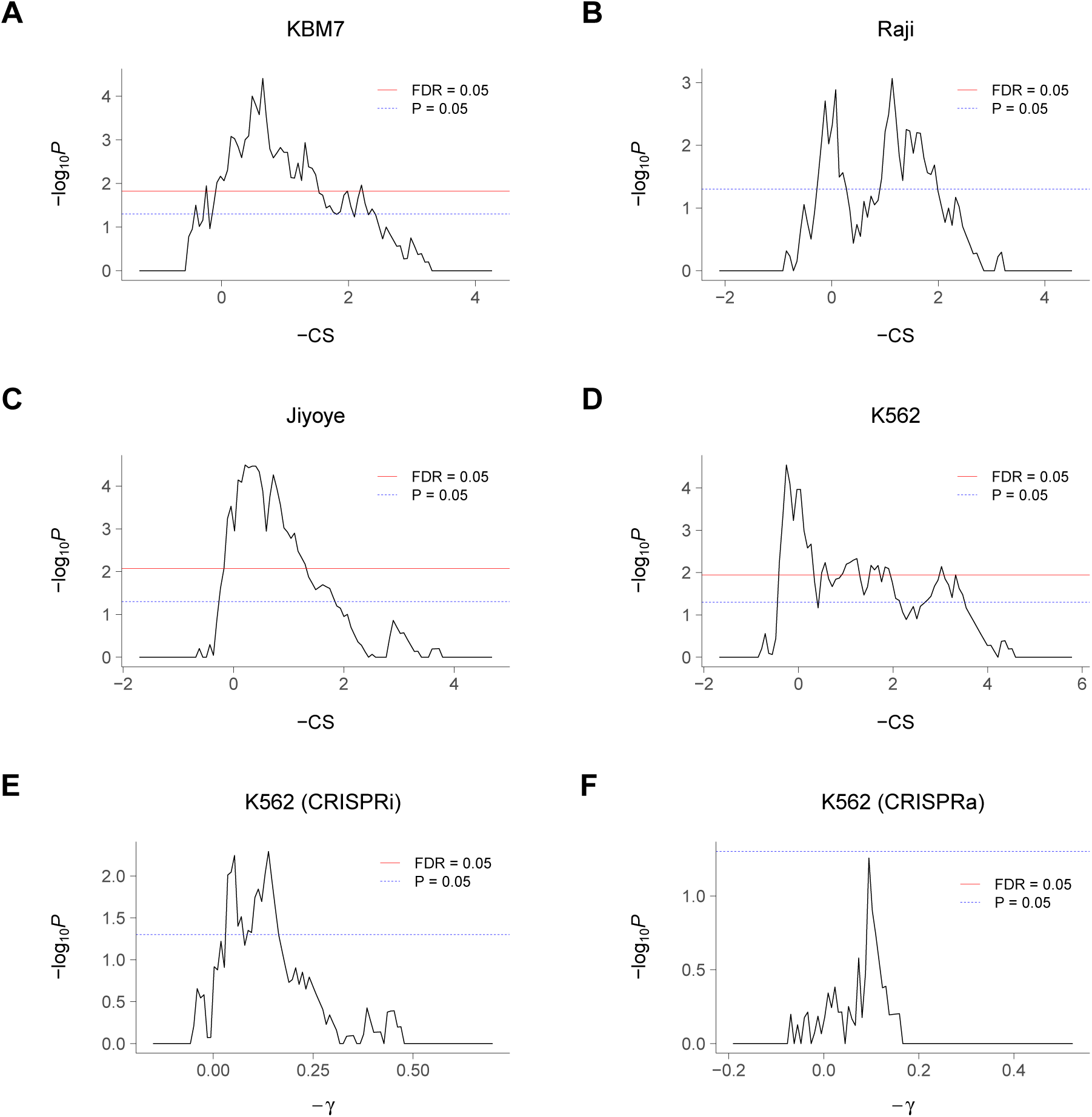
Further non-overlap between RH and CRISPR growth genes. (**A**) Comparison of RH and CRISPR loss-of-function growth genes in KBM7 cells. Fisher’s exact test used to derive −log_10_ *P* values, thresholded on CRISPR growth effect measured by CRISPR score (CS). (**B**) Raji cells. (**C**) Jiyoye cells. (**D**) K562 cells. Loss-of-function CRISPR data from (Wang, Birsoy, et al. 2015). (**E**) Comparison of RH and CRISPRi growth genes in K562 cells, thresholded on −*γ* score. (**F**) Comparison of RH and CRISPRa growth genes in K562 cells. CRISPRi and CRISPRa data from (Gilbert et al. 2014).

**Figure S26.**
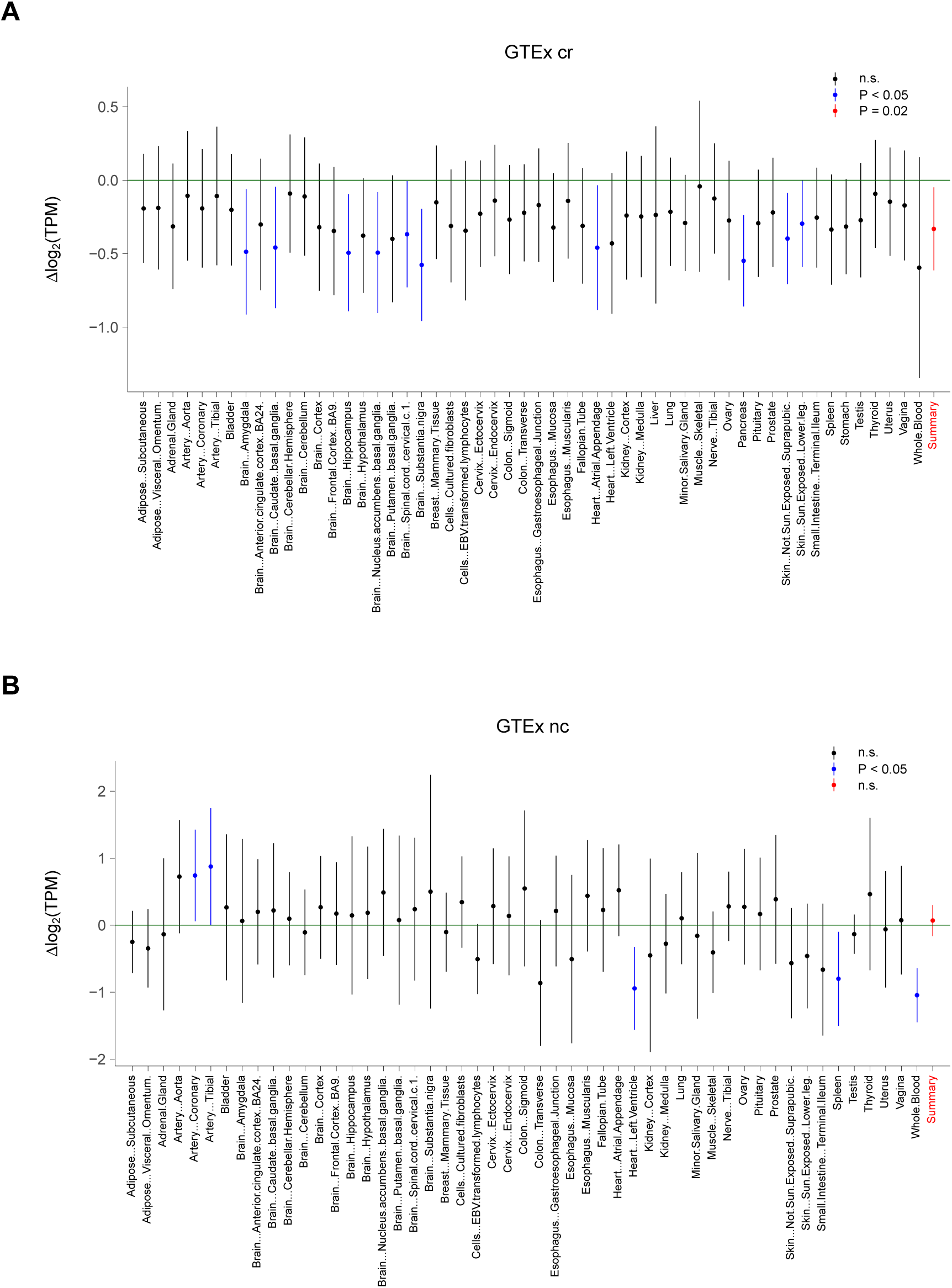
RH coding region growth genes have lower expression. (**A**) Expression differences between RH growth and non-growth coding region (cr) genes based on GTEx RNA-Seq data from various tissues. (**B**) Expression differences between RH growth and non-growth non-coding (nc) genes. Mean differences and 95% confidence intervals. Significant *P* values based on Welch Two Sample t-test shown in blue, summaries in red. Transcripts per million (TPM) thresholded at *≥* 5.

**Figure S27.**
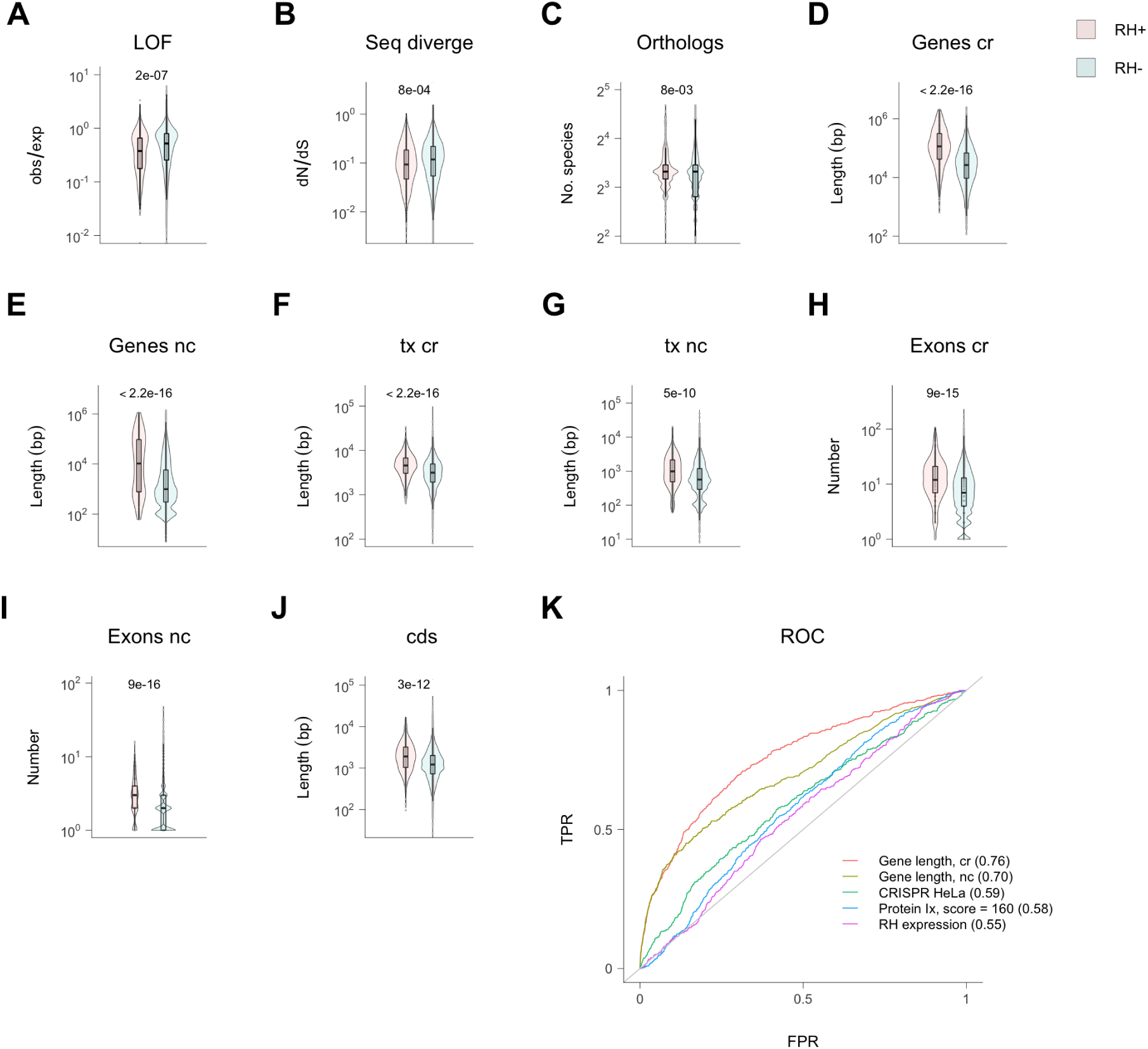
Further properties of RH growth genes. (**A**) Increased sensitivity to loss-of-function (LOF) variants (observed/expected) for RH growth genes. (**B**) Decreased human/mouse functional sequence divergence (dN/dS) of RH growth genes. (**C**) Increased number of species with orthologs of RH growth genes. (**D**) Increased gene lengths for coding region (cr) and (**E**) non-coding (nc) RH growth genes. (**F**) Increased transcript lengths for cr and (**G**) nc RH growth genes. (**H**) Increased exon numbers for cr and (**I**) nc RH growth genes. (**J**) Increased coding sequence (cds) lengths for cr RH growth genes. (**K**) ROC curves for RH growth genes. Area under the curve (AUC) given in brackets. *P* values from Welch Two Sample t-test, except ROC curves *P <* 4.9 × 10^−4^, Wilcoxon rank sum test.

**Figure S28.**
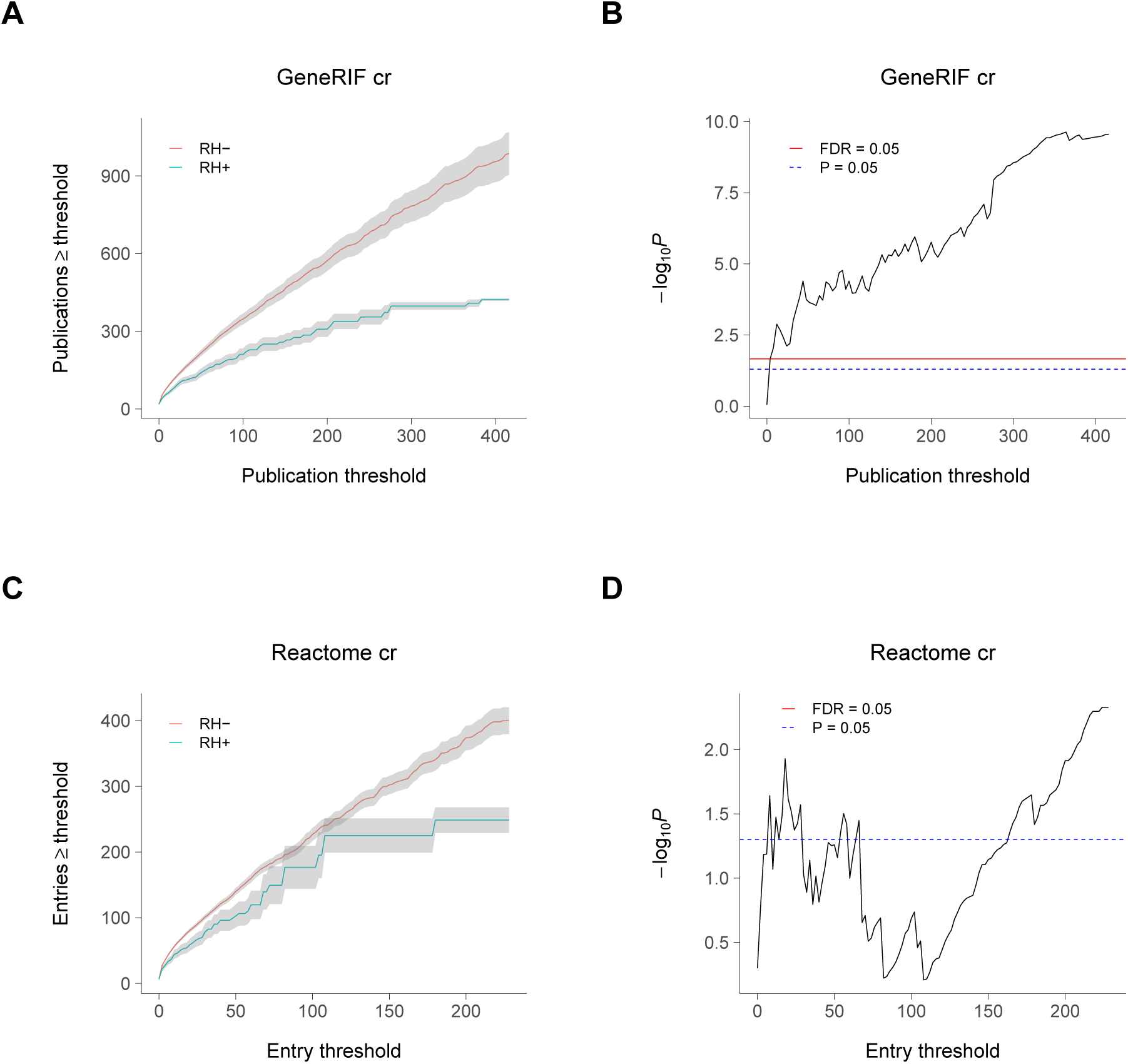
RH growth genes are relatively poorly studied. (**A**) RH coding region (cr) growth genes have fewer PubMed entries in the GeneRIF database than non-growth genes. Means ± s.e.m. (**B**) Significance of decreased publication numbers for RH cr growth genes. (**C**) RH cr growth genes have fewer entries in the Reactome database than non-growth genes. (**D**) Significance of decreased Reactome entries for RH cr growth genes.

**Figure S29.**
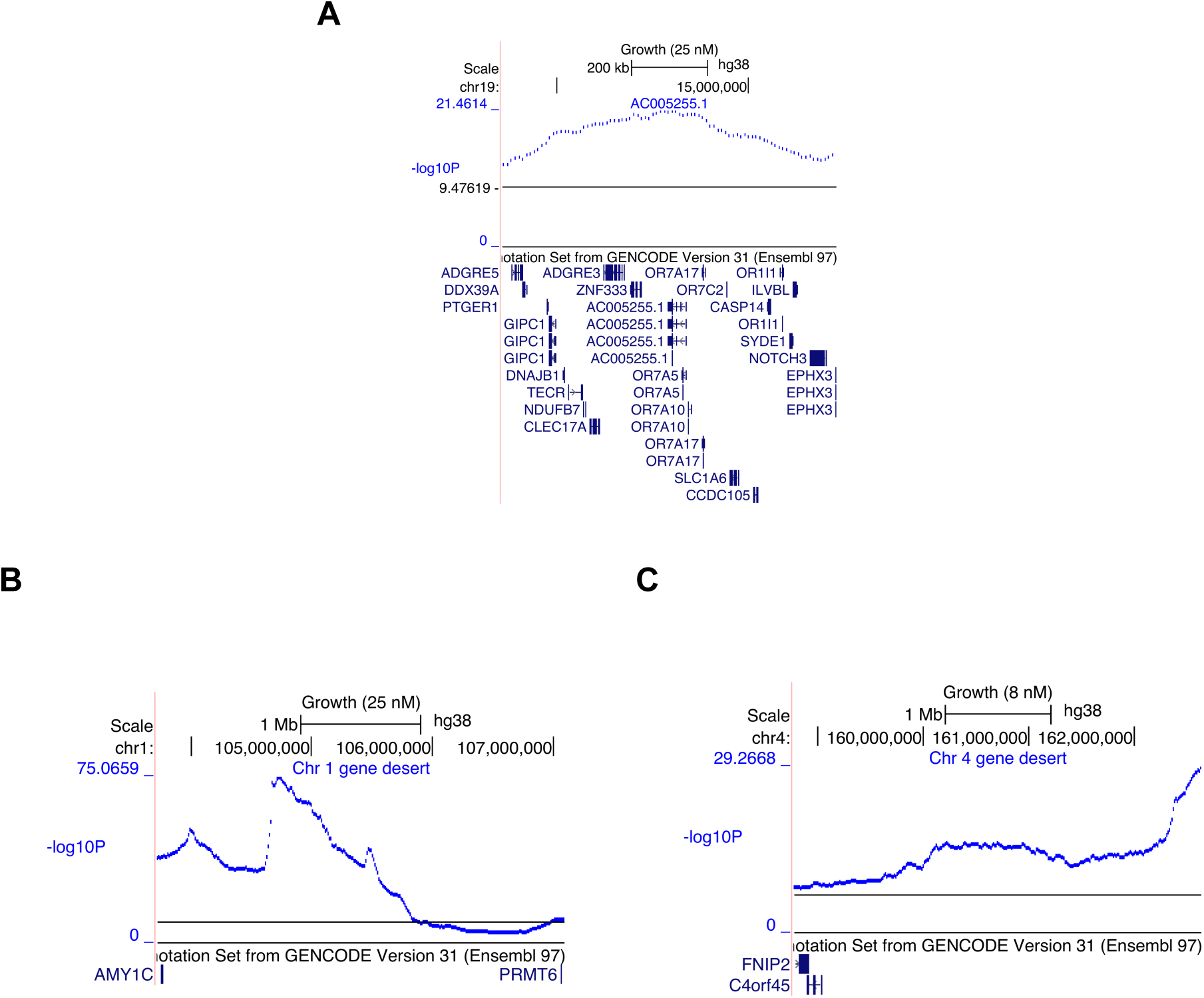
Olfactory and gene desert growth loci. (**A**) Growth locus mapping to olfactory receptor cluster on chromosome 19, with peak −log_10_ *P* value located in AC005255.1 (OR7C1) (25 nM paclitaxel). (**B**) Growth locus in gene desert on chromosome 1 at 104 730 000 bp (25 nM paclitaxel). (**C**) Growth locus in gene desert on chromosome 4 at 160 220 000 bp (8 nM paclitaxel). Horizontal black lines, permutation significance thresholds.

**Figure S30.**
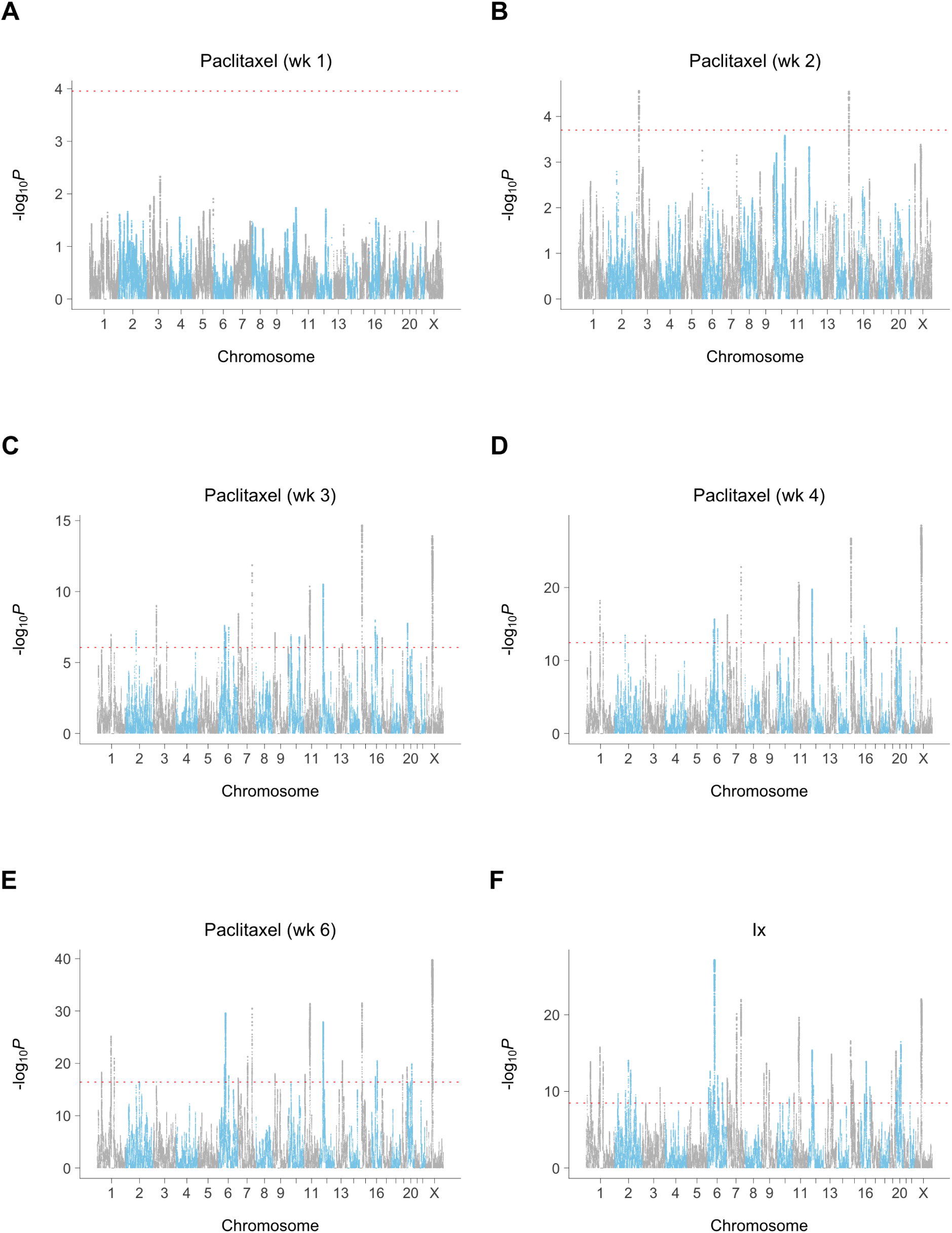
Paclitaxel and interaction loci. (**A**) Week 1. (**B**) Week 2. (**C**) Week 3. (**D**) Week 4. (**E**) Week 6. (**F**) Growth/paclitaxel interaction. Red dotted line, permutation significance threshold.

**Figure S31.**
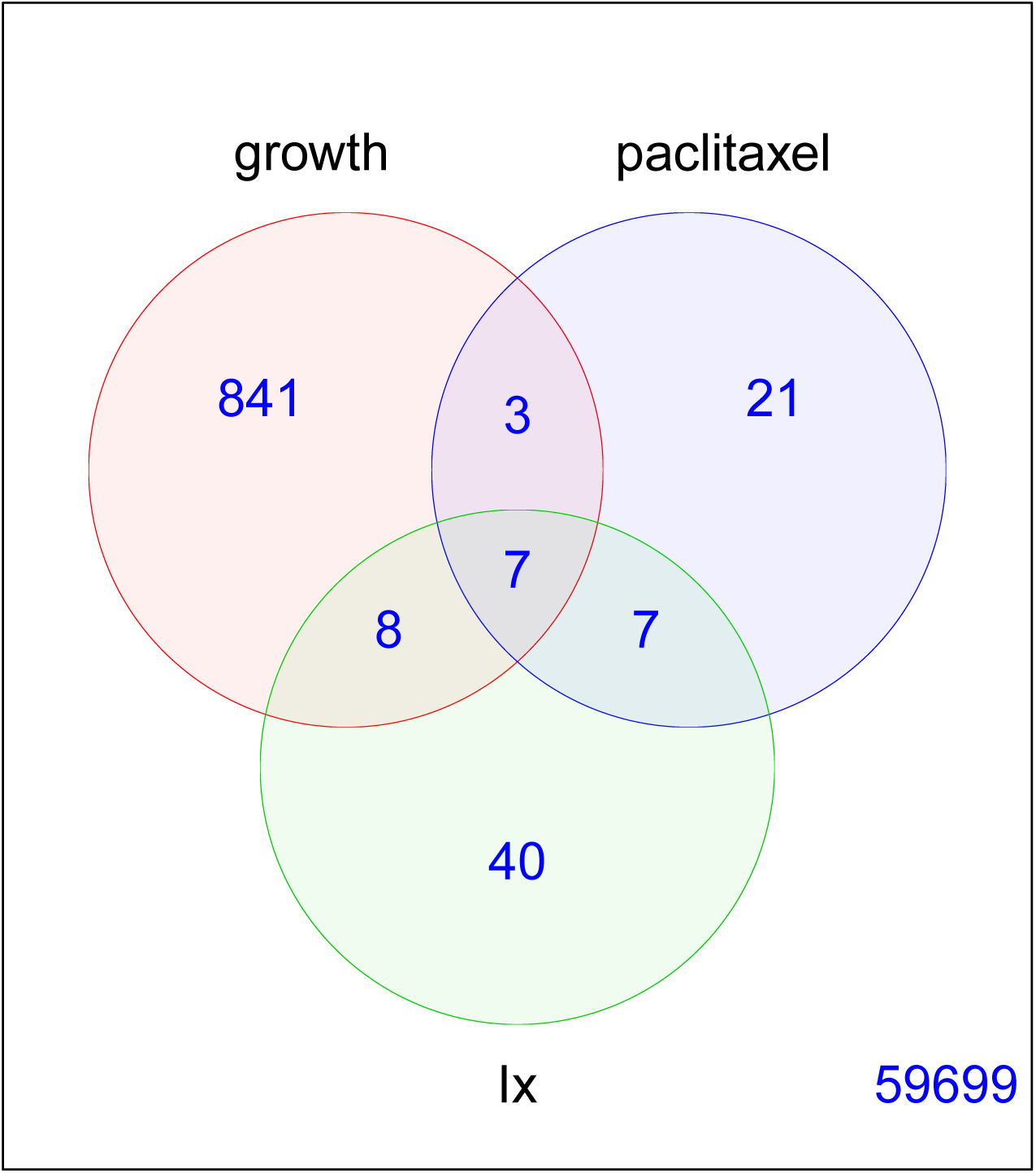
Overlap of unique growth, paclitaxel and interaction loci. Total gene numbers from GENCODE v31 (Frankish et al. 2019) plus centromeres on chromosomes 1–22 and X.

**Figure S32.**
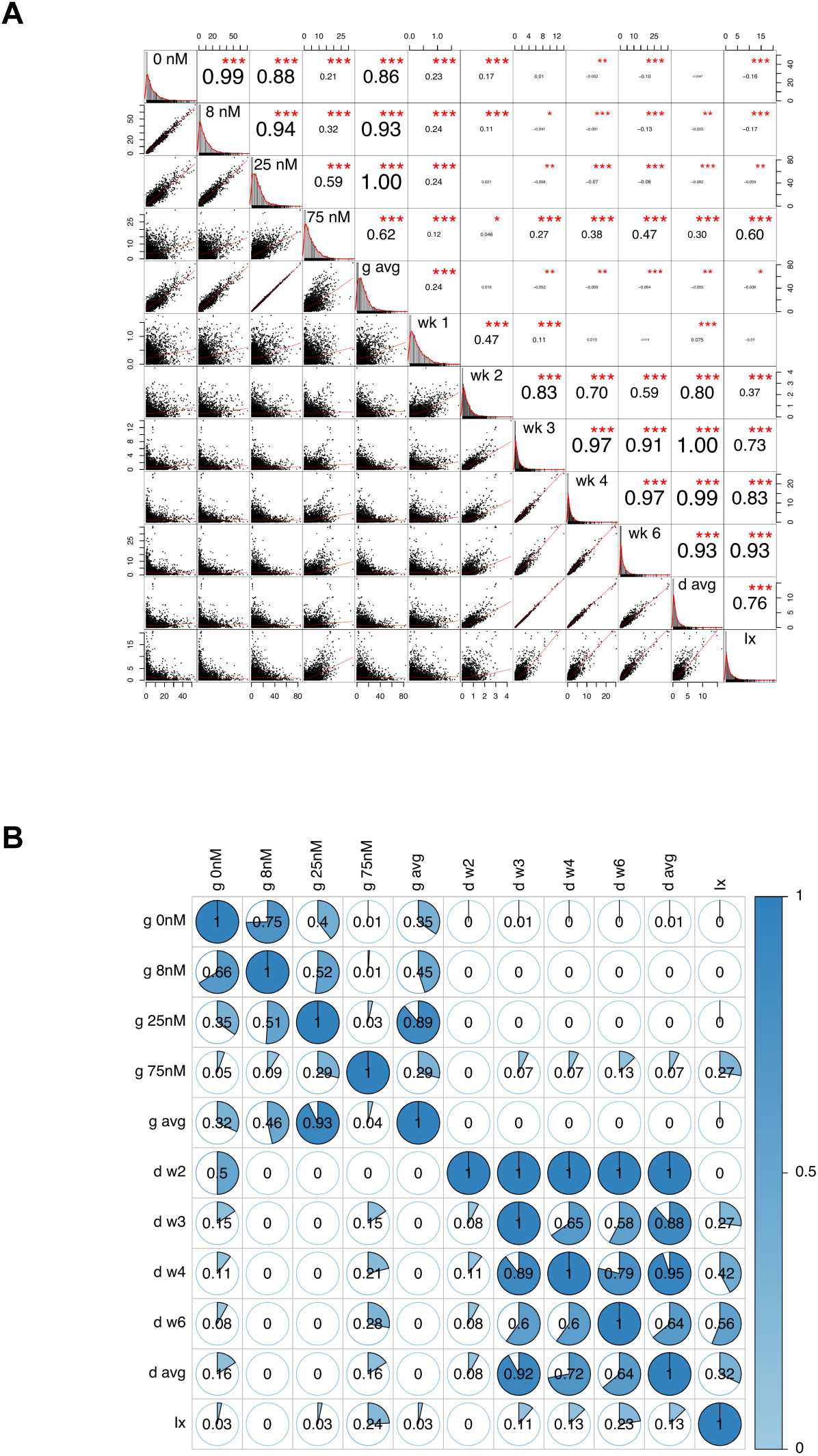
Overlap of non-unique growth, paclitaxel and interaction loci. (**A**) Correlations of −log_10_ *P* values for growth (0, 8, 25 and 75 nM; g avg), paclitaxel (w1, w2, w3, w4, w6 and d avg) and interaction (Ix). “g”, growth; “d”, drug; “avg”, average conditional effect. Numbers of varying size are *R* values for correlations between various genome scans. \**P <* 0.05, \*\**P <* 0.01, \*\*\**P <* 0.001. Non-overlapping 1 Mb windows. (**B**) Overlap of significant non-unique growth, paclitaxel and interaction loci. Numbers inside circles represent overlap ratios using diagonal element of relevant row.

**Figure S33.**
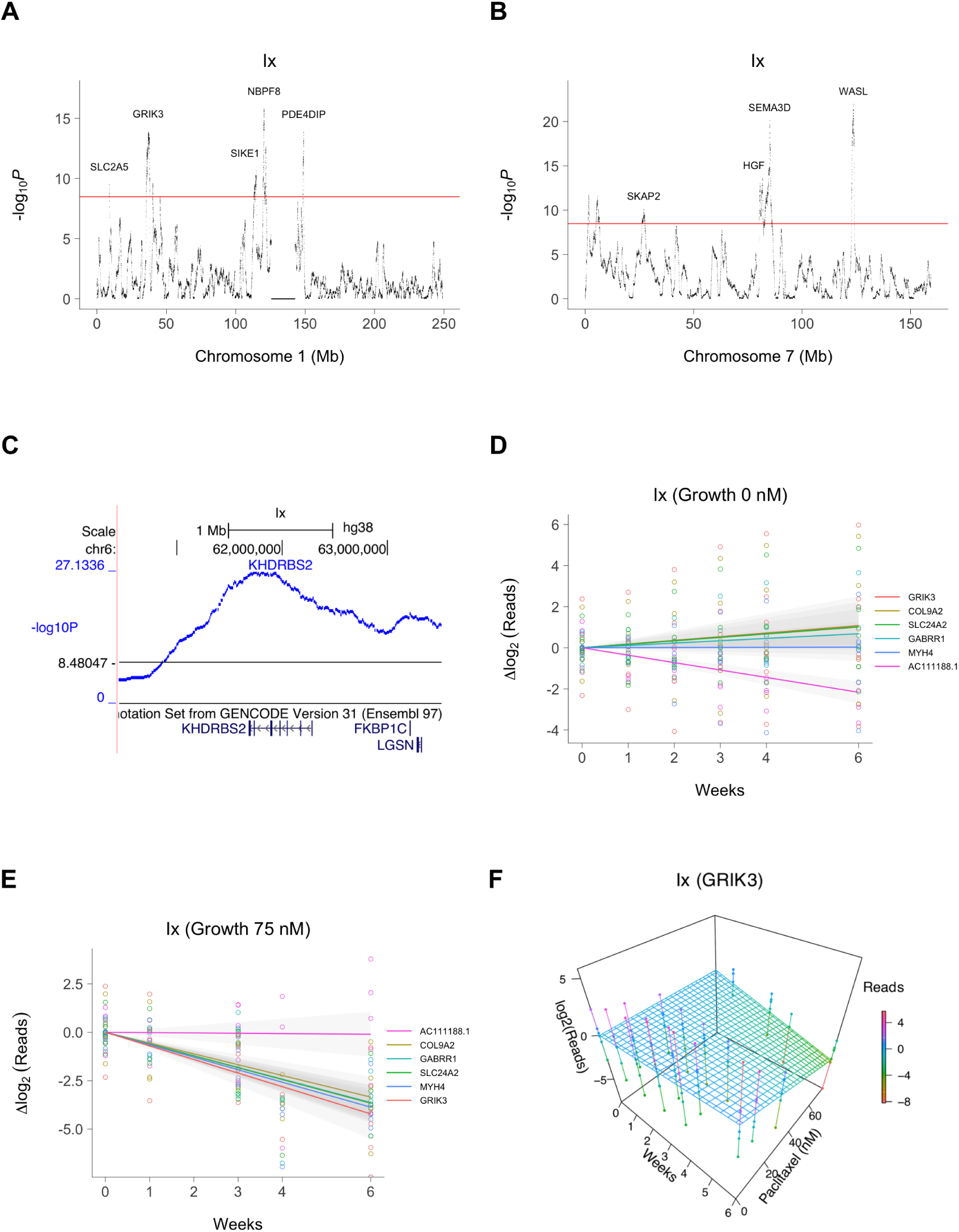
Close up views of interaction loci. (**A**) Chromosome 1. (**B**) Chromosome 7. (**C**) Locus for KHDRBS2 on chromosome 6. Horizontal red and black lines, permutation significance thresholds. (**D**) Normalized sequence reads on log_2_ scale showing growth at 0 xnM paclitaxel for six interaction loci. Colored lines, best fit. Grey bands, 95% confidence intervals. (**D**) Same loci showing growth at 75 nM paclitaxel. (**F**) GRIK3 interaction locus on chromosome 1, best fit.

**Figure S34.**
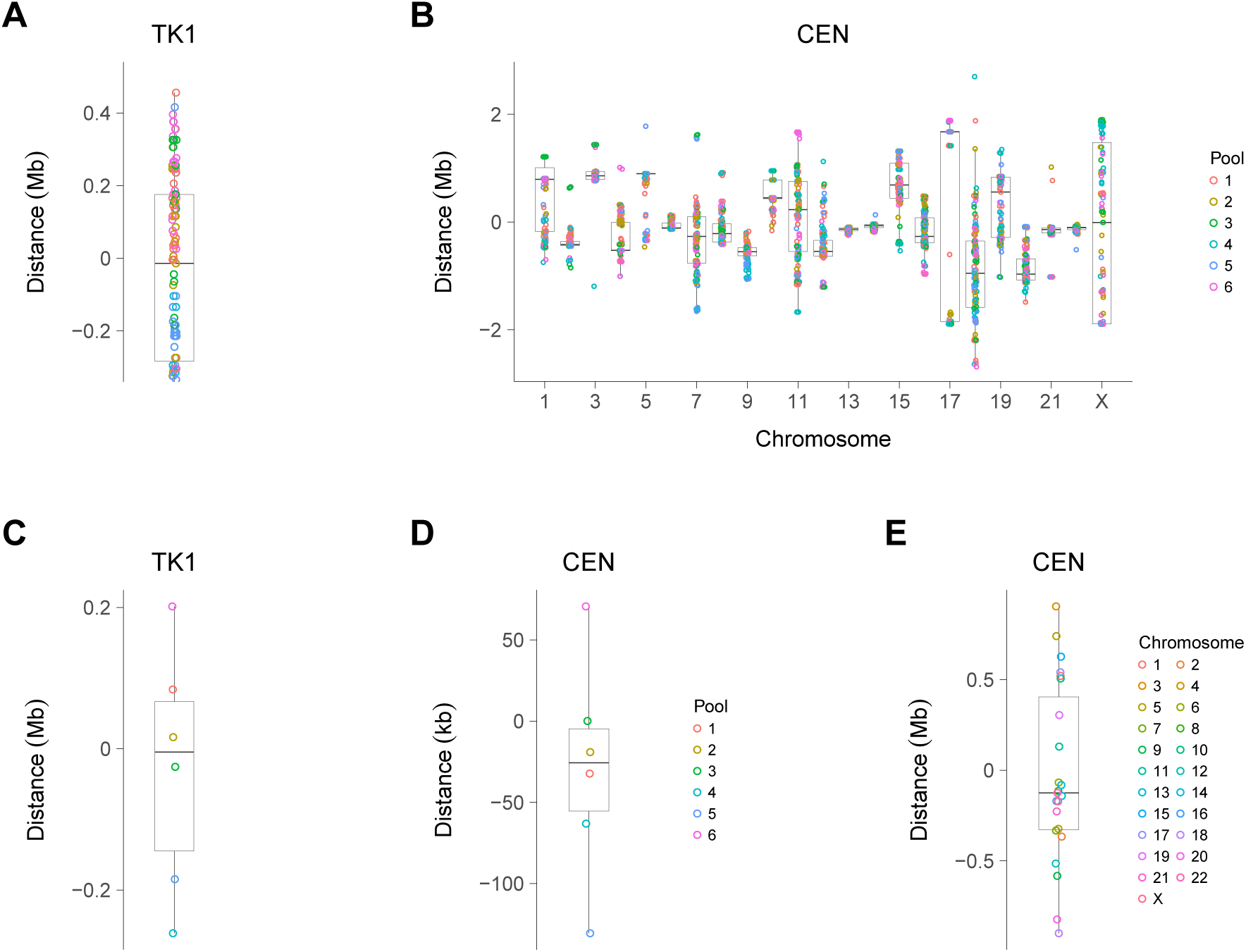
Mapping resolution. (**A**) Distance between genome location of TK1 and its retention peak. (**B**) Distance between assigned centromere genome locations and retention peaks. (**C**) Distance between TK1 and its retention peak averaged across the six RH lineages. (**D**) Distance between assigned centromere locations and retention peaks averaged across the six RH lineages. (**E**) Distance between assigned centromere locations and retention peaks averaged across chromosomes.

**Figure S35.**
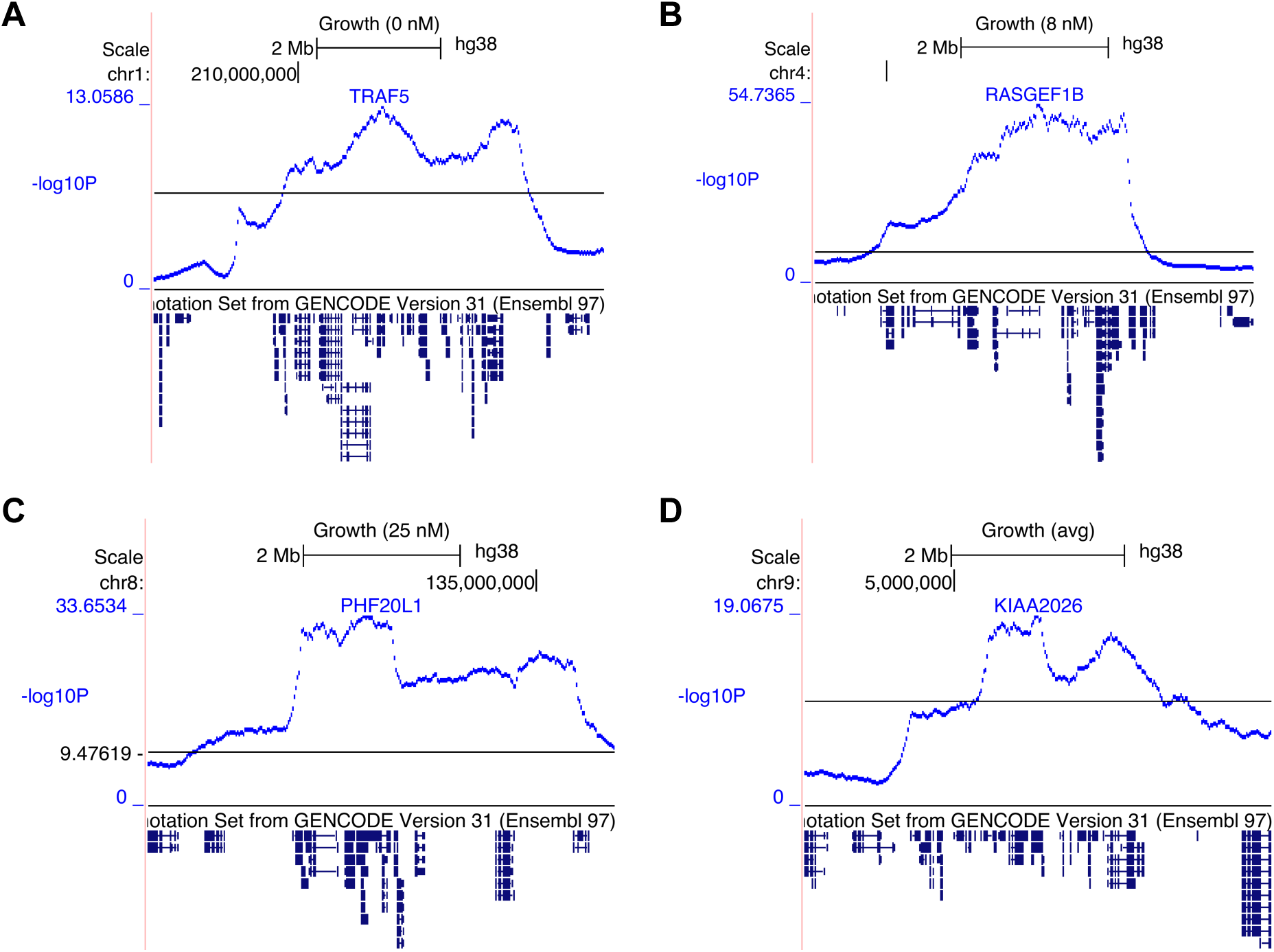
Regions of high gene density. (**A**) TRAF5. (**B**) RASGEF1B. (**C**) PHF20L1. (**D**) KIAA2026. In some regions, mapping resolution was insufficient to distinguish between closely packed genes. Genes with peak −log_10_ *P* values shown. Horizontal black lines, permutation significance thresholds.

**Figure S36.**
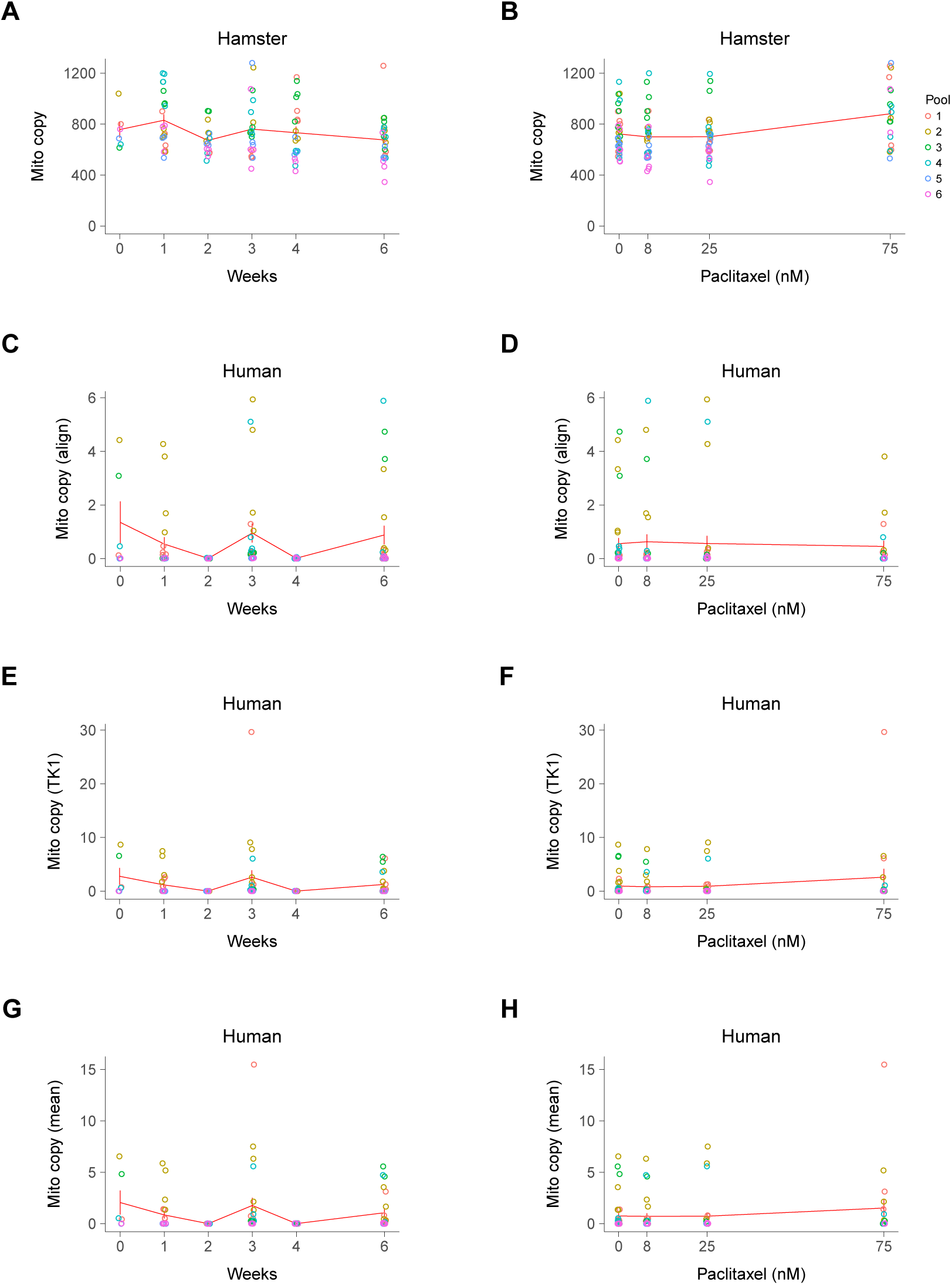
Mitochondrial copy number. (**A**) Hamster mitochrondrial copy number vs. growth time. (**B**) Hamster mitochondrial copy number vs. paclitaxel concentration. (**C**) Human mitochrondrial copy number based on sequence alignments vs. growth time. (**D**) Human mitochrondrial copy number based on sequence alignments vs. paclitaxel concentration. (**E**) Human mitochrondrial copy number based on TK1 corrected for revertants vs. growth time. (**F**) Human mitochrondrial copy number based on TK1 corrected for revertants vs. paclitaxel concentration. (**G**) Human mitochrondrial copy number based on mean of sequence alignments and TK1 corrected for revertants vs. growth time. (**H**) Human mitochrondrial copy number based on mean of sequence alignments and TK1 corrected for revertants vs. paclitaxel concentration. Means ± s.e.m.

**Figure S37.**
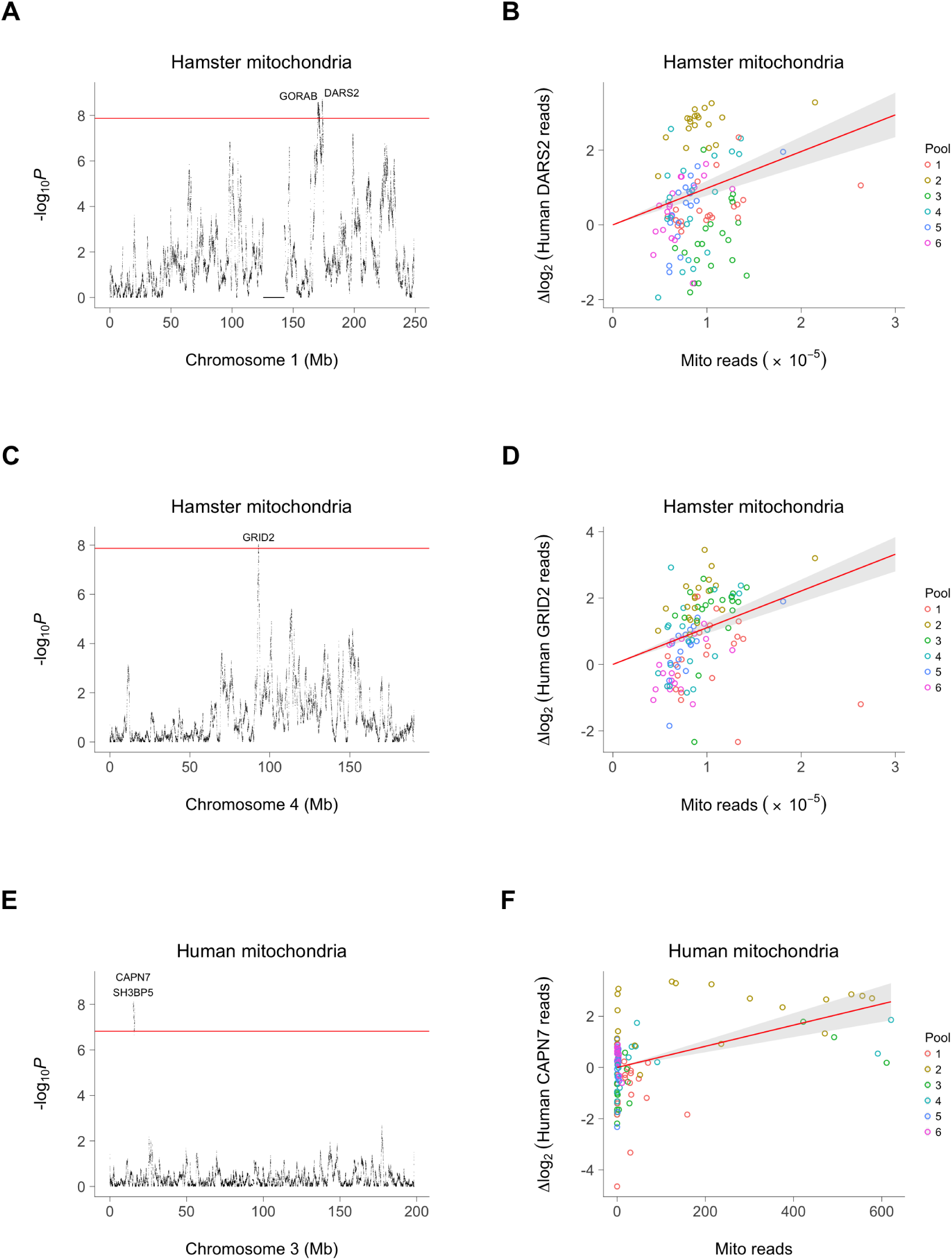
Loci affecting mitochondrial copy number. (**A**) GORAB and DARS2 on human chromosome 1 regulate hamster mitochondrial copy number. (**B**) Increase in normalized log_2_ human DARS2 sequence reads associated with an increase in hamster mitochondrial reads. Red line, best fit. Grey band, 95% confidence interval. (**C**) GRID2 locus on human chromosome 4 regulates hamster mitochondrial copy number. (**D**) Increase in human GRID2 sequence reads associated with increase in hamster mitochondria. (**E**) Locus on human chromosome 3 mapping to CAPNN7 and near SH3BP5 regulates human mitochondrial copy number. (**F**) Increase in human CAPNN7 sequence reads associated with increase in human mitochondria. Horizontal red lines (A, C), permutation significance thresholds; (E), FDR = 0.05.

**Figure S38.**
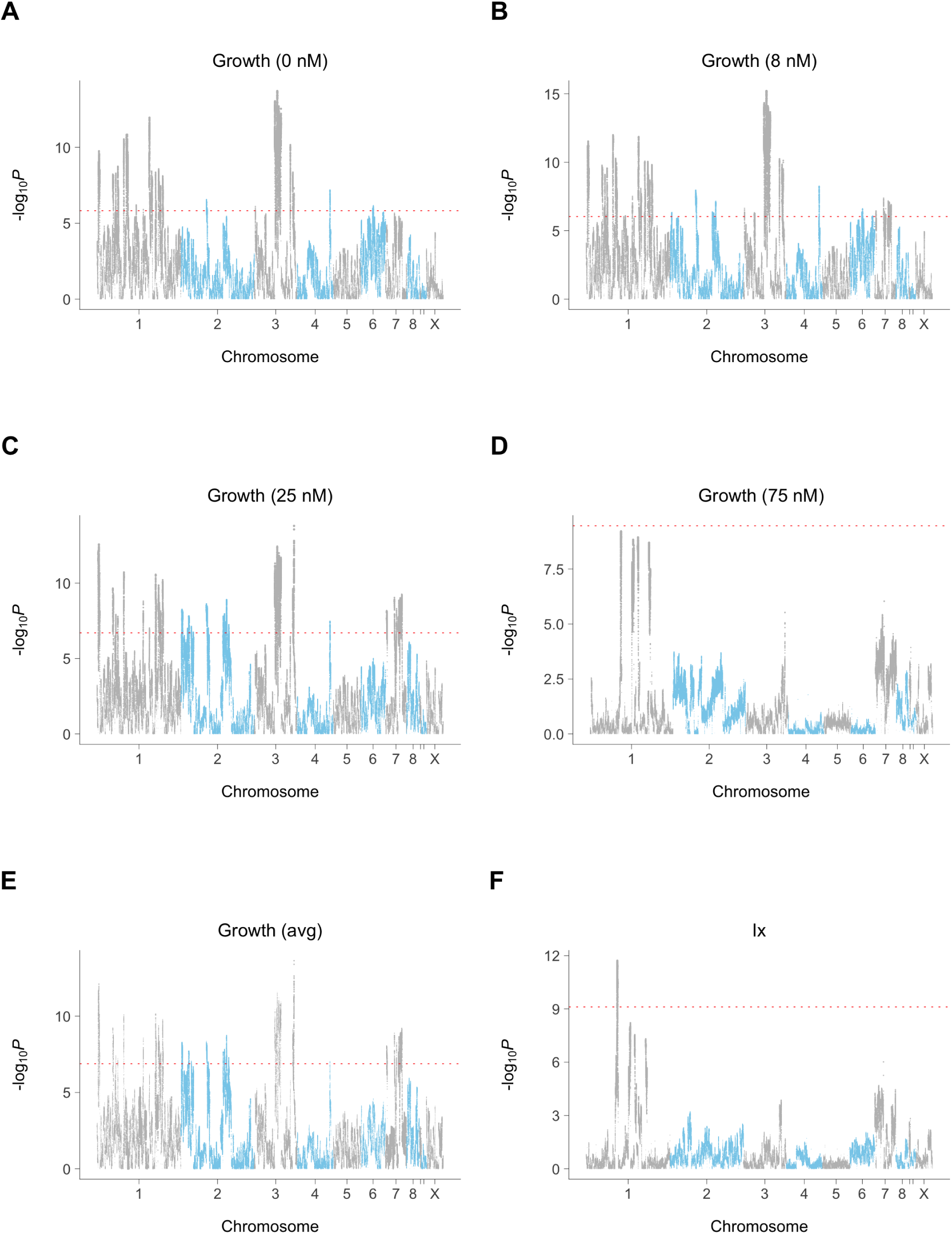
Hamster growth loci. (**A**) 0 nM paclitaxel. (**B**) 8 nM paclitaxel. (**C**) 25 nM paclitaxel. (**D**) 75 nM paclitaxel. (**E**) Average conditional effect of growth. (**F**) Growth/paclitaxel interaction. Red dotted line, permutation significance threshold.

**Figure S39.**
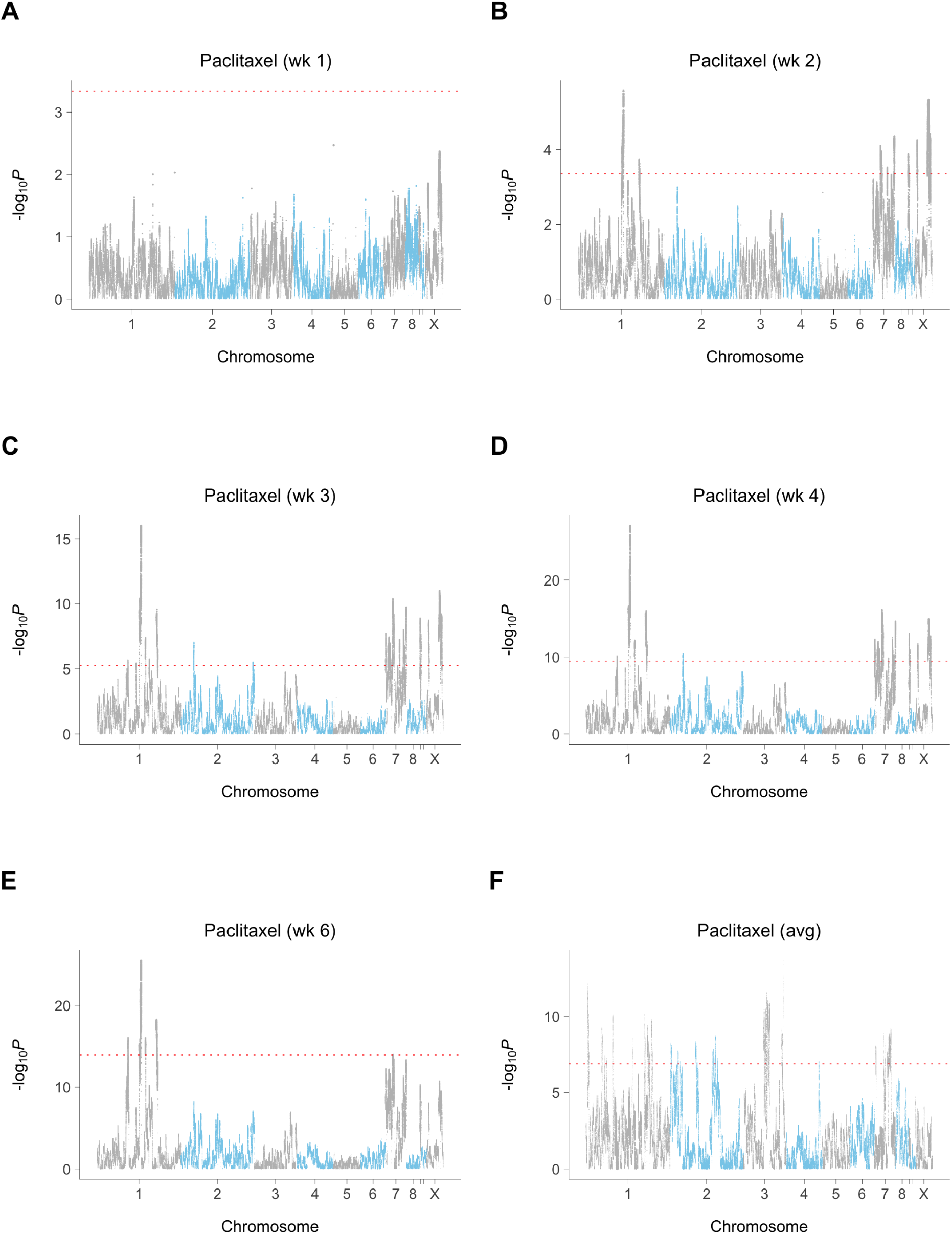
Hamster paclitaxel loci. (**A**) Week 1. (**B**) Week 2. (**C**) Week 3. (**D**) Week 4. (**E**) Week 6. (**F**) Average conditional effect of paclitaxel. Red dotted line, permutation significance threshold.

**Figure S40.**
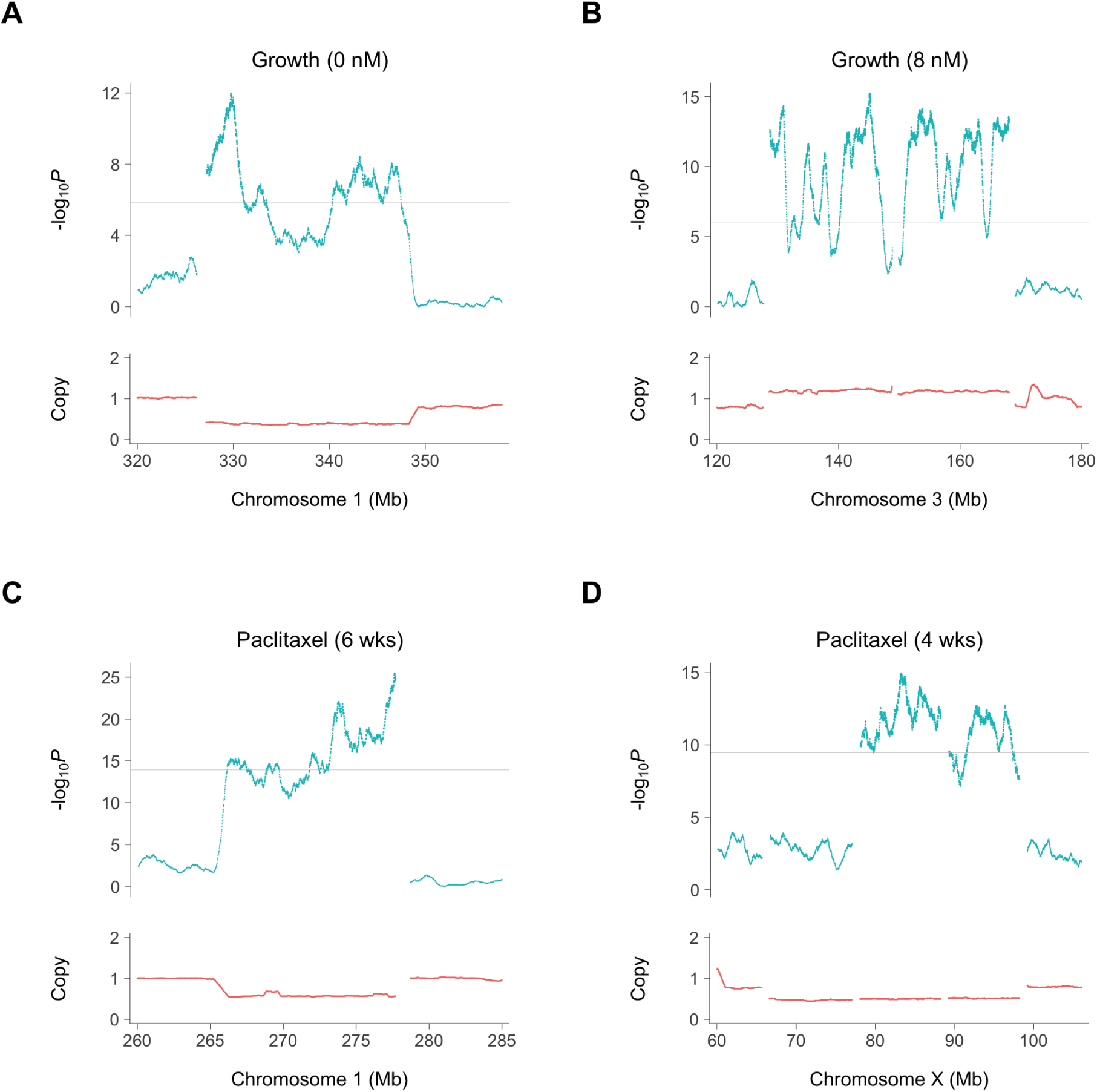
Hamster growth and paclitaxel loci due to pre-existing copy number alterations (CNAs). (**A**) Growth (0 nM paclitaxel). (**B**) Growth (8 nM paclitaxel). (**C**) Paclitaxel (week 6). (**D**) Paclitaxel (week 4). Horizontal gray lines, permutation significance thresholds. Relative copy numbers are for hamster reads averaged across the six RH pools. Breaks represent contig transitions.

**Table S1.**
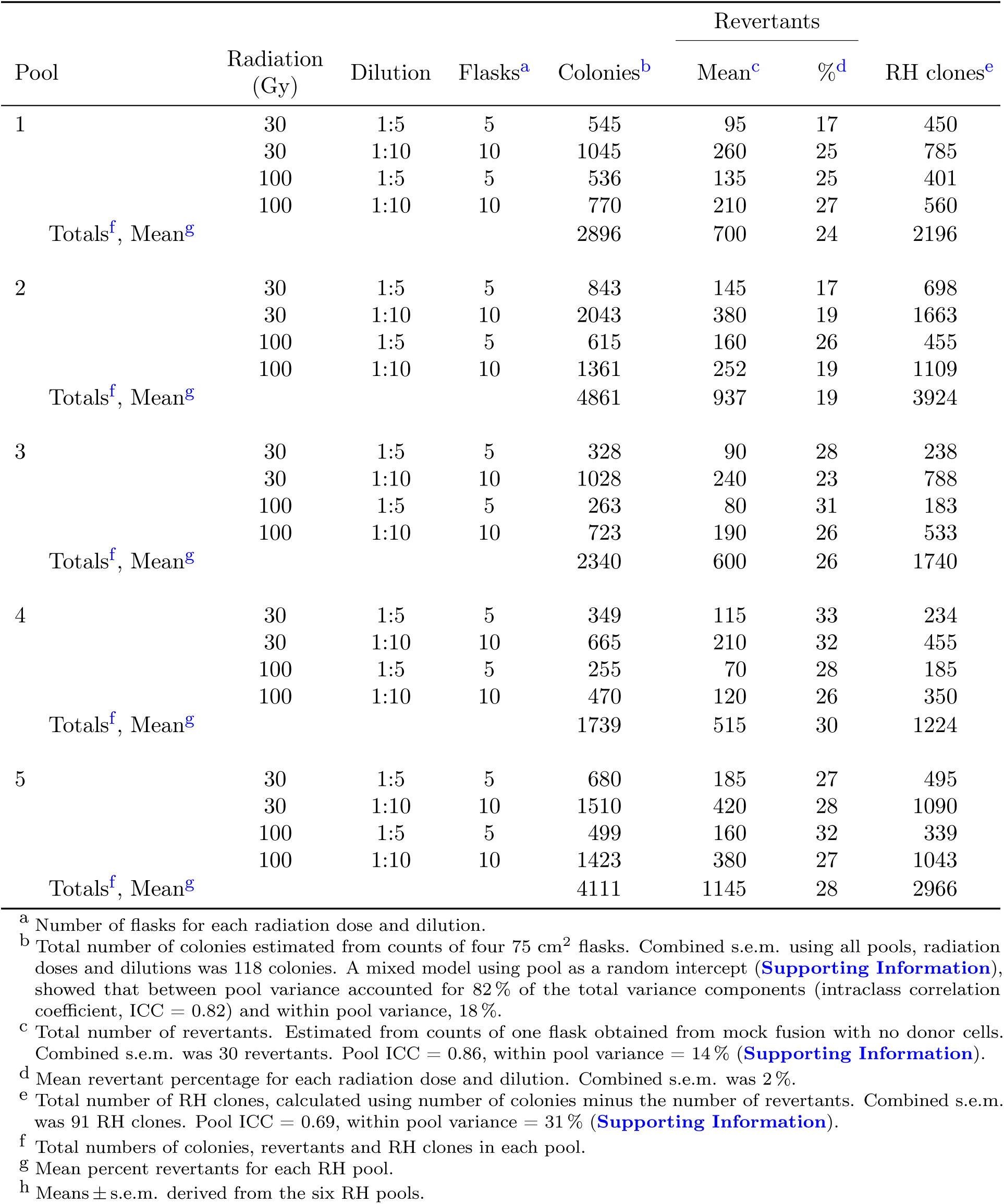
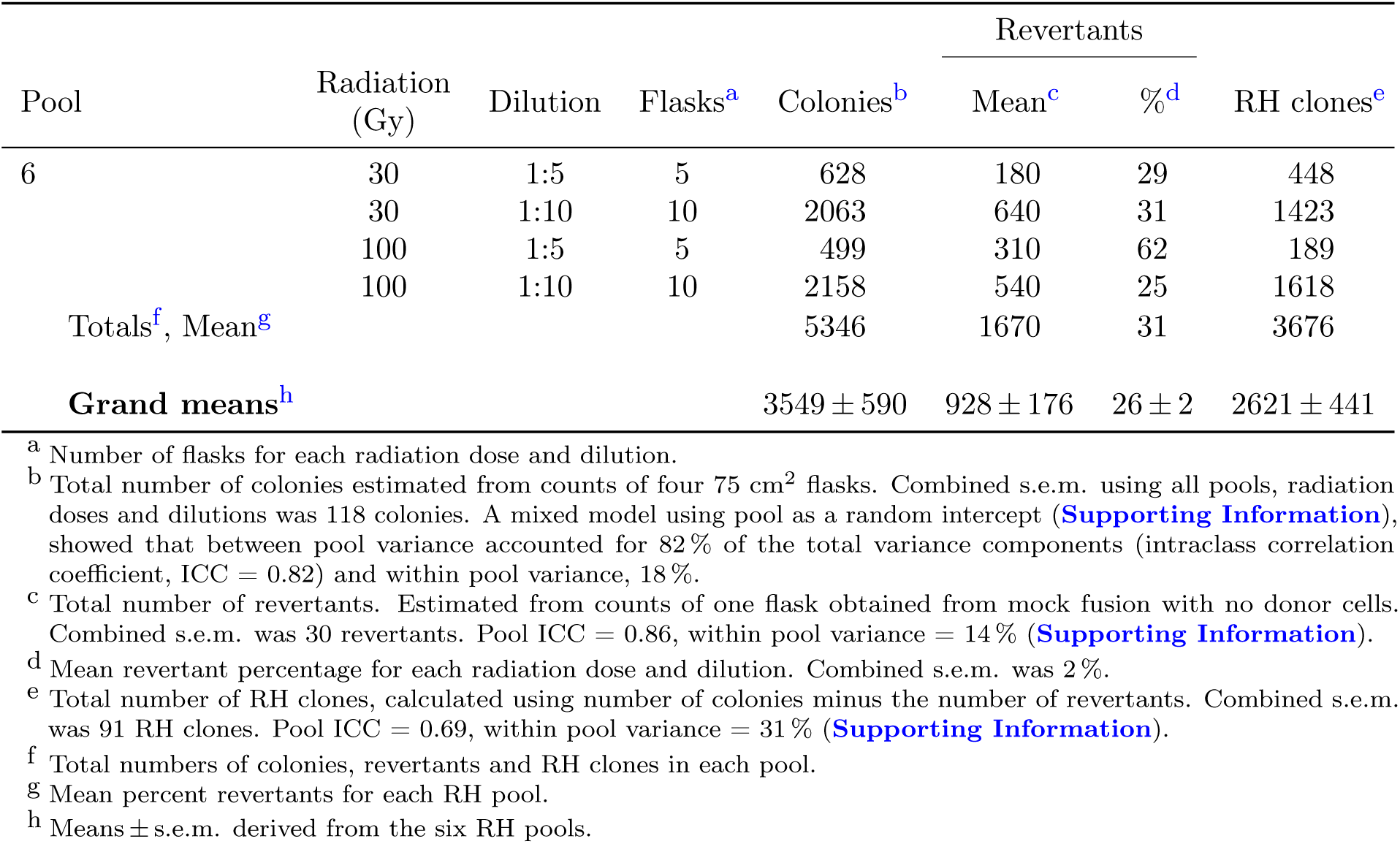
Clone numbers

| Pool | Radiation<br>(Gy) | Dilution | Flasks<sup>a</sup> | Colonies<sup>b</sup> | Revertants |  | RH clones<sup>e</sup> |
| --- | --- | --- | --- | --- | --- | --- | --- |
|  |  |  |  |  | Mean<sup>c</sup> | %<sup>d</sup> |  |
| 1 | 30 | 1:5 | 5 | 545 | 95 | 17 | 450 |
|  | 30 | 1:10 | 10 | 1045 | 260 | 25 | 785 |
|  | 100 | 1:5 | 5 | 536 | 135 | 25 | 401 |
|  | 100 | 1:10 | 10 | 770 | 210 | 27 | 560 |
|  | Totals<sup>f</sup>, Mean<sup>g</sup> |  |  | 2896 | 700 | 24 | 2196 |
| 2 | 30 | 1:5 | 5 | 843 | 145 | 17 | 698 |
|  | 30 | 1:10 | 10 | 2043 | 380 | 19 | 1663 |
|  | 100 | 1:5 | 5 | 615 | 160 | 26 | 455 |
|  | 100 | 1:10 | 10 | 1361 | 252 | 19 | 1109 |
|  | Totals<sup>f</sup>, Mean<sup>g</sup> |  |  | 4861 | 937 | 19 | 3924 |
| 3 | 30 | 1:5 | 5 | 328 | 90 | 28 | 238 |
|  | 30 | 1:10 | 10 | 1028 | 240 | 23 | 788 |
|  | 100 | 1:5 | 5 | 263 | 80 | 31 | 183 |
|  | 100 | 1:10 | 10 | 723 | 190 | 26 | 533 |
|  | Totals<sup>f</sup>, Mean<sup>g</sup> |  |  | 2340 | 600 | 26 | 1740 |
| 4 | 30 | 1:5 | 5 | 349 | 115 | 33 | 234 |
|  | 30 | 1:10 | 10 | 665 | 210 | 32 | 455 |
|  | 100 | 1:5 | 5 | 255 | 70 | 28 | 185 |
|  | 100 | 1:10 | 10 | 470 | 120 | 26 | 350 |
|  | Totals<sup>f</sup>, Mean<sup>g</sup> |  |  | 1739 | 515 | 30 | 1224 |
| 5 | 30 | 1:5 | 5 | 680 | 185 | 27 | 495 |
|  | 30 | 1:10 | 10 | 1510 | 420 | 28 | 1090 |
|  | 100 | 1:5 | 5 | 499 | 160 | 32 | 339 |
|  | 100 | 1:10 | 10 | 1423 | 380 | 27 | 1043 |
|  | Totals<sup>f</sup>, Mean<sup>g</sup> |  |  | 4111 | 1145 | 28 | 2966 |
|  |  |  |  |  | Mean<sup>c</sup> | %<sup>d</sup> |  |
| 6 | 30 | 1:5 | 5 | 628 | 180 | 29 | 448 |
|  | 30 | 1:10 | 10 | 2063 | 640 | 31 | 1423 |
|  | 100 | 1:5 | 5 | 499 | 310 | 62 | 189 |
|  | 100 | 1:10 | 10 | 2158 | 540 | 25 | 1618 |
| Totals<sup>f</sup>, Mean<sup>g</sup> |  |  |  | 5346 | 1670 | 31 | 3676 |
| **Grand means<sup>h</sup>** |  |  |  | 3549 ± 590 | 928 ± 176 | 26 ± 2 | 2621 ± 441 |

<sup>a</sup> Number of flasks for each radiation dose and dilution. <sup>b</sup> Total number of colonies estimated from counts of four 75 cm<sup>2</sup> flasks. Combined s.e.m. using all pools, radiation doses and dilutions was 118 colonies. A mixed model using pool as a random intercept (**Supporting Information**), showed that between pool variance accounted for 82 % of the total variance components (intraclass correlation coefficient, ICC = 0.82) and within pool variance, 18 %. <sup>c</sup> Total number of revertants. Estimated from counts of one flask obtained from mock fusion with no donor cells. Combined s.e.m. was 30 revertants. Pool ICC = 0.86, within pool variance = 14 % (**Supporting Information**). <sup>d</sup> Mean revertant percentage for each radiation dose and dilution. Combined s.e.m. was 2 %. <sup>e</sup> Total number of RH clones, calculated using number of colonies minus the number of revertants. Combined s.e.m. was 91 RH clones. Pool ICC = 0.69, within pool variance = 31 % (**Supporting Information**). <sup>f</sup> Total numbers of colonies, revertants and RH clones in each pool. <sup>g</sup> Mean percent revertants for each RH pool. <sup>h</sup> Means ± s.e.m. derived from the six RH pools.

**Table S2.** Experimental design^a^

| Weeks | Paclitaxel<br>(nM) |  |  |  |
| --- | --- | --- | --- | --- |
|  | 0 | 8 | 25 | 75 |
| 0 | 6<sup>b</sup> | - | - | - |
| 1 | 6 | 6 | 6 | 4 |
| 2 | 6 | 6 | 6 | - |
| 3 | 6 | 6 | 6 | 6 |
| 4 | 6 | 6 | 6 | 3 |
| 6 | 6 | 6 | 6 | 6 |

<sup>a</sup> Number of samples analyzed. <sup>b</sup> The RH pools.

**Table S3.**
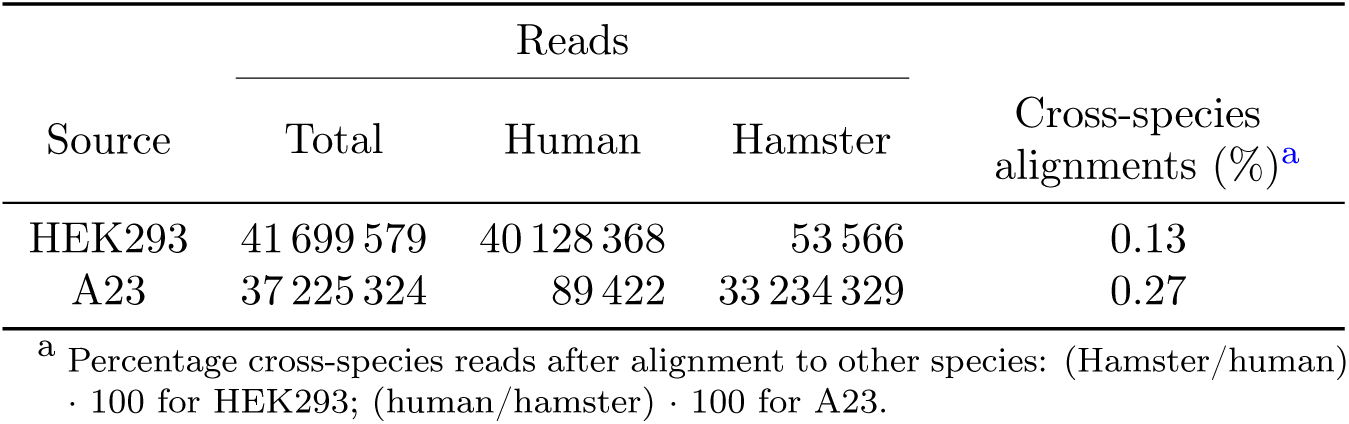
Genome alignments and misalignments

**Table S4.**
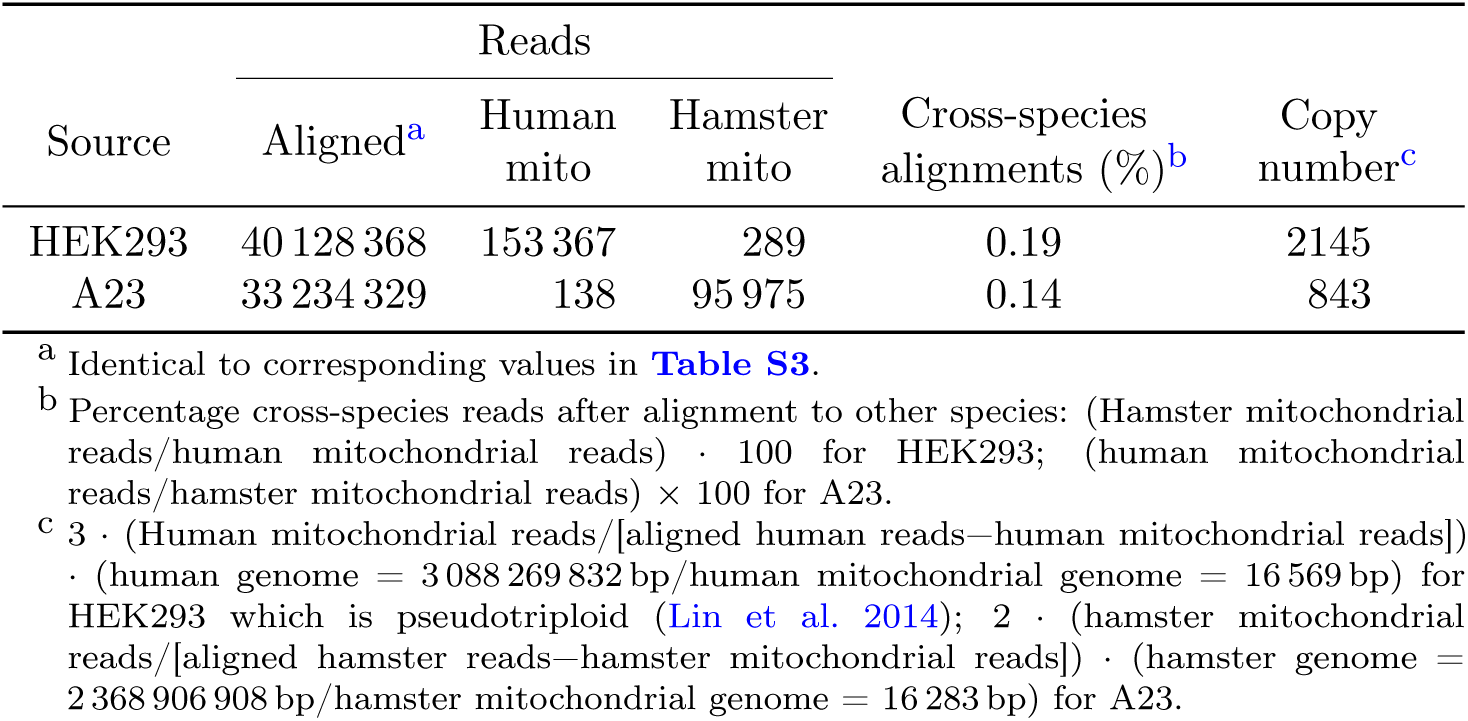
Mitochondrial alignments and misalignments

| Source | Reads |  |  | Cross-species alignments (%)<sup>b</sup> | Copy number<sup>c</sup> |
| --- | --- | --- | --- | --- | --- |
|  | Aligned<sup>a</sup> | Human mito | Hamster mito |  |  |
| HEK293 | 40 128 368 | 153 367 | 289 | 0.19 | 2145 |
| A23 | 33 234 329 | 138 | 95 975 | 0.14 | 843 |

<sup>a</sup> Identical to corresponding values in **Table S3**.
<sup>b</sup> Percentage cross-species reads after alignment to other species: (Hamster mitochondrial reads/human mitochondrial reads) · 100 for HEK293; (human mitochondrial reads/hamster mitochondrial reads) · 100 for A23.
<sup>c</sup> $3 \cdot (\text{Human mitochondrial reads} / [\text{aligned human reads} - \text{human mitochondrial reads}]) \cdot (\text{human genome} = 3\,088\,269\,832 \text{ bp} / \text{human mitochondrial genome} = 16\,569 \text{ bp})$ for HEK293 which is pseudotriploid (Lin et al. 2014); $2 \cdot (\text{hamster mitochondrial reads} / [\text{aligned hamster reads} - \text{hamster mitochondrial reads}]) \cdot (\text{hamster genome} = 2\,368\,906\,908 \text{ bp} / \text{hamster mitochondrial genome} = 16\,283 \text{ bp})$ for A23.

**Table S5.** Human genome aligments in the RH pools

| Pool | Reads |  |  |  | Cross-species reads (%)<sup>d</sup> |
| --- | --- | --- | --- | --- | --- |
|  | Total | Human alignments<sup>a</sup> | Corrected human<sup>b</sup> | Both species<sup>c</sup> |  |
| 1 | 42 000 172 | 398 642 | 298 433 | 100 209 | 25.1 |
| 2 | 35 358 581 | 868 990 | 785 571 | 83 419 | 9.6 |
| 3 | 52 166 656 | 1 173 763 | 1 063 450 | 110 313 | 9.4 |
| 4 | 32 756 205 | 1 121 927 | 1 048 516 | 73 411 | 6.5 |
| 5 | 42 876 176 | 853 173 | 638 622 | 214 551 | 25.1 |
| 6 | 41 449 894 | 842 565 | 632 519 | 210 046 | 24.9 |
| **Mean** | 41 101 281 | 876 510 | 744 518 | 131 992 | 16.8 |
| **s.e.m.** | 2 763 700 | 112 354 | 118 120 | 25 936 | 3.7 |

<sup>a</sup> Aligned to human genome.
<sup>b</sup> Human reads minus those that also align to hamster genome.
<sup>c</sup> Aligned to both human and hamster.
<sup>d</sup> (Aligned to both species/human alignments) · 100.

**Table S6.** Hamster genome aligments in the RH pools

| Pool | Reads |  |  |  | Cross-species reads (%)<sup>e</sup> |
| --- | --- | --- | --- | --- | --- |
|  | Total<sup>a</sup> | Hamster alignments<sup>b</sup> | Corrected hamster<sup>c</sup> | Both species<sup>d</sup> |  |
| 1 | 42 000 172 | 37 695 387 | 37 595 178 | 100 209 | 0.27 |
| 2 | 35 358 581 | 30 838 307 | 30 754 890 | 83 417 | 0.27 |
| 3 | 52 166 656 | 45 779 128 | 45 668 817 | 110 311 | 0.24 |
| 4 | 32 756 205 | 28 166 022 | 28 092 611 | 73 411 | 0.26 |
| 5 | 42 876 176 | 38 382 035 | 38 167 486 | 214 549 | 0.56 |
| 6 | 41 449 894 | 37 403 393 | 37 193 349 | 210 044 | 0.56 |
| **Mean** | 41 101 281 | 36 377 379 | 36 245 388 | 131 990 | 0.36 |
| **s.e.m.** | 2 763 700 | 2 538 432 | 2 528 698 | 25 935 | 0.06 |

<sup>a</sup> Identical to same column in **Table S5**. <sup>b</sup> Aligned to hamster genome. <sup>c</sup> Hamster reads minus those that also align to human genome. <sup>d</sup> Aligned to both human and hamster. Identical to same column in **Table S5**, apart from minor alignment differences. <sup>e</sup> (Aligned to both species/hamster alignments) · 100.

Table S7. Human DNA retention in the RH pools

| Pool | Reads<sup>a</sup> |  |  | Revert freq | Retention |  |  |
| --- | --- | --- | --- | --- | --- | --- | --- |
|  | Human reads<sup>b</sup> | Hamster reads<sup>b</sup> | Human TK1<sup>c</sup> |  | align (%)<sup>d</sup> | TK1 (%)<sup>e</sup> | mean (%) |
| 1 | 76 | 15 130 | 997 | 0.24 | 1.00 | 5.82 | 3.41 |
| 2 | 194 | 12 372 | 2636 | 0.20 | 3.14 | 5.88 | 4.51 |
| 3 | 246 | 18 255 | 3302 | 0.27 | 2.70 | 5.45 | 4.07 |
| 4 | 259 | 11 237 | 3125 | 0.29 | 4.61 | 5.85 | 5.23 |
| 5 | 155 | 15 082 | 2234 | 0.28 | 2.06 | 4.96 | 3.51 |
| 6 | 154 | 14 877 | 2573 | 0.37 | 2.07 | 3.79 | 2.93 |
| **Mean** | 181 | 14 492 | 2478 | 0.28 | 2.60 | 5.29 | 3.94 |
| **s.e.m.** | 28 | 1003 | 336 | 0.02 | 0.50 | 0.33 | 0.34 |

<sup>a</sup> All reads purged of cross-species alignments. <sup>b</sup> Medians of non-overlapping 1 Mb windows. <sup>c</sup> Maximum 1 Mb window encompassing TK1. <sup>d</sup> $2 \cdot (\text{Human reads} / \text{hamster reads}) \cdot 100$ . <sup>e</sup> $(\text{Human reads} / \text{TK1}) \cdot (1 - \text{reversion frequency}) \cdot 100$ .

**Table S8.** Hamster-specific reads in RH samples. (**A**) Contig identification (ID) number. (**B**) Chromosome. (**C**) Window start position. (**D**) Window stop position. (**E**) Average (midpoint) window position. (**F** to **DP**) RH1_d0_w0 refers to RH pool 1 at 0 nM paclitaxel (drug), week 0, etc. Windows of 1 Mb, 10 kb steps.

**Table S9.** Human-specific reads in RH samples. (**A**) Chromosome. (**B**) Window start position. (**C**) Window stop position. (**D**) Average (midpoint) window position. (**E**–**DO**) RH1_d0_w0 refers to RH pool 1 at 0 nM paclitaxel (drug), week 0, etc. Windows of 1 Mb, 10 kb steps.

**Table S10.** Significance values in human genome for growth, paclitaxel and interaction loci. (**A**) Chromosome. (**B**) Window start position. (**C**) Window stop position. (**D**) Average (midpoint) window position. (**E**–**H**) log10p_g_0nM refers to −log_10_ *P* values for growth at 0 nM paclitaxel, etc. (**I**) log10p_g_avg refers to −log_10_ *P* values for average conditional effect of growth. (**J**–**N**) log10p_d_w1 refers to −log_10_ *P* values for paclitaxel (drug) action at week 1, etc. (**O**) log10p_d_avg refers to −log_10_ *P* values for average conditional effect of paclitaxel. (**P**) log10p_g_d_Ix refers to −log_10_ *P* values for interaction between growth time and paclitaxel concentration. (**Q**–**AB**) Coefficients of slope for corresponding −log_10_ *P* values in (E–P). Windows of 1 Mb, 10 kb steps.

**Table S11.** Significance thresholds in human genome for growth, paclitaxel and interaction loci. (**A**) Genome scans. log10p_g_0nM refers to −log_10_ *P* values for growth at 0 nM paclitaxel, etc., log10p_g_avg refers to −log_10_ *P* values for average conditional effect of growth, log10p_d_w1 refers to −log_10_ *P* values for paclitaxel (drug) action at week 1, etc., log10p_d_avg refers to −log_10_ *P* values for average conditional effect of paclitaxel, log10p_g_d_Ix refers to −log_10_ *P* values for interaction between growth time and paclitaxel concentration. (**B**) Significance thresholds at 5% family-wise error rate (FWER) obtained using permutation.

**Table S12.** Growth loci numbers

| Paclitaxel (nM) | Number<sup>a</sup> |
| --- | --- |
| 0 | 409 |
| 8 | 460 |
| 25 | 466 |
| 75 | 55 |
| Avg<sup>b</sup> | 446 |
| Total | 1836 |
| Non-redundant | 859 |

<sup>a</sup> Exceeding permutation significance threshold. <sup>b</sup> Average conditional effect.

**Table S13.** Growth loci. (**A**) Chromosome. (**B**) Window start position. (**C**) Window stop position. (**D**) Average (midpoint) window position. (**E**) Paclitaxel concentration of growth locus. “avg” means average conditional effect. (**F**) Significance of locus measured by −log_10_ *P*. (**G**) Coefficient of slope for locus. (**H**) Distance (bp) between locus peak and closest end of nearest gene (5′ or 3′). (**I**) ENSEMBL gene ID. (**J**) ENSEMBL transcript ID. (**K**) Nearest gene symbol. (**L**) Sense strand. (**M**) Gene start coordinates. (**N**) Gene end coordinates. (**0**) Gene length. (**P**) Longest transcript length. (**Q**) Coding sequence length. (**R**) 5′ untranslated region start coordinates. (**S**) 5′ untranslated region end coordinates. (**T**) 5′ untranslated region length. (**U**) 3′ untranslated region start coordinates. (**V**) 3′ untranslated region end coordinates. (**W**) 3′ untranslated region length. 5′ and 3′ untranslated regions are longest alternatives. (**X**) Exon count. (**Y**) Gene type. (**Z**) Gene description. (**AA** to **AD**) log10p_g_0nM refers to −log_10_ *P* values for growth at 0 nM paclitaxel, etc. (**AE**) log10p_g_avg refers to −log_10_ *P* values for average conditional effect of growth. (**AF**–**AJ**) log10p_d_w1 refers to −log_10_ *P* values for paclitaxel (drug) action at week 1, etc. (**AK**) log10p_d_avg refers to −log_10_ *P* values for average conditional effect of paclitaxel. (**AL**) log10p_g_d_Ix refers to −log_10_ *P* values for interaction between growth time and paclitaxel concentration. (**AM**–**AX**) Coefficients of slope for corresponding −log_10_ *P* values in (AA–AL).

**Table S14.** Correlations of −log_10_ *P* values

|  |  | Growth<br>(nM) |  |  |  |  | Paclitaxel<br>(weeks) |  |  |  |  |  |  |
| --- | --- | --- | --- | --- | --- | --- | --- | --- | --- | --- | --- | --- | --- |
|  |  | 0 | 8 | 25 | 75 | avg<sup>a</sup> | 1 | 2 | 3 | 4 | 6 | avg<sup>a</sup> | Ix |
| Growth (nM) | 0 | 1.00 | 0.99 | 0.88 | 0.21 | 0.86 | 0.23 | 0.17 | 0.01 | −0.05 | −0.10 | −0.01 | −0.16 |
|  | 8 | 0.99 | 1.00 | 0.94 | 0.32 | 0.93 | 0.24 | 0.11 | −0.04 | −0.09 | −0.13 | −0.05 | −0.17 |
|  | 25 | 0.88 | 0.94 | 1.00 | 0.59 | 1.00 | 0.24 | 0.02 | −0.06 | −0.07 | −0.08 | −0.06 | −0.06 |
|  | 75 | 0.21 | 0.32 | 0.59 | 1.00 | 0.62 | 0.12 | 0.05 | 0.27 | 0.38 | 0.47 | 0.30 | 0.60 |
|  | avg | 0.86 | 0.93 | 1.00 | 0.62 | 1.00 | 0.24 | 0.02 | −0.05 | −0.06 | −0.06 | −0.05 | −0.04 |
| Pacli (wks) | 1 | 0.23 | 0.24 | 0.24 | 0.12 | 0.24 | 1.00 | 0.47 | 0.11 | 0.01 | −0.02 | 0.08 | −0.01 |
|  | 2 | 0.17 | 0.11 | 0.02 | 0.05 | 0.02 | 0.47 | 1.00 | 0.83 | 0.70 | 0.59 | 0.80 | 0.37 |
|  | 3 | 0.01 | −0.04 | −0.06 | 0.27 | −0.05 | 0.11 | 0.83 | 1.00 | 0.97 | 0.91 | 1.00 | 0.73 |
|  | 4 | −0.05 | −0.09 | −0.07 | 0.38 | −0.06 | 0.01 | 0.70 | 0.97 | 1.00 | 0.97 | 0.99 | 0.83 |
|  | 6 | −0.10 | −0.13 | −0.08 | 0.47 | −0.06 | −0.02 | 0.59 | 0.91 | 0.97 | 1.00 | 0.93 | 0.93 |
|  | avg | −0.01 | −0.05 | −0.06 | 0.30 | −0.05 | 0.08 | 0.80 | 1.00 | 0.99 | 0.93 | 1.00 | 0.76 |
| Interaction | - | −0.16 | −0.17 | −0.06 | 0.60 | −0.04 | −0.01 | 0.37 | 0.73 | 0.83 | 0.93 | 0.76 | 1.00 |

<sup>a</sup> Average conditional effect

**Table S15.** Overlap of non-unique loci

|  |  | Growth<br>(nM) |  |  |  |  | Paclitaxel<br>(weeks) <sup>a</sup> |  |  |  |  | Ix |
| --- | --- | --- | --- | --- | --- | --- | --- | --- | --- | --- | --- | --- |
|  |  | 0 | 8 | 25 | 75 | avg<sup>b</sup> | 2 | 3 | 4 | 6 | avg<sup>b</sup> |  |
| Growth (nM) | 0 | 409 | 305 | 162 | 3 | 142 | 1 | 4 | 2 | 2 | 4 | 2 |
|  | 8 | 305 | 460 | 238 | 5 | 207 | 0 | 0 | 0 | 0 | 0 | 0 |
|  | 25 | 162 | 238 | 466 | 16 | 414 | 0 | 0 | 0 | 0 | 0 | 2 |
|  | 75 | 3 | 5 | 16 | 55 | 16 | 0 | 4 | 4 | 7 | 4 | 15 |
|  | avg<sup>b</sup> | 142 | 207 | 414 | 16 | 446 | 0 | 0 | 0 | 0 | 0 | 2 |
| Pacli (wks) <sup>a</sup> | 2 | 1 | 0 | 0 | 0 | 0 | 2 | 2 | 2 | 2 | 2 | 0 |
|  | 3 | 4 | 0 | 0 | 4 | 0 | 2 | 26 | 17 | 15 | 23 | 7 |
|  | 4 | 2 | 0 | 0 | 4 | 0 | 2 | 17 | 19 | 15 | 18 | 8 |
|  | 6 | 2 | 0 | 0 | 7 | 0 | 2 | 15 | 15 | 25 | 16 | 14 |
|  | avg<sup>b</sup> | 4 | 0 | 0 | 4 | 0 | 2 | 23 | 18 | 16 | 25 | 8 |
| Interaction | - | 2 | 0 | 2 | 15 | 2 | 0 | 7 | 8 | 14 | 8 | 62 |

<sup>a</sup> No significant paclitaxel loci for week 1.
<sup>b</sup> Average conditional effect

**Table S16.** Non-redundant growth loci. (**A**) Chromosome. (**B**) Window start position. (**C**) Window stop position. (**D**) Average (midpoint) window position. (**E**) Paclitaxel concentration of growth locus with maximum −log_10_ *P*. “avg” means average conditional effect. (**F**) Significance of locus measured by −log_10_ *P*. (**G**) Coefficient of slope for locus. (**H**) Distance (bp) between locus peak and closest end of nearest gene (5′ or 3′). (**I**) ENSEMBL gene ID. (**J**) ENSEMBL transcript ID. (**K**) Nearest gene symbol. (**L**) Sense strand. (**M**) Gene start coordinates. (**N**) Gene end coordinates. (**0**) Gene length. (**P**) Longest transcript length. (**Q**) Coding sequence length. (**R**) 5′ untranslated region start coordinates. (**S**) 5*^I^* untranslated region end coordinates. (**T**) 5′ untranslated region length. (**U**) 3′ untranslated region start coordinates. (**V**) 3*^I^* untranslated region end coordinates. (**W**) 3*^I^* untranslated region length. 5′ and 3′ untranslated regions are longest alternatives. (**X**) Exon count. (**Y**) Gene type. (**Z**) Gene description. (**AA** to **AD**) log10p_g_0nM refers to −log_10_ *P* values for growth at 0 nM paclitaxel, etc. (**AE**) log10p_g_avg refers to −log_10_ *P* values for average conditional effect of growth. (**AF**–**AJ**) log10p_d_w1 refers to −log_10_ *P* values for paclitaxel (drug) action at week 1, etc. (**AK**) log10p_d_avg refers to −log_10_ *P* values for average conditional effect of paclitaxel. (**AL**) log10p_g_d_Ix refers to −log_10_ *P* values for interaction between growth time and paclitaxel concentration. (**AM**–**AX**) Coefficients of slope for corresponding −log_10_ *P* values in (AA–AL).

**Table S17.**
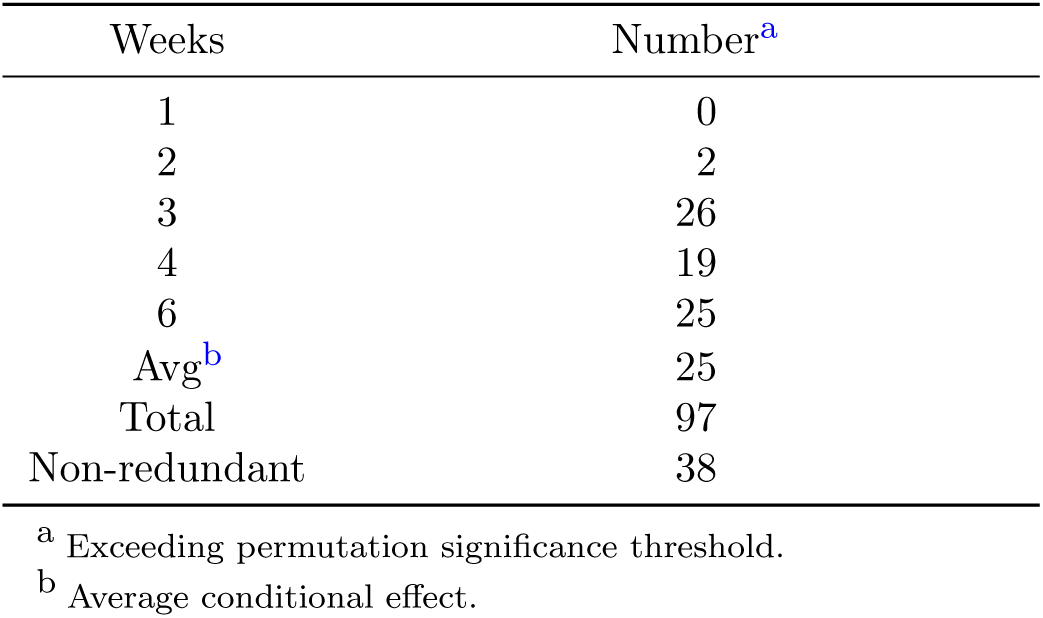
Paclitaxel loci numbers

**Table S18.** Paclitaxel loci. (**A**) Chromosome. (**B**) Window start position. (**C**) Window stop position. (**D**) Average (midpoint) window position. (**E**) Growth week of paclitaxel locus. “avg” means average conditional effect. (**F**) Significance of locus measured by −log_10_ *P*. (**G**) Coefficient of slope for locus. (**H**) Distance (bp) between locus peak and closest end of nearest gene (5′ or 3′). (**I**) ENSEMBL gene ID. (**J**) ENSEMBL transcript ID. (**K**) Nearest gene symbol. (**L**) Sense strand. (**M**) Gene start coordinates. (**N**) Gene end coordinates. (**0**) Gene length. (**P**) Longest transcript length. (**Q**) Coding sequence length. (**R**) 5′ untranslated region start coordinates. (**S**) 5′ untranslated region end coordinates. (**T**) 5′ untranslated region length. (**U**) 3′ untranslated region start coordinates. (**V**) 3′ untranslated region end coordinates. (**W**) 3′ untranslated region length. 5′ and 3′ untranslated regions are longest alternatives. (**X**) Exon count. (**Y**) Gene type. (**Z**) Gene description. (**AA** to **AD**) log10p_g_0nM refers to −log_10_ *P* values for growth at 0 nM paclitaxel, etc. (**AE**) log10p_g_avg refers to −log_10_ *P* values for average conditional effect of growth. (**AF**–**AJ**) log10p_d_w1 refers to −log_10_ *P* values for paclitaxel (drug) action at week 1, etc. (**AK**) log10p_d_avg refers to −log_10_ *P* values for average conditional effect of paclitaxel. (**AL**) log10p_g_d_Ix refers to −log_10_ *P* values for interaction between growth time and paclitaxel concentration. (**AM**–**AX**) Coefficients of slope for corresponding −log_10_ *P* values in (AA–AL).

**Table S19.** Non-redundant paclitaxel loci. (**A**) Chromosome. (**B**) Window start position. (**C**) Window stop position. (**D**) Average (midpoint) window position. (**E**) Growth week of paclitaxel locus with maximum −log_10_ *P*. “avg” means average conditional effect. (**F**) Significance of locus measured by −log_10_ *P*. (**G**) Coefficient of slope for locus. (**H**) Distance (bp) between locus peak and closest end of nearest gene (5′ or 3′). (**I**) ENSEMBL gene ID. (**J**) ENSEMBL transcript ID. (**K**) Nearest gene symbol. (**L**) Sense strand. (**M**) Gene start coordinates. (**N**) Gene end coordinates. (**0**) Gene length. (**P**) Longest transcript length. (**Q**) Coding sequence length. (**R**) 5′ untranslated region start coordinates. (**S**) 5′ untranslated region end coordinates. (**T**) 5′ untranslated region length. (**U**) 3′ untranslated region start coordinates. (**V**) 3′ untranslated region end coordinates. (**W**) 3′ untranslated region length. 5*^I^* and 3*^I^* untranslated regions are longest alternatives. (**X**) Exon count. (**Y**) Gene type. (**Z**) Gene description. (**AA** to **AD**) log10p_g_0nM refers to −log_10_ *P* values for growth at 0 nM paclitaxel, etc. (**AE**) log10p_g_avg refers to −log_10_ *P* values for average conditional effect of growth. (**AF**–**AJ**) log10p_d_w1 refers to −log_10_ *P* values for paclitaxel (drug) action at week 1, etc. (**AK**) log10p_d_avg refers to −log_10_ *P* values for average conditional effect of paclitaxel. (**AL**) log10p_g_d_Ix refers to −log_10_ *P* values for interaction between growth time and paclitaxel concentration. (**AM**–**AX**) Coefficients of slope for corresponding −log_10_ *P* values in (AA–AL).

**Table S20.** Interaction loci. (**A**) Chromosome. (**B**) Window start position. (**C**) Window stop position. (**D**) Average (midpoint) window position. (**E**) Type of locus. (**F**) Significance of locus measured by −log_10_ *P*. (**G**) Coefficient of slope for locus. (**H**) Distance (bp) between locus peak and closest end of nearest gene (5′ or 3′). (**I**) ENSEMBL gene ID. (**J**) ENSEMBL transcript ID. (**K**) Nearest gene symbol. (**L**) Sense strand. (**M**) Gene start coordinates. (**N**) Gene end coordinates. (**0**) Gene length. (**P**) Longest transcript length. (**Q**) Coding sequence length. (**R**) 5′ untranslated region start coordinates. (**S**) 5′ untranslated region end coordinates. (**T**) 5′ untranslated region length. (**U**) 3′ untranslated region start coordinates. (**V**) 3′ untranslated region end coordinates. (**W**) 3′ untranslated region length. 5′ and 3′ untranslated regions are longest alternatives. (**X**) Exon count. (**Y**) Gene type. (**Z**) Gene description. (**AA** to **AD**) log10p_g_0nM refers to −log_10_ *P* values for growth at 0 nM paclitaxel, etc. (**AE**) log10p_g_avg refers to −log_10_ *P* values for average conditional effect of growth. (**AF**–**AJ**) log10p_d_w1 refers to −log_10_ *P* values for paclitaxel (drug) action at week 1, etc. (**AK**) log10p_d_avg refers to −log_10_ *P* values for average conditional effect of paclitaxel. (**AL**) log10p_g_d_Ix refers to −log_10_ *P* values for interaction between growth time and paclitaxel concentration. (**AM**–**AX**) Coefficients of slope for corresponding −log_10_ *P* values in (AA–AL).

**Table S21.** Mitochondrial DNA in the RH pools

| Pool | Reads<sup>a</sup> |  |  |  |  | Revert<br>freq | Mito copy number |  |  |
| --- | --- | --- | --- | --- | --- | --- | --- | --- | --- |
|  | Total<sup>b</sup> | Hamster<sup>c</sup> | Human<br>TK1<sup>d</sup> | Hamster<br>mito | Human<br>mito |  | Hamster<br>(align)<sup>e</sup> | Human<br>(align)<sup>f</sup> | Human<br>(TK1)<sup>g</sup> |
| 1 | 42 000 172 | 37 595 178 | 997 | 103 116 | 16 | 0.24 | 800 | 0.12 | 0.74 |
| 2 | 35 358 581 | 30 754 890 | 2636 | 109 494 | 474 | 0.2 | 1040 | 4.42 | 8.67 |
| 3 | 52 166 656 | 45 668 817 | 3302 | 96 305 | 492 | 0.27 | 615 | 3.09 | 6.57 |
| 4 | 32 756 205 | 28 092 611 | 3125 | 61 749 | 45 | 0.29 | 641 | 0.46 | 0.61 |
| 5 | 42 876 176 | 38 167 486 | 2234 | 89 994 | 0 | 0.28 | 688 | 0.00 | 0.00 |
| 6 | 41 449 894 | 37 193 349 | 2573 | 96 898 | 2 | 0.37 | 760 | 0.02 | 0.03 |
| **Mean** | 41 101 281 | 36 245 388 | 2478 | 92 926 | 172 | 0.28 | 757 | 1.35 | 2.77 |
| **s.e.m.** | 2 763 700 | 2 528 698 | 336 | 6797 | 99 | 0.02 | 63 | 0.78 | 1.56 |

<sup>a</sup> All reads except “Total” purged of cross-species alignments.
<sup>b</sup> Identical to same column in **Tables S5** and **S6**.
<sup>c</sup> Identical to “Corrected hamster” column in **Table S6**.
<sup>d</sup> Identical to same column in **Table S7**.
<sup>e</sup> Hamster mitochondrial copy number: $2 \cdot (\text{hamster mitochondrial reads} / [\text{hamster reads} - \text{hamster mitochondrial reads}]) \cdot (\text{hamster genome length} = 2\,368\,906\,908 \text{ bp} / \text{hamster mitochondrial genome length} = 16\,283 \text{ bp})$ .
<sup>f</sup> Human mitochondrial copy number deduced from hamster genome alignments: $2 \cdot (\text{human mitochondrial reads} / [\text{hamster reads} - \text{hamster mitochondrial reads}]) \cdot (\text{hamster genome length} = 2\,368\,906\,908 \text{ bp} / \text{human mitochondrial genome length} = 16\,569 \text{ bp})$ .
<sup>g</sup> Human mitochondrial copy number deduced from TK1 alignments: $1 \times (\text{human mitochondrial reads} / \text{TK1 reads}) \cdot (\text{TK1 window length} = 1 \times 10^6 \text{ bp} / \text{human mitochondrial genome length} = 16\,569 \text{ bp}) \cdot (1 - \text{reversion frequency})$ .

**Table S22.** Significance values in human genome for growth, paclitaxel, interaction and hamster mitochondrial loci. (**A**) Chromosome. (**B**) Window start position. (**C**) Window stop position. (**D**) Average (midpoint) window position. (**E**–**H**) log10p_g_0nM refers to −log_10_ *P* values for growth at 0 nM paclitaxel, etc. (**I**) log10p_g_avg refers to −log_10_ *P* values for average conditional effect of growth. (**J**–**N**) log10p_d_w1 refers to −log_10_ *P* values for paclitaxel (drug) action at week 1, etc. (**O**) log10p_d_avg refers to −log_10_ *P* values for average conditional effect of paclitaxel. (**P**) log10p_g_d_Ix refers to −log_10_ *P* values for interaction between growth time and paclitaxel concentration. (**Q**) log10p_hamster_mito refers to −log_10_ *P* values for regulation of hamster mitochondrial copy number by human genome. (**R**– **AD**) Coefficients of slope for corresponding −log_10_ *P* values in (E–Q). Windows of 1 Mb, 10 kb steps.

**Table S23.** Significance thresholds in human genome for growth, paclitaxel, interaction and hamster mitochondrial loci. (**A**) Genome scans. log10p_g_0nM refers to −log_10_ *P* values for growth at 0 nM paclitaxel, etc., log10p_g_avg refers to −log_10_ *P* values for average conditional effect of growth, log10p_d_w1 refers to −log_10_ *P* values for paclitaxel (drug) action at week 1, etc., log10p_d_avg refers to −log_10_ *P* values for average conditional effect of paclitaxel, log10p_g_d_Ix refers to −log_10_ *P* values for interaction between growth time and paclitaxel concentration. log10p_hamster_mito refers to −log_10_ *P* values for regulation of hamster mitochondrial copy number by human genome. (**B**) Significance thresholds at 5% family-wise error rate (FWER) obtained using permutation.

**Table S24.** Significance values in human genome for growth, paclitaxel, interaction and human mitochondrial loci. (**A**) Chromosome. (**B**) Window start position. (**C**) Window stop position. (**D**) Average (midpoint) window position. (**E**–**H**) log10p_g_0nM refers to −log_10_ *P* values for growth at 0 nM paclitaxel, etc. (**I**) log10p_g_avg refers to −log_10_ *P* values for average conditional effect of growth. (**J**–**N**) log10p_d_w1 refers to −log_10_ *P* values for paclitaxel (drug) action at week 1, etc. (**O**) log10p_d_avg refers to −log_10_ *P* values for average conditional effect of paclitaxel. (**P**) log10p_g_d_Ix refers to −log_10_ *P* values for interaction between growth time and paclitaxel concentration. (**Q**) log10p_human_mito refers to −log_10_ *P* values for regulation of human mitochondrial copy number by human genome. (**R**– **AD**) Coefficients of slope for corresponding −log_10_ *P* values in (E–Q). Windows of 1 Mb, 10 kb steps.

**Table S25.** Significance thresholds in human genome for growth, paclitaxel, interaction and human mitochondrial loci. (**A**) Genome scans. log10p_g_0nM refers to −log_10_ *P* values for growth at 0 nM paclitaxel, etc., log10p_g_avg refers to −log_10_ *P* values for average conditional effect of growth, log10p_d_w1 refers to −log_10_ *P* values for paclitaxel (drug) action at week 1, etc., log10p_d_avg refers to −log_10_ *P* values for average conditional effect of paclitaxel, log10p_g_d_Ix refers to −log_10_ *P* values for interaction between growth time and paclitaxel concentration. log10p_human_mito refers to −log_10_ *P* values for regulation of human mitochondrial copy number by human genome. (**B**) Significance thresholds at 5% family-wise error rate (FWER) obtained using permutation.

**Table S26.** Significance values in hamster genome for growth, paclitaxel and interaction loci. (**A**) Contig ID. (**B**) Chromosome. (**C**) Window start position. (**D**) Window stop position. (**E**) Average (midpoint) window position. (**F**–**I**) log10p_g_0nM refers to −log_10_ *P* values for growth at 0 nM paclitaxel, etc. (**J**) log10p_g_avg refers to −log_10_ *P* values for average conditional effect of growth. (**K**–**O**) log10p_d_w1 refers to −log_10_ *P* values for paclitaxel (drug) action at week 1, etc. (**P**) log10p_d_avg refers to −log_10_ *P* values for average conditional effect of paclitaxel. (**Q**) log10p_g_d_Ix refers to −log_10_ *P* values for interaction between growth time and paclitaxel concentration. (**R**–**AC**) Coefficients of slope for corresponding −log_10_ *P* values in (F–Q). Windows of 1 Mb, 10 kb steps.

**Table S27.** Significance thresholds in hamster genome for growth, paclitaxel and interaction loci. (**A**) Genome scans. log10p_g_0nM refers to −log_10_ *P* values for growth at 0 nM paclitaxel, etc., log10p_g_avg refers to −log_10_ *P* values for average conditional effect of growth, log10p_d_w1 refers to −log_10_ *P* values for paclitaxel (drug) action at week 1, etc., log10p_d_avg refers to −log_10_ *P* values for average conditional effect of paclitaxel, log10p_g_d_Ix refers to −log_10_ *P* values for interaction between growth time and paclitaxel concentration. (**B**) Significance thresholds at 5% family-wise error rate (FWER) obtained using permutation.

**Table S28.**
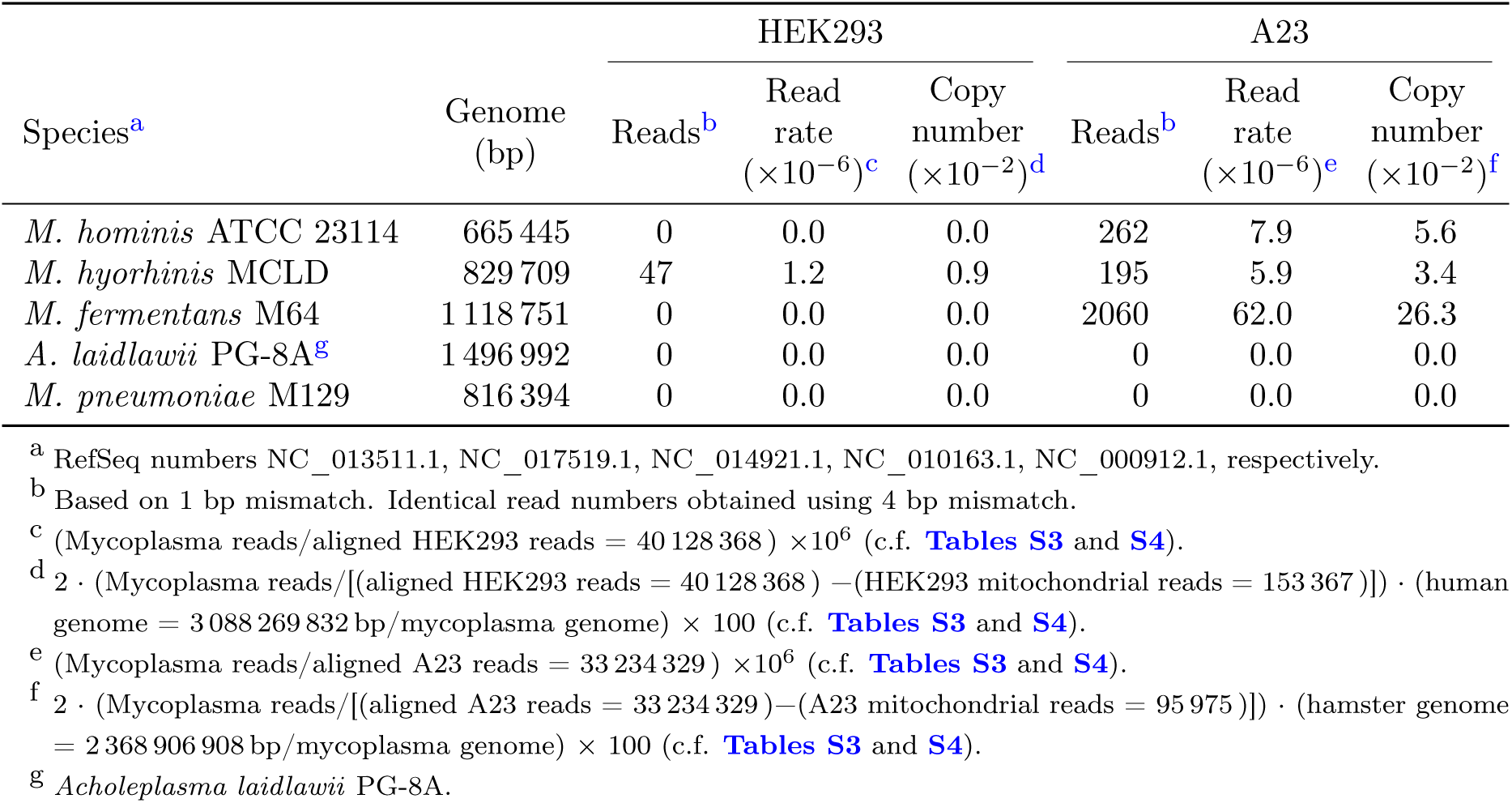
Mycoplasma contamination

**Table S29.** *M. fermentans* reads in RH samples. (**A**) RH sample ID. RH1_d0_w0 refers to RH pool 1 at 0 nM paclitaxel (drug) and 0 weeks of growth, etc. (**B**) Pool. (**C**) Paclitaxel concentration (nM). (**D**) Weeks of growth. (**E**) Paclitaxel history nested inside pool. (**F**) *M. fermentans* reads.

**Table S30.** Significance values in human genome for growth, paclitaxel, interaction and *M. fermentans* loci. (**A**) Chromosome. (**B**) Window start position. (**C**) Window stop position. (**D**) Average (midpoint) window position. (**E**–**H**) log10p_g_0nM refers to −log_10_ *P* values for growth at 0 nM paclitaxel, etc. (**I**) log10p_g_avg refers to −log_10_ *P* values for average conditional effect of growth. (**J**–**N**) log10p_d_w1 refers to −log_10_ *P* values for paclitaxel (drug) action at week 1, etc. (**O**) log10p_d_avg refers to −log_10_ *P* values for average conditional effect of paclitaxel. (**P**) log10p_g_d_Ix refers to −log_10_ *P* values for interaction between growth time and paclitaxel concentration. (**Q**) log10p_myco_ferm refers to −log_10_ *P* values for regulation of *M. fermentans* copy number by human genome. (**R**–**AD**) Coefficients of slope for corresponding −log_10_ *P* values in (E–Q). Windows of 1 Mb, 10 kb steps.

**Table S31.** Significance thresholds in human genome for growth, paclitaxel, interaction and *M. fermentans* loci. (**A**) Genome scans. log10p_g_0nM refers to −log_10_ *P* values for growth at 0 nM paclitaxel, etc., log10p_g_avg refers to −log_10_ *P* values for average conditional effect of growth, log10p_d_w1 refers to −log_10_ *P* values for paclitaxel (drug) action at week 1, etc., log10p_d_avg refers to −log_10_ *P* values for average conditional effect of paclitaxel, log10p_g_d_Ix refers to −log_10_ *P* values for interaction between growth time and paclitaxel concentration. log10p_myco_ferm refers to −log_10_ *P* values for regulation of *M. fermentans* copy number by human genome. (**B**) Significance thresholds at 5% family-wise error rate (FWER) obtained using permutation.

